# A Glimpse Into the Cenomanian: Palynology of the Arlington Archosaur Site, Late Cretaceous Western Interior Seaway, Texas, USA

**DOI:** 10.1101/2023.12.04.569281

**Authors:** Maria Antonieta Lorente, Christopher Noto, Peter Flaig

## Abstract

The Arlington Archosaur Site (AAS) between Dallas and Fort Worth, Texas, is known as a rich fossiliferous section. The age of these rocks is generally considered to be mid-Cenomanian, but conflicting evidence suggests the age may be as young as the late Cenomanian–early Turonian. To address the issue, a palynological study was designed and conducted based on the close sampling of the lithofacies. Palynological samples were processed according to the standard acid preparation. The study was quantitative and focused on associations to determine the paleoenvironment, paleoclimate, biostratigraphy, and age of exposure. The rich palynological assemblages comprise spores from seedless plants, gymnosperms, angiosperms, fungi, algae, and dinoflagellate cysts. Bryophytes were abundant mainly in Facies A and B, with *Zlivisporis cenomanianus* taking over the bryophytes’ habitat in Facies D. Lycophytes abundant in the alluvial and coastal plains are considered to have been transported. Conifers were the predominant group of gymnosperms, also mainly transported into the section. Freshwater algal remains include *Schizophacus laevigatus*/*Ovoidites parvus*, *Schizosporis reticulatus*, *Botryococcus* sp., and *Pediastrum* sp. Acanthomorph acritarchs present in low abundance and diversity appear following shallow marine dinoflagellates’ spikes and before freshwater colonial algal spikes. The vegetation signal at Noto’s Facies A and B indicates tropical to subtropical shallow marine to coastal plains, while Noto’s Facies D indicates tidally influenced areas. Also, picks of the diversity and abundance of dinoflagellate cysts are interpreted as an increased marine influence and proposed as possible flooding surfaces. The results support the alternation of marine incursions within deltaic and floodplain sequences, related to regional climate oscillation that affected the vegetation on the upland drainage area.

Key palynological markers point to an early Late Cenomanian age, and the presence of the *Cyclonephelium compactum–C. membraniphorum* (Ccm morphological plexus) signals that the incursion of boreal waters during the Plenus Cold Event of the Ocean Anoxic Event 2 may have reached as far south as the AAS area. This coincides with vegetation trends that suggest a cooler and less humid climate at the start of Facies A, where Ccm is more abundant.

## INTRODUCTION

The Arlington Archosaur Site (AAS, Figure 1) is between Dallas and Fort Worth, Texas. The AAS was discovered independently in 2003 by local fossil collector Art Sahlstein and University of Texas at Arlington students Phil Kirchhoff and Bill Walker. Since then, the site has undergone multiple detailed surveys and samplings, most recently in 2017. The site is rich in well-preserved dinosaurs, crocodiles, turtles, amphibians, fish, invertebrates, and plant remains. Based on reports of ammonite species *Conlinoceras tarrentense* found elsewhere in Bear Creek, near Dallas–Fort Worth, the age of the rocks at the Arlington Archosaur Site is generally accepted as mid-Cenomanian. Rocks exposed at the site have been identified as belonging to the Lewisville Formation. However, conflicting stratigraphic, lithostratigraphic, and biostratigraphic evidence for the Lewisville Formation suggests some parts may be as young as the late Cenomanian/early Turonian (Ambrose et al., 2009; Christopher, 1982; Gifford, 2021; Jacobs et al., 2005; Kennedy & Cobban, 1990).

**Figure 1.**
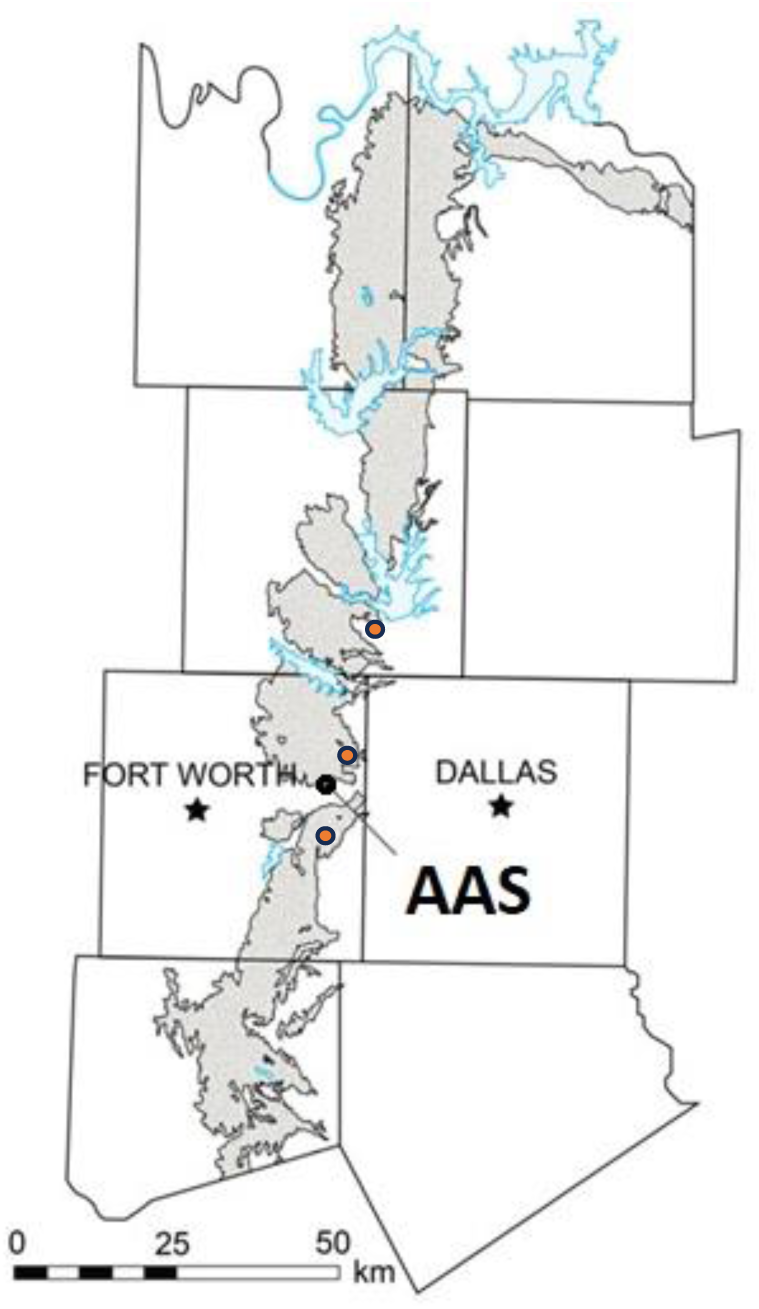
Surface exposures of the Woodbine Group (gray) in north-central Texas, with the Arlington Archosaur Site (AAS) location (black dot) and the three locations (red dots) of the palynology study by Cloos (2018). Base map of the Upper Cretaceous Outcrop Belt created by Thomas Adams.

Despite a relatively long history of paleontological research, the Woodbine Group has produced a limited number of palynological studies. Hedlund (1966) was the first to document fossil pollen assemblages from members of the Red Branch in Oklahoma, which is stratigraphically close to the base of the Lewisville Formation (Bergquist, 1949; Denne et al., 2016; Stephenson, 1952). He identified 75 palynomorphs, divided into five assemblages dominated by ferns and angiosperms, and interpreted them as growing in humid tropical climates (Hedlund, 1966). This work was recently followed up by Cloos (2018), who sampled three separate Woodbine Group sites along a north-south transect in the Upper Cretaceous Outcrop Belt of the East Texas Basin. She found significant overlap in palynotypes between the sites, particularly among Cupressaceae, cycads, ferns, and some angiosperms that live in hot, humid, subtropical to tropical climates (Cloos, 2018).

Most significant is the first documented detection of tree-pore pollen (*Pseudoplicapollis* sp.) of Normapolles affinity. The genus *Pseudoplicapollis* first appeared in the early to mid- Turonian (Christopher, 1979), suggesting that the age of this region was much younger than currently generally believed.

Some studies of the stratigraphy of the Woodbine and Eagle Ford Groups place the Tarrant Member of the Eagle Ford Group within the Lewisville Formation of the Woodbine Group, where it is interpreted to be coeval with or overlying the Arlington Member (Denne et al., 2016; Stephenson, 1952). This placement is based in part on similarities in their palynological assemblages (Brown & Pierce, 1962; Christopher, 1982), but many of the palynotypes identified are long-lived taxa of little biostratigraphic value.

It is a widespread practice to use the lithostratigraphic, biostratigraphic, and chronostratigraphic units as interchangeable and equivalent terms for certain stratigraphic intervals. This practice can lead to a disruption in the understanding of the stratigraphic architecture of an area. For example, when the Lewisville Formation is identified at the surface (or subsurface), its age is assumed to be Middle Cenomanian. However, changes in lateral facies may produce similar lithologies at different times. For example, Denne et al. (2016, p. 39, Fig. 23) showed “Lewisville”-like phases below and above the Middle Cenomanian discontinuity. In addition, there are also sporadic occurrences within the Upper Cenomanian of similar facies, although mainly towards the Red River area. Using lithology (facies) identification as age evidence has led to controversy and apparent inconsistencies in defining the chronostratigraphic position of specific outcrops.

Considering that the AAS outcrop is a few meters thick, it is difficult to say to which of the “Lewisville facies” occurring at different times it belongs.

To address this issue and separate overlapping contributions of tectonics, climate, and evolutionary changes, during the 2016 fieldwork campaign, Noto specifically sampled the outcrop for palynological studies.

The study’s objectives included (1) helping to elucidate the true chronostratigraphic position of rocks exposed at the AAS and (2) determining the paleoenvironments and possible climate changes recorded at the site.

## MATERIAL AND METHODS

### Studied Section and Sampling

The stratigraphy, sedimentology, and depositional environments present at the AAS have been described in detail by Noto et al. (2015, 2023) and Adams et al. (2017). Noto described four facies based on lithology at the AAS named using consecutive letters: Facies A, B, C, and D.

The surface outcrop consists of a hillside approximately 200 m long, with the largest fossil concentrations within a main quarry about 50 m long. Vertical exposure varies from 1 m to 2.5 m due to surface erosion and 5° east–west dip. A 70-m-long outcrop section was chosen for palynological sampling, including where most fossils have been recovered. The location of each profile was chosen to maximize either the vertical extent of facies exposure and/or the clarity of the borders between facies. The relative position of each section is marked according to a meter-square grid system used for recording the location of fossils uncovered in the quarry, which is anchored by a permanent, georeferenced origin stake. GPS coordinates for each section were also recorded using a handheld Garmin unit. A total of 31 samples of 15–20 g each were collected in 2016 from fresh surface exposures at different heights measured from the base of the exposure when visible.

The location of each section and each sample and its corresponding position in relation to Noto’s facies are shown in Figure 2.

**Figure 2.**
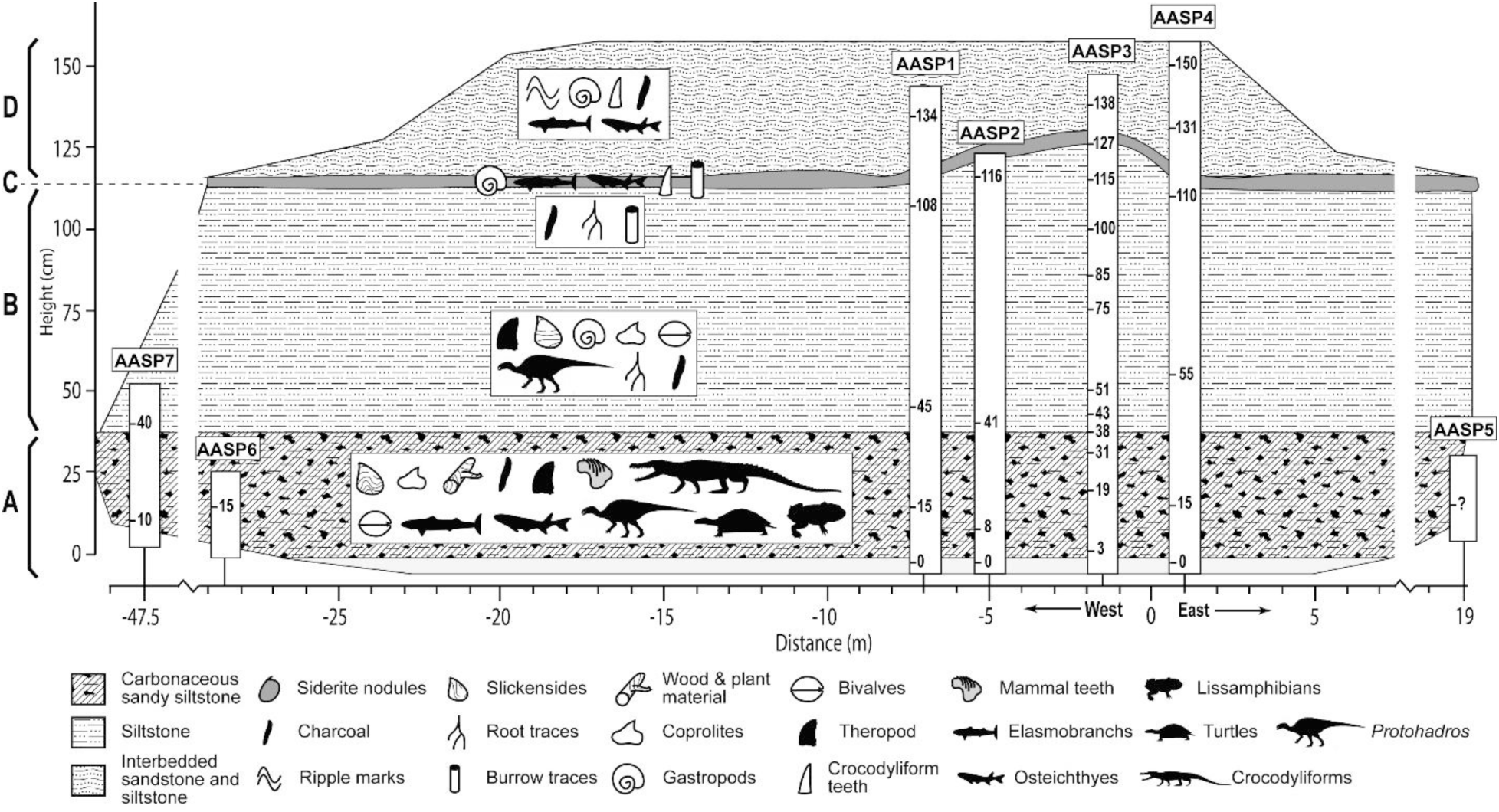
Location of the samples Christopher Noto took for palynological analyses during the 2016 field campaign at the AAS.

### Palynological Sample Preparation

There are different techniques for palynological sample preparation. Especially in the last years, much effort has been placed into developing nonacid techniques with a lower risk for humans and the environment. Extensive revisions of palynological sample preparation and the effects on the palynological assemblage recovered have shown variations in abundance and preservation of the recovered palynological assemblages (e.g., Lorente, 1986; O’Keefe & Eble, 2012; Pound et al., 2021; Riding, 2022). We used conventional acid preparation techniques for this study because the samples were hardened, and most publications on samples from similar time intervals have been prepared using acid techniques. Thus, we ensure that the assemblages recovered from the AAS sections are comparable to assemblages from the same stratigraphic interval.

Samples were prepared following standard palynological techniques involving hydrochloric acid (10% HCl) and hydrofluoric acid (70% HF) maceration to dissolve carbonate and siliceous contents. Then, the organic fraction was separated using centrifugation in a heavy liquid (ZnBr2, ρ 2.0). A Schultz solution was applied before sieving the final organic concentrate with a 10-μm mesh for light oxidation of the organic concentrate. A drop of the final sieved palynological residue was pipetted off, and one drop of polyvinyl alcohol was mixed in with a glass stirring rod. Once the polyvinyl alcohol/residue had dried, one drop of clear casting resin was added to fix the coverslip.

### Palynological Sample Analysis

All samples were studied with a Leitz Dialux microscope equipped with Leitz NPL fluotar objectives (10X, 25X, 40X, and 100X) and 10X WF oculars. Species identification was done at 400X and counting at 250X and 100X under white transmitted light. The following counting technique was routinely used to obtain quantitative data for abundance analysis of individual species. All specimens in 150 fields of view (FOV) at 250X were counted for each slide, and a subsequent screening at 100X of the entire slide (22 x 40 mm) was conducted to locate scarce species. All palynological data was transferred to Excel files and exported to Tilia (Grimm, 1991) for further display and analysis of results. Due to the proximity and overall similarity, the data from sections AASP1, AASP2, AASP3, and AASP4 were combined into two composite sections, named AASP1–2 and AASP 3–4, and for final interpretation, both were combined into one section, AASP 1–2-3-4. These labels will be used for the remainder of the paper when referring to the data from these sections. Samples for each section are referenced in the palynological diagrams using their measured height from the basal contact.

## RESULTS

This study evaluates the palynological associations obtained from outcrops at the Arlington Archosaur Site (AAS) to determine the biostratigraphy, paleoenvironment, paleoclimate, and age of exposure to achieve the highest possible resolution of these aspects. Due to the tight sampling, the resolution of this palynological work is very high.

We identified as many terrestrial pollens, spores, and dinoflagellate cysts as possible. Based on the presence of dinoflagellate cysts, we support marine incursions within deltaic and floodplain sequences, which represent high-frequency marine flooding surfaces.

### AASP 3–4

The assemblage observed in the combined section AASP 3–4 comprises a rich terrestrial sporomorph component and a less diversified dinoflagellate cysts assemblage.

The terrestrial sporomorph assemblage is rich. Over 80 different species were identified in total (Figure 3, Plate I), with diversity per individual sample reaching between 16 and 35 species (Figure 4).

**Figure 3.**
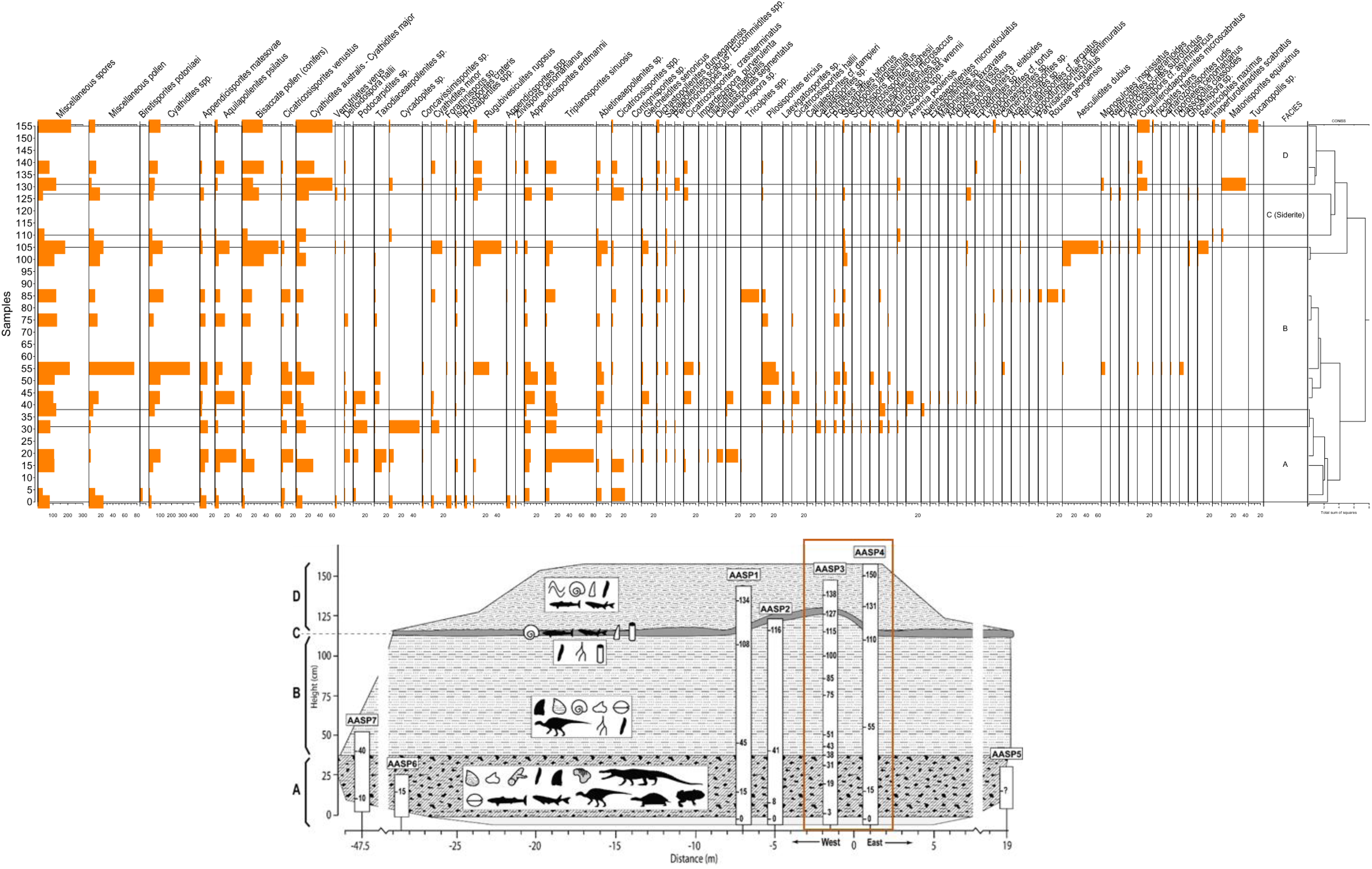
Combined section AASP 3-4 pollen and spores taxa abundance diagram with stratigraphically constrained cluster analysis (total sum of squares).

**Figure 4.**
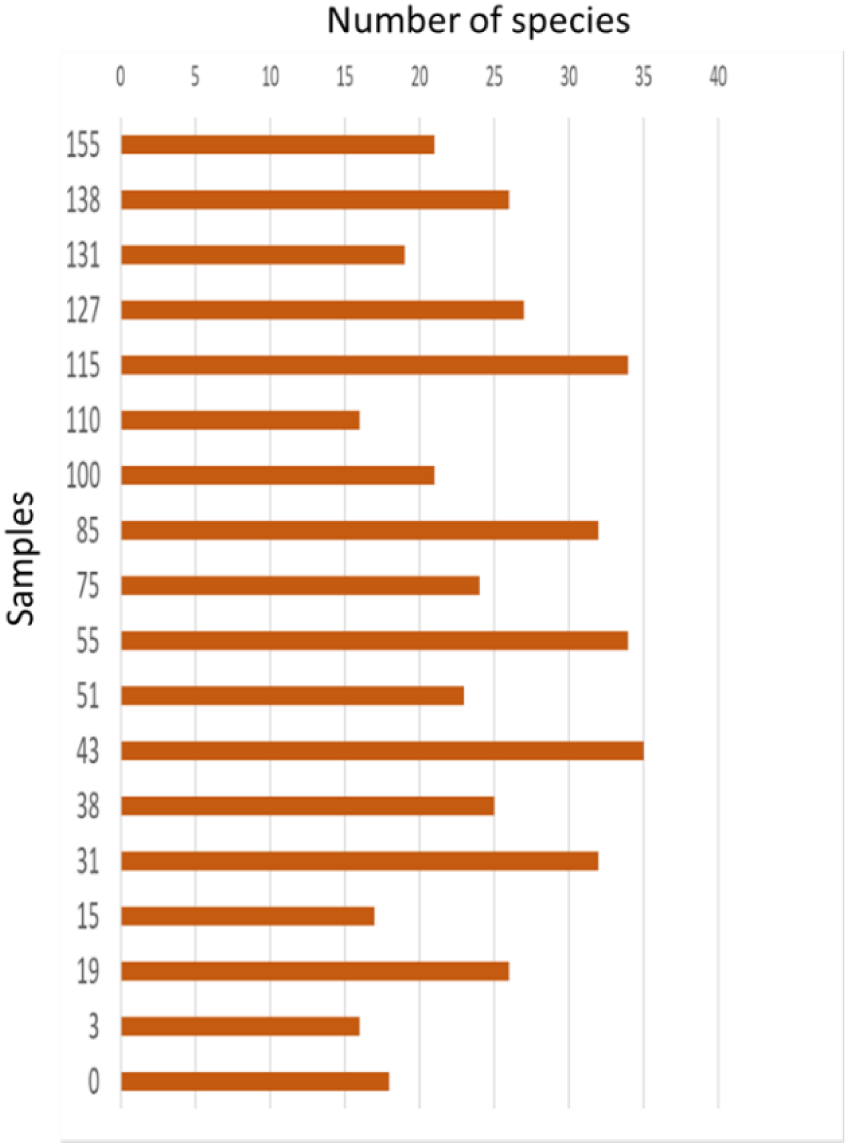
AASP 3–4 sporomorph species diversity histogram.

The assemblages are rich in spores, predominantly from pteridophytes, and pollen from gymnosperms, mostly conifers (Figure 5) and angiosperms. Some samples show an increase in the presence of angiosperms. There is an apparent increase in dicotyledon angiosperms from Noto’s Facies B upwards.

**Figure 5.**
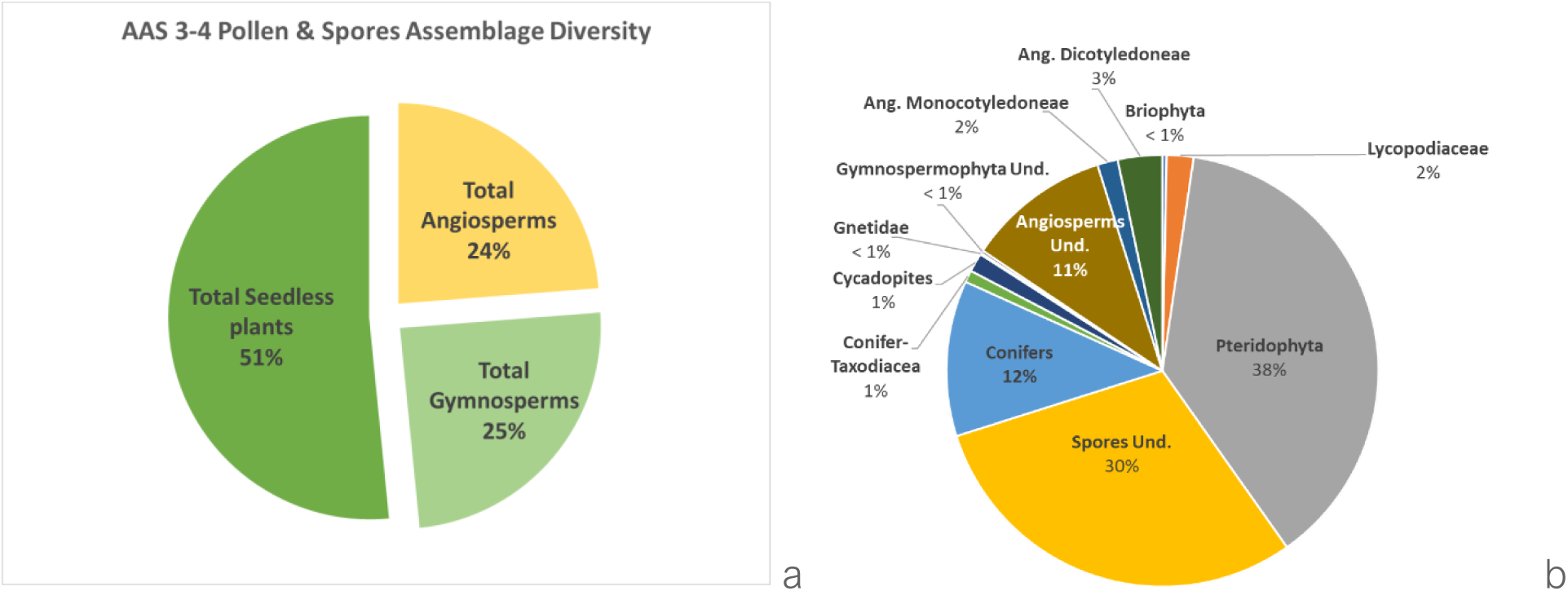
(a) The average composition of the AASP 3–4 sporomorph assemblage. Most sporomorphs belong to the spores of seedless plants, while the pollen from gymnosperms and angiosperms is present in the section in about the same proportion. (b) Percentage composition of the average pollen and spore assemblage in AASP 3–4 by major sporomorphs associated with plants’ taxa contribution.

Dinoflagellate cysts are present throughout the section but in lower quantities than sporomorphs. Twenty-four species were identified (Figure 6, Plate II), with a diversity from one to 14 species per sample (Figure 7).

**Figure 6.**
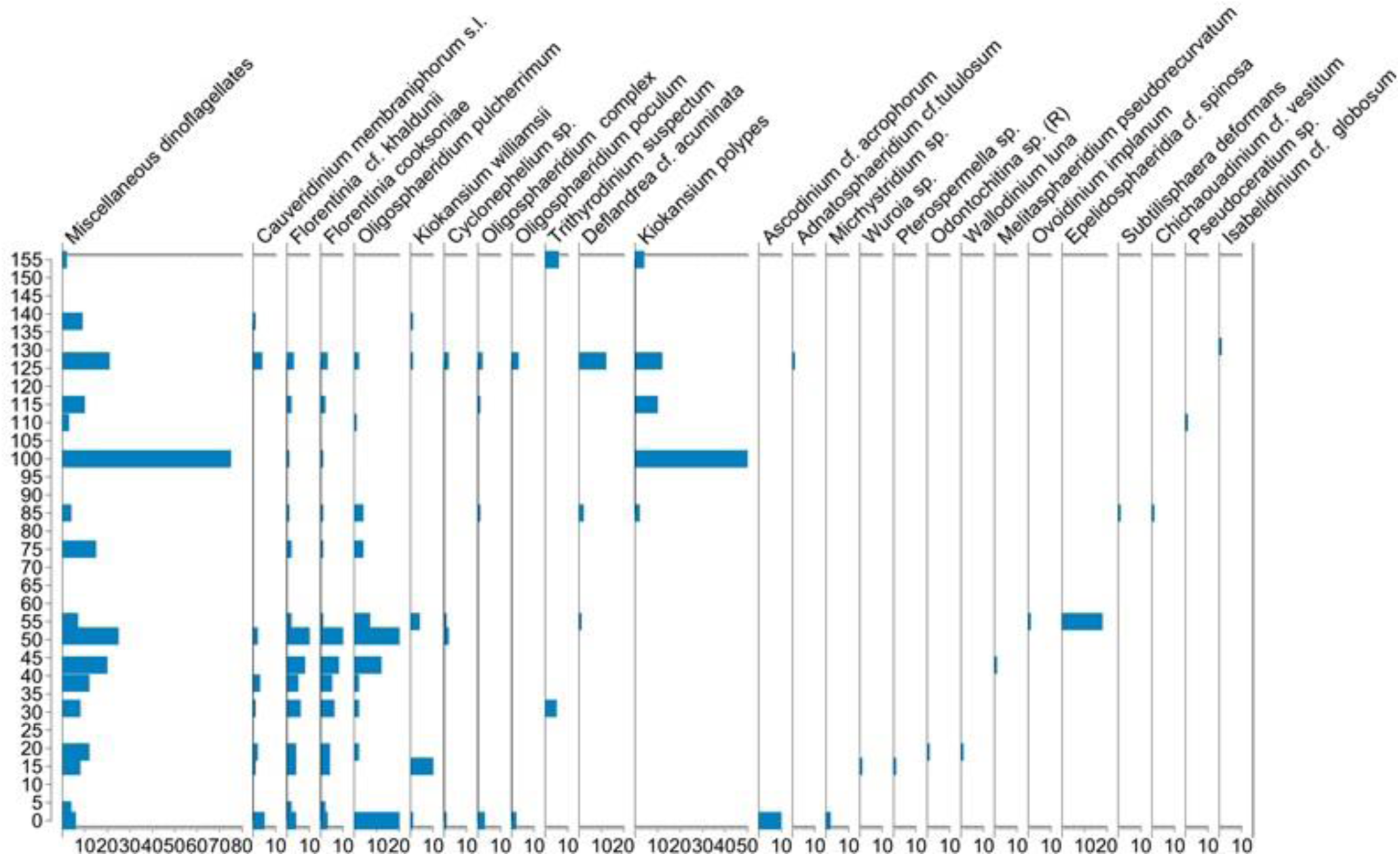
The dinoflagellate cysts’ species abundance diagram shows all species’ occurrences in AASP 3–4.

**Figure 7.**
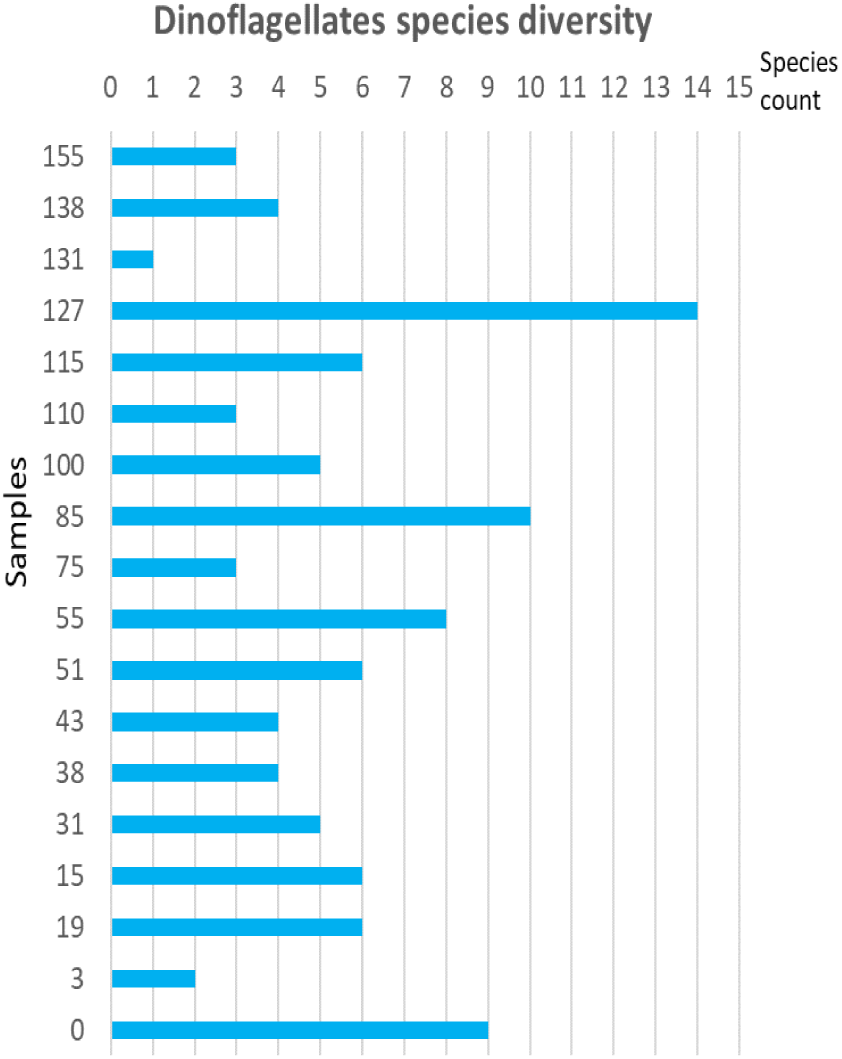
AASP 3–4 dinoflagellate cysts’ species diversity histogram per sample.

Some of the characteristics observed in the assemblages in the combined section AASP 3– 4 are as follows:

- The assemblage’s richness (abundance and diversity) along the section is highly variable. Still, more than half of the species in each sample are from ferns or other seedless plants (Figure 5b).

- Pteridophytes (mostly ferns) are the most varied single component of the assemblage throughout the section. Spores from other botanical groups, e.g., lycophytes and bryophytes, are present but are scarce (Figure 5b).

- *Cupuliferoidaepollenites microscabratus*, an angiosperm, is present in the uppermost part of Noto’s Facies B and throughout Facies D.

### AASP 1–2

The assemblage observed in the combined section AASP 1–2 comprises a rich terrestrial sporomorph component and a less diversified dinoflagellate cysts assemblage, similar to those observed in sections AASP 3–4.

The abundance of specimens per major palynomorph group is variable (Figure 8), with the lowermost part of Noto’s Facies A generally having the most abundant assemblages, although the abundance strongly decreases toward the top of the lithofacies. Noto’s Facies B is moderately rich, and Noto’s Facies D shows a strong increase in sporomorphs (terrestrial) and a decrease in the abundance of dinoflagellate cysts (marine) palynomorphs.

**Figure 8.**
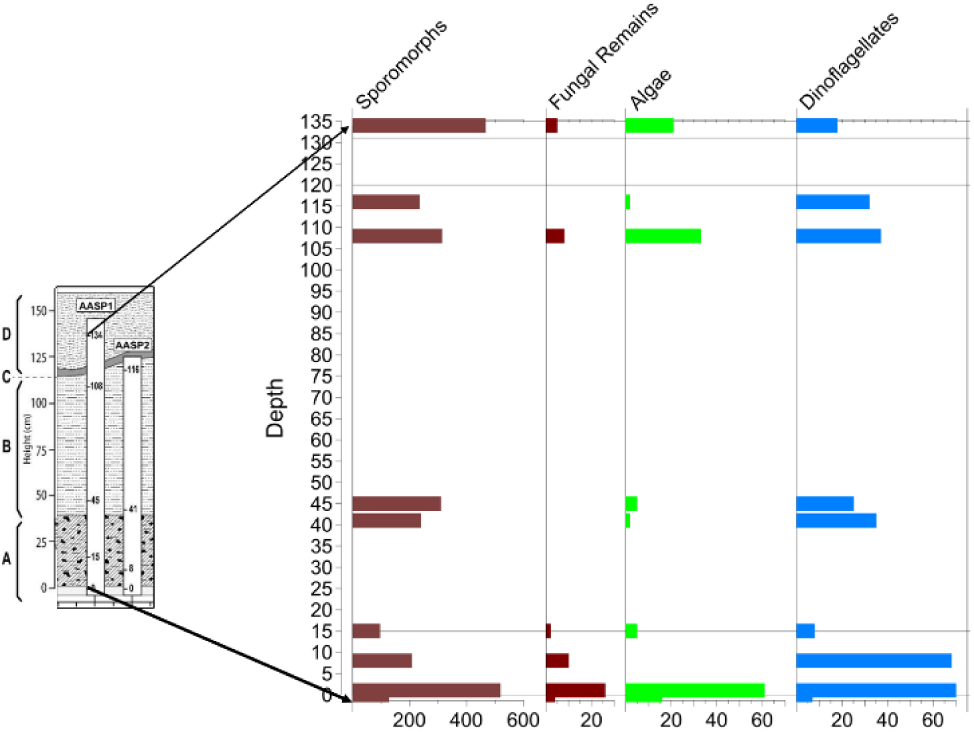
Abundance of major palynomorph groups, section AASP 1-2.

The terrestrial sporomorph assemblage is varied, with 38 spore species and 42 pollen species identified—in total, over 80 different species (Figure 9). The spores and pollen assemblage is dominated by species of the Cyatheaceae (*Cyathidites* spp.), a family of tropical ferns that includes tree ferns. Other important components of the assemblage are conifers. The pollen from these gymnosperms is also very abundant. The pollen of *Cupuliferoidaepollenites microscabratus*, an angiosperm, is abundant at the base of Noto’s Facies A but not in the other facies. Cluster analysis of the sporomorph assemblage shows two major clusters: one that includes the assemblages in the lower part, which includes samples in Noto’s Facies A and the lowermost part of Noto’s Facies B, and a second cluster that includes the samples taken in the uppermost part of Noto’s Facies B and Facies D.

**Figure 9.**
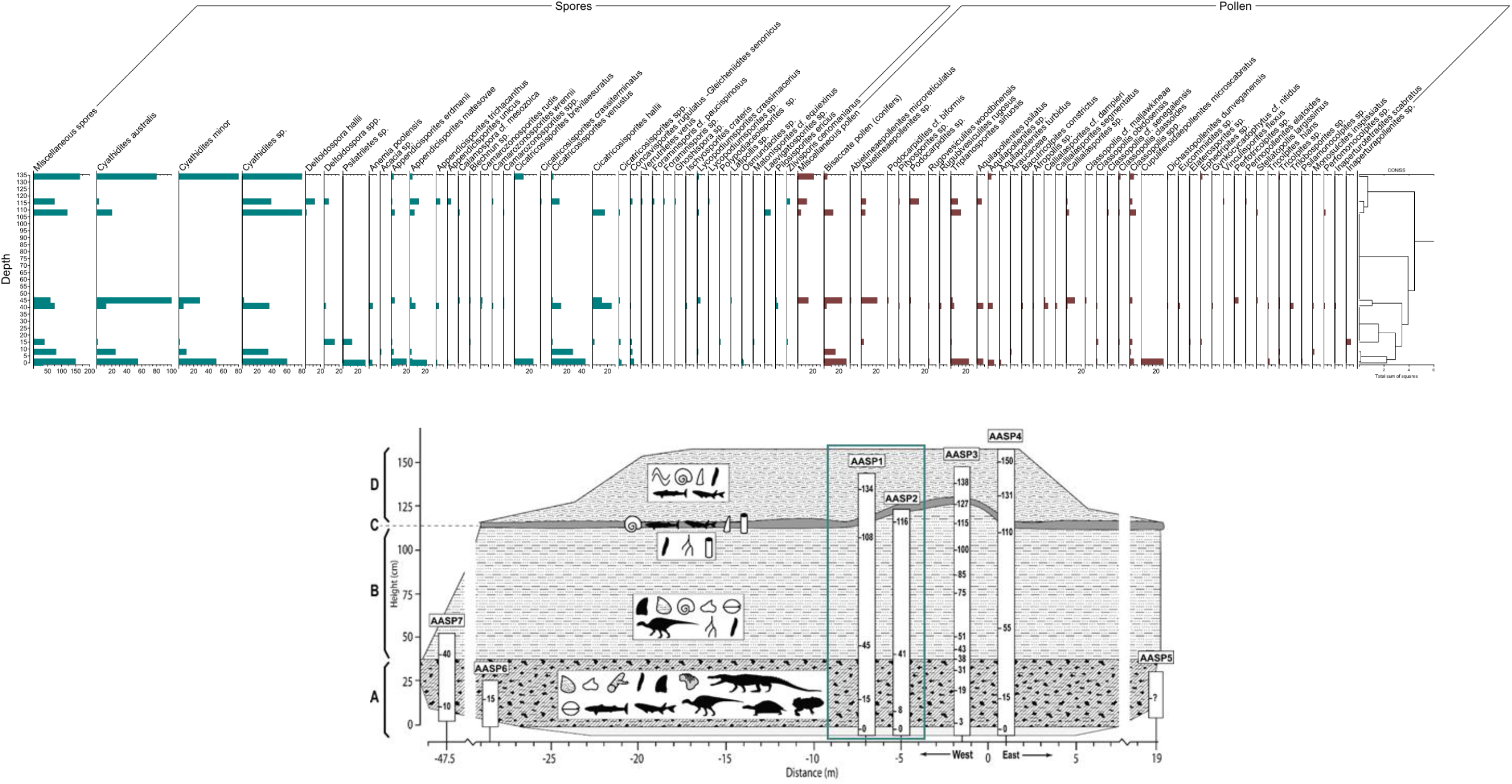
Combined section AASP 1–2. Pollen and spores’ abundance diagram with stratigraphically constrained cluster analysis (CONISS, total sum of squares).

Dinoflagellate cysts are present throughout the section, but in lower quantities than sporomorphs. The same happens with freshwater algae and fungal remains. Twenty-eight dinoflagellate cyst species were identified, along with nine types of freshwater algae and different types of fungal remains (Figure 10).

**Figure 10.**
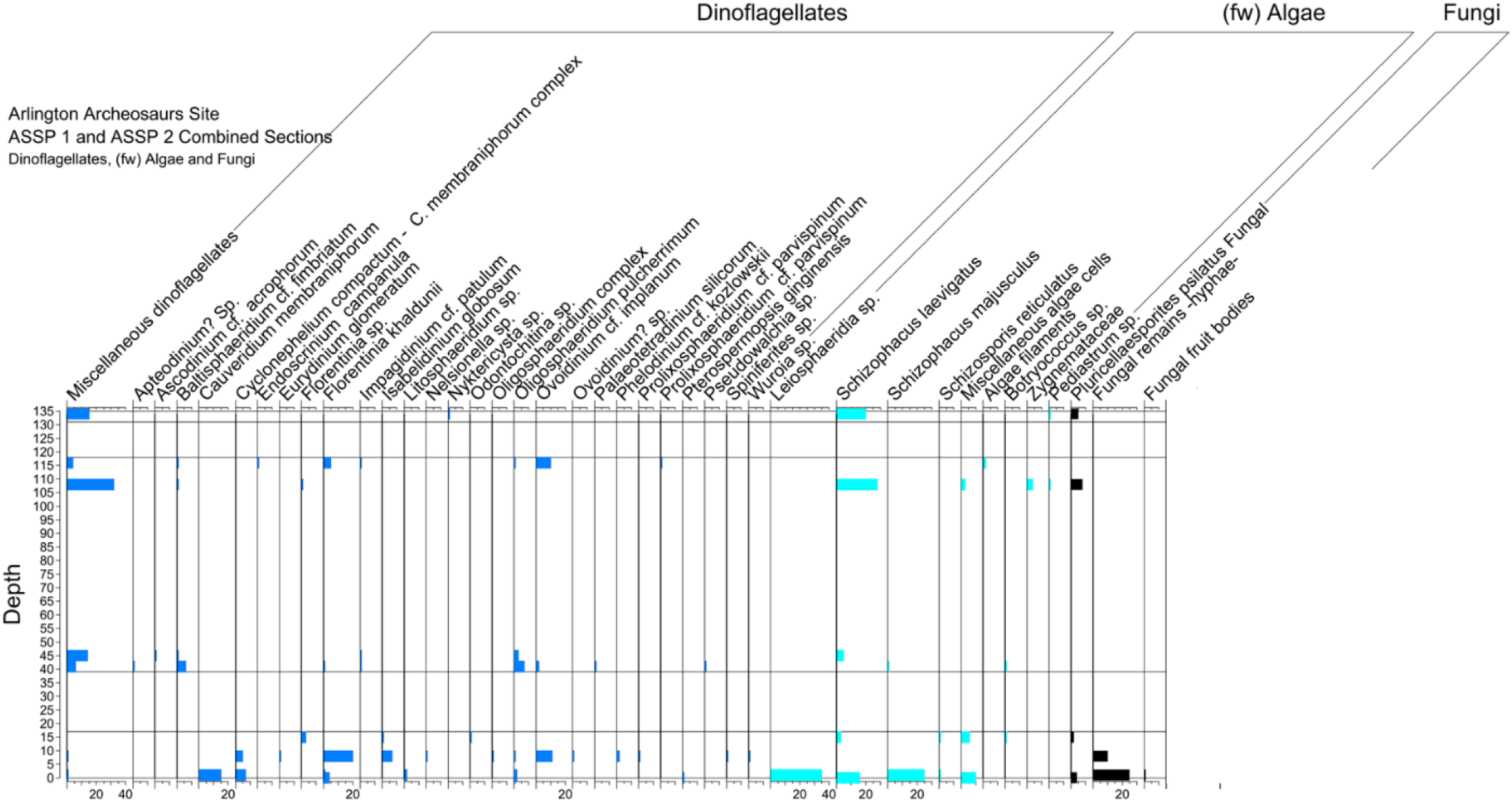
Combined section AASP 1–2. The dinoflagellate cysts, freshwater algae, and fungi types diagram shows the diversity and abundance of the assemblage. Includes all species, even those of rare occurrence.

The following are some characteristics observed in the assemblages in the combined section AASP 1–2:

- The sporomorph assemblage’s richness (abundance and diversity) along the section is highly variable. Still, more than half of the species in each sample are from ferns or other seedless plants (Figure 9).
- Pteridophytes (mostly ferns) are the most varied single component of the assemblage throughout the section. Spores from other botanical groups, e.g., lycophytes and bryophytes, are present but scarce.
- *Cupuliferoidaepollenites microscabratus* is present only in Noto’s Facies A.
- The general assemblage’s composition and abundance is similar to the one observed in AASP 3–4.

### AASP 6–7 and AASP 5

Sections AASP 6–7 and AASP 5 are restricted to Noto’s Facies A.

The assemblage observed in the combined section AASP 6–7 is less rich in palynomorph species than those in combined sections AASP 1–2 and AASP 3–4. This is because AASP 6–7 and AASP 5 are mainly restricted to Noto’s Facies A and the lowermost part of Facies B. *Cupuliferoidaepollenites microscabratus*, an angiosperm, is absent in both sections, as expected from the distribution of the species seen in the combined section AASP 3–4. The distribution chart for Noto’s AASP 6–7 is in Figure 11.

**Figure 11.**
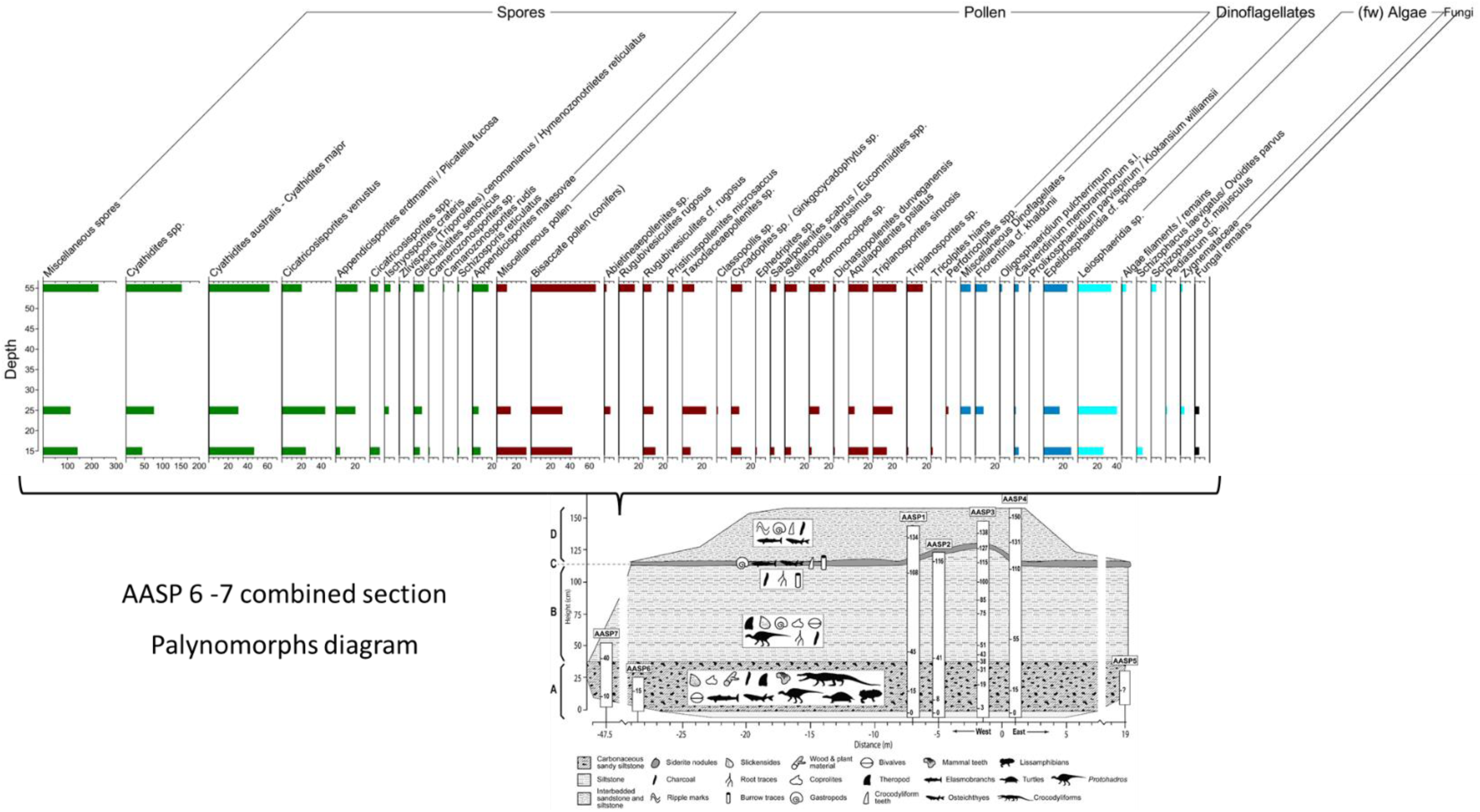
AASP 6–7. Palynomorph distribution chart showing the diversity and richness of the major palynomorph groups.

The total number of spore species is nine, with the Cyatheaceae (tree ferns) being largely dominant, including *Cyathidites* spp. and *Cyathidites australis*–*Cyathidites major*. Other abundant pteridophytes’ spores are species of *Cicatricosisporites* sp., *Appendicisporites matesovae*, and *Concavisporites rugulatus*–*Gleicheiidites senonicus*.

There are 17 species of pollen, the assemblage being dominated by the pollen of conifers, which is similar to the assemblages in sections AASP 1–2 and AASP 3–4. Gymnosperms are dominated by the pollen of conifers (*Rugubivesiculites* cf. *rugosus*), *Cycadopites* sp. / *Ginkgocycadophytus* sp. and *Taxodiaceaepollenites* sp.

The angiosperms are represented by monocotyledons (palm types) as *Perfomonocolpites* sp. Other dubious angiosperms are *Triplanosporites sinuosis*, *Aquilapollenites psilatus*, and miscellaneous pollen types.

In AASP 6–7, *Leiosphaeridia* sp. is very abundant. as well as dinoflagellate cysts such as *Epelidosphaeridia* cf. *spinosa*, and in fewer amounts, *Cauveridinium membraniphorum* complex, *Florentinia* cf. *khaldunii*, and other miscellaneous dinoflagellate cysts.

In AASP 5, the sporomorph group is represented by *Taxodiaceaepollenites* sp., *Cyathidites* spp., *Cyathidites australis*–*Cyathidites major*, and bisaccate pollen (conifers).

*Leiosphaeridia* sp. is a very abundant algae, followed by dinoflagellates as *Trithyrodinium* cf. *suspectum*, and in fewer amounts, *Cauveridinium membraniphorum* complex, *Oligosphaeridium pulcherrimum,* and other miscellaneous dinoflagellate cysts.

AASP 5 is represented by only one sample that belongs to Noto’s Facies A. This is the least varied assemblage observed in the entire AAS. The terrestrial sporomorph assemblage has seven spore species and 13 pollen species—in total, 20 species, about a fourth of the richness of the assemblages observed in AASP 1–2 and AASP 3–4. The spore assemblage is dominated by species of Cyatheaceae (*Cyathidites* spp.), a family of tropical ferns that include tree ferns, as in the other sections. Other important components of the assemblage are conifers. The pollen from these gymnosperms is also very abundant. The pollen from Taxodiaceae is more abundant here than in any of the other sections. Dinoflagellate cysts and freshwater algae are present but not as diverse or abundant as in the other sections (Figure 12). All these may indicate that Noto’s Facies A near AASP 5 was deposited in brackish conditions.

**Figure 12.**
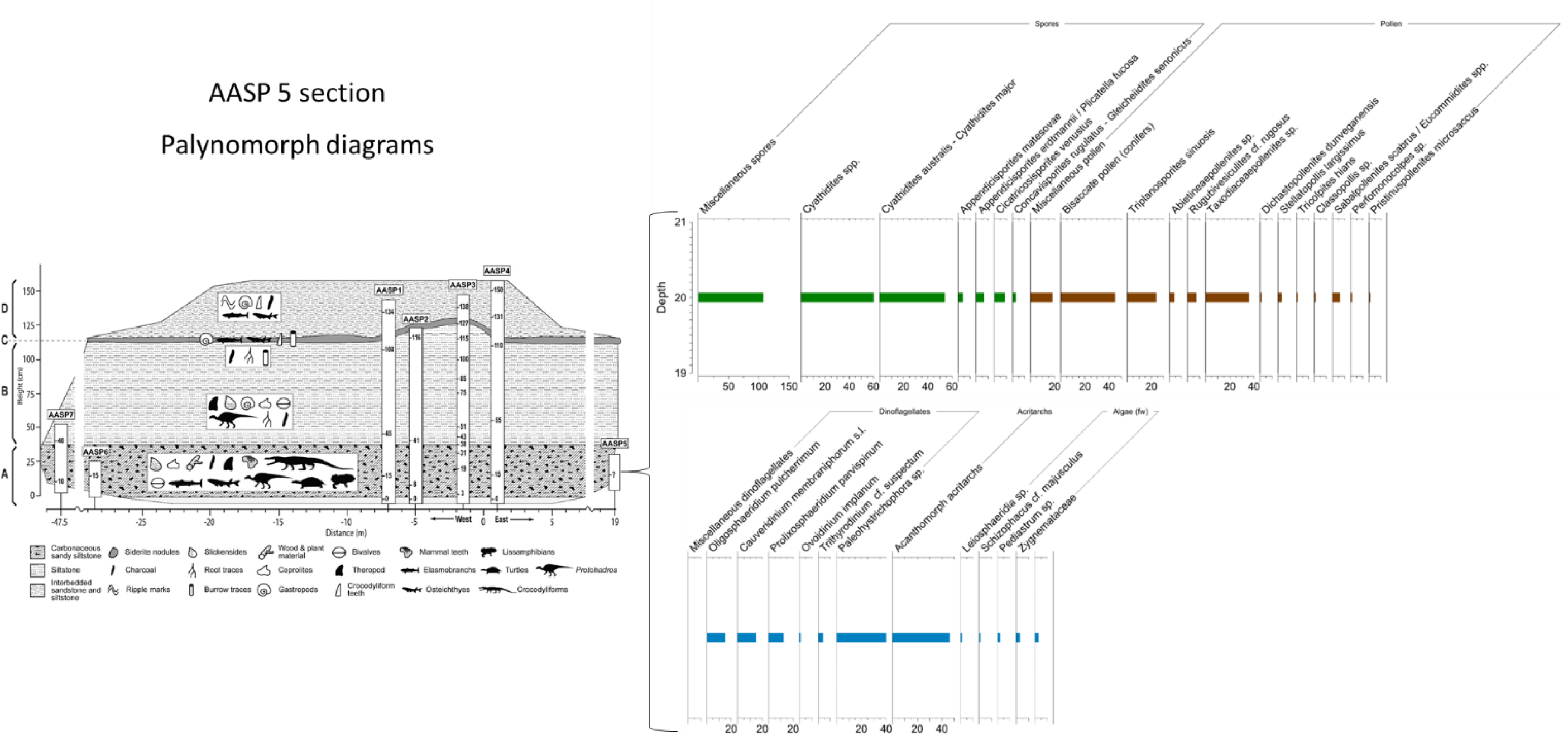
Section AASP 5. Pollen and spores’ abundance diagram and other palynomorphs diagram.

Characteristics of the assemblages in the combined sections AASP 6–7 and AASP 5 can be summarized as follows:

- The assemblages only represent Noto’s Facies A and the base of Noto’s Facies B. The assemblages’ richness (abundance and diversity) along the section is lower than in AASP 1–2 and AASP 3–4. Still, the most abundant components are spores from ferns or other seedless plants, followed by conifers’ pollen.
- The presence of relatively abundant Taxodiaceae pollen in AASP 6–7 and AASP 5, together with restricted assemblages of dinoflagellate cysts and low amounts of freshwater algae, distinguish them from the rest of the studied assemblages in AASP 1–2 and AASP 3–4.

## DISCUSSION

We will first analyze the different groups of plants present and their paleoecology based on the botanical affinities of the palynomorphs recovered at the AAS. After that, we will discuss the paleoenvironments and their relationships with Noto’s Facies identified in the outcrop, as well as some paleoclimate signals recorded. The last part of the discussion is the palynostratigraphy and the age interpretation for the AAS section.

### PALEOECOLOGY OF TERRESTRIAL PALYNOMORPHS

The palynological assemblage in the AAS comprises different types of palynomorphs. On the sporomorphs side, the spores of bryophytes, lycophytes, and pteridophytes represent the seedless plants. Gymnosperms are mainly represented by pollen from conifers, Cycadaceae, and Ephedraceae. The angiosperm pollen includes both monocots’ and dicots’ pollen grains. There are also palynomorphs from fungi (mostly spores and hyphae), algal spores, and filaments.

### BRYOPHYTES (MOSSES, HORNWORTS, AND LIVERWORTS)

Bryophytes consist of mosses (Musci), liverworts (Hepaticae), and hornworts (Anthocerotae), which are organizationally intermediate between green algae and green vascular plants (Playford & Dettmann, 1996; Sandersen, 2006). In the AAS assemblages, the following species were identified: *Contignisporites* sp., *Zlivisporis cenomanianus* (*Triporoletes cenomanianus*), *Camarozonosporites wrennii*, and *Foraminisporis* sp. A summary of botanical affinities and paleoenvironment conditions for some angiosperm markers is in Table 1.

**Table 1:** Terrestrial palynomorphs of bryophytes found at the AAS, their botanical affinity, and their paleoecological habitat. Based on (i) Rustad (2013); (m) Warny et al. (2012); (n) Costamagna et al. (2018).

| Taxa | Botanical Affinities | Paleoenvironment |
| --- | --- | --- |
| *Foraminisporis*<br>Krutzsch, 1959 | Bryophytes<br>(Embryophytes)? | Tend to grow only in places that are damp or humid. |
| *Camarozonosporites*<br>Potonié, 1956 | Bryophytes | The more humid parts of the lowland environment (n) |
| *Zlavisporis cenomanianus* | Bryophytes.<br>Hepaticae | Liverworts and hornworts favor soil with high moisture and tolerate minor flooding, but these plants would die if fully submerged (m) |

They are present in a few samples and usually a single specimen, except *Zlivisporis cenomanianus*, which is relatively abundant throughout the site sections. They are all considered to have been transported from the coastal and/or alluvial plains (Figure 13).

**Figure 13.**
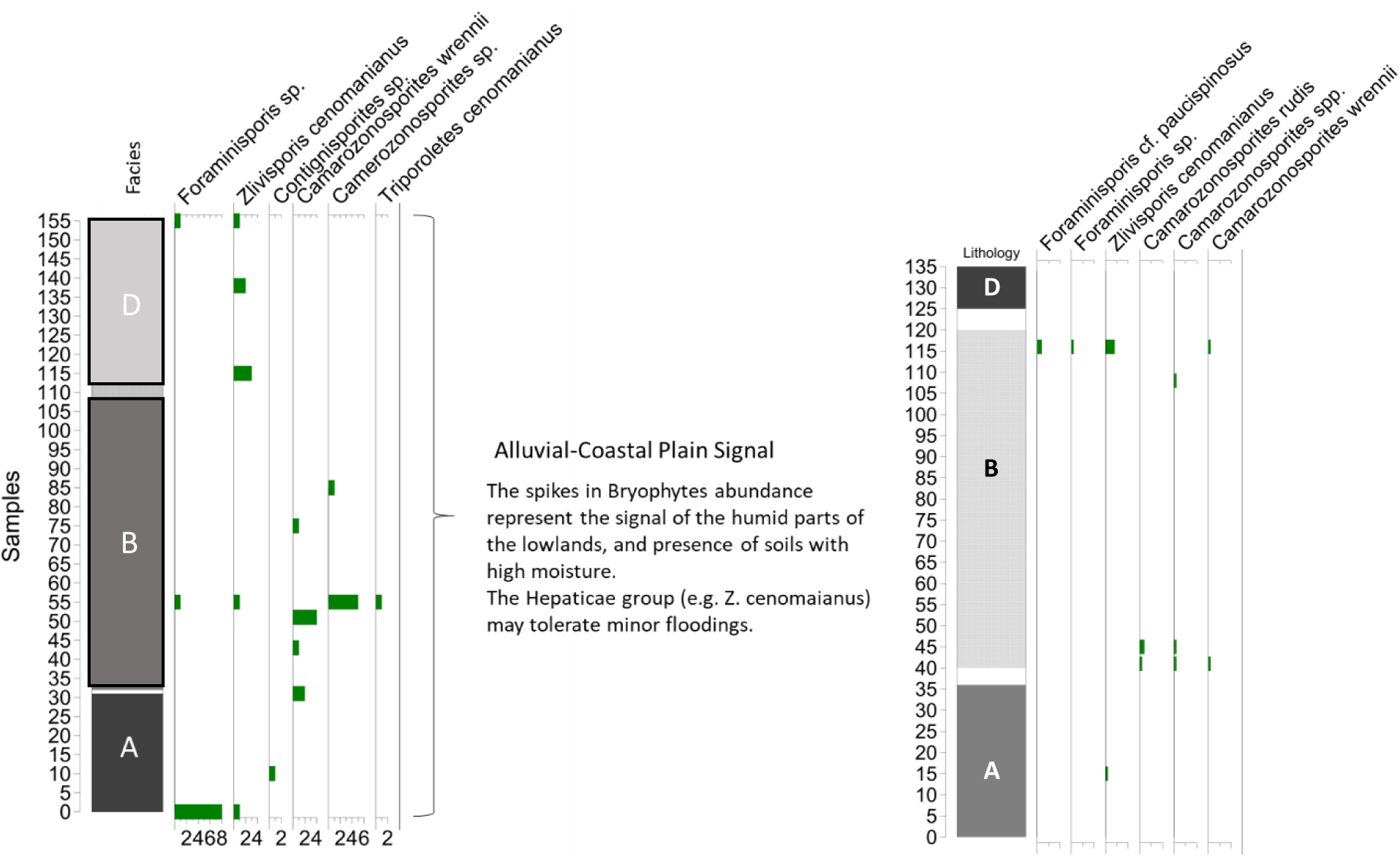
Bryophytes’ distribution in AASP 3–4 (left) and AASP 1–2 (right). These spores are considered to have been transported and represent the humid signal, sometimes even flooded parts of the alluvial and coastal plain. They are abundant in Noto’s Facies A and the lower part of Noto’s Facies B. *Zlivisporis cenomanianus* seems to take over the other bryophytes’ habitat in Noto’s Facies D.

### LYCOPHYTES (CLUBMOSSES, SPIKEMOSSES, AND QUILLWORTS)

In general, clubmosses or lycophytes are terrestrial or epiphytic vascular plants. “Common club moss, also known as running pine (*Lycopodium clavatum*), is native to open, dry woods and rocky places in the Northern Hemisphere” (Encyclopaedia Britannica, 2020).

During the Late Cretaceous, *Plicatella fucosa* inhabited lacustrine environments, and (a probably similar species) *Lycopodiumsporites marginatus*, lacustrine and coastal swamp environments (Hu, 2006). In the AAS assemblages, the following species of lycophytes’ spores were identified: *Lycopodiumsporites cf. dentimuratus*, *Lycopodiacidites tortus*, *Appendicisporites erdtmannii/Plicatella fucosa*, and *Reticulatisporites cf. arcuatus*.

A summary of paleoenvironmental and paleoclimate conditions for lycophytes’ to thrive is in Table 2. Figure 14 shows the occurrence of lycophytes in AASP 3–4.

**Figure 14.**
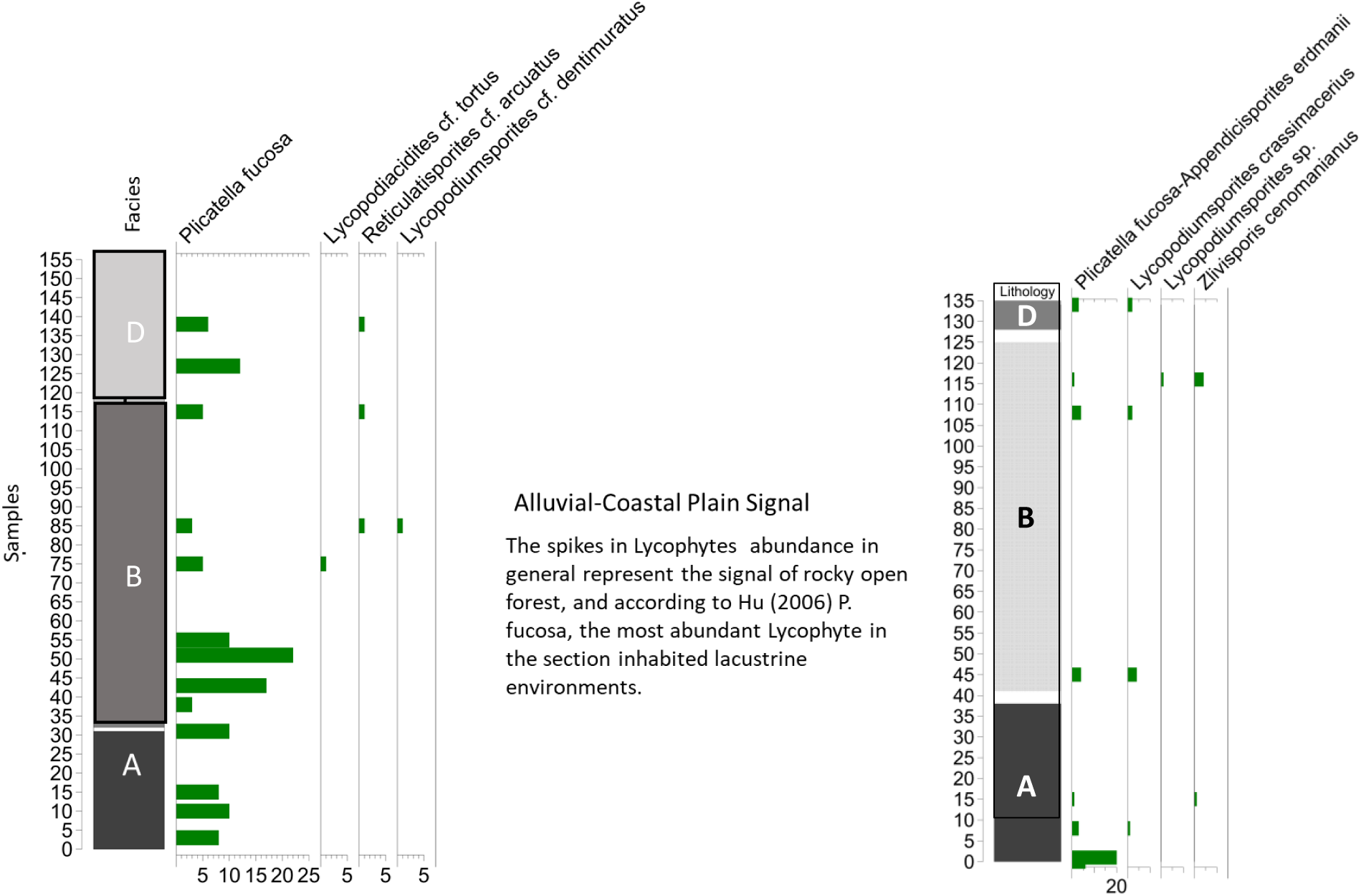
Lycophytes’ distribution and abundance in AASP 3–4 (left) and AASP 1–2 (right). These spores are considered to have been transported. The abundance of *P. fucosa* represents the signal from lakes and ponds in the alluvial and coastal plains, apparently more abundant during the sedimentation of Noto’s Facies A and the lower part of Noto’s Facies B.

**Table 2:** Terrestrial palynomorphs of lycophytes at the AAS, their botanical affinity, and their paleoecological habitat. Based on (i) Rustad (2013); (j) Gary et al. (2009).

| Taxa | Botanical Affinities | Paleoenvironment | Paleoclimate |
| --- | --- | --- | --- |
| *Lycopodiumsporites* (Thiergart) Delcourt & Sprumont, 1955 | Curly grass ferns. (Filicopsida—Ophioglossaceae) Filicales | Lycopodium, the terrestrial cosmopolitan fern, is included in clubmosses, firmosses, and quillworts. Lowland (i) | Warmer (?) (j) |

### PTERIDOPHYTES (FERNS, HORSETAILS)

The pteridophytes are seedless vascular plants. Although the term “pteridophyte” is no longer formal, it is still widely used, and the nomenclature continues to be discussed.

According to the Pteridophyte Phylogeny Group (2016), ferns and horsetails belong to the class Polypodiopsida but have different subclasses.

In the AAS assemblages, the following genera/species were identified (Figure 15):

**Figure 15.**
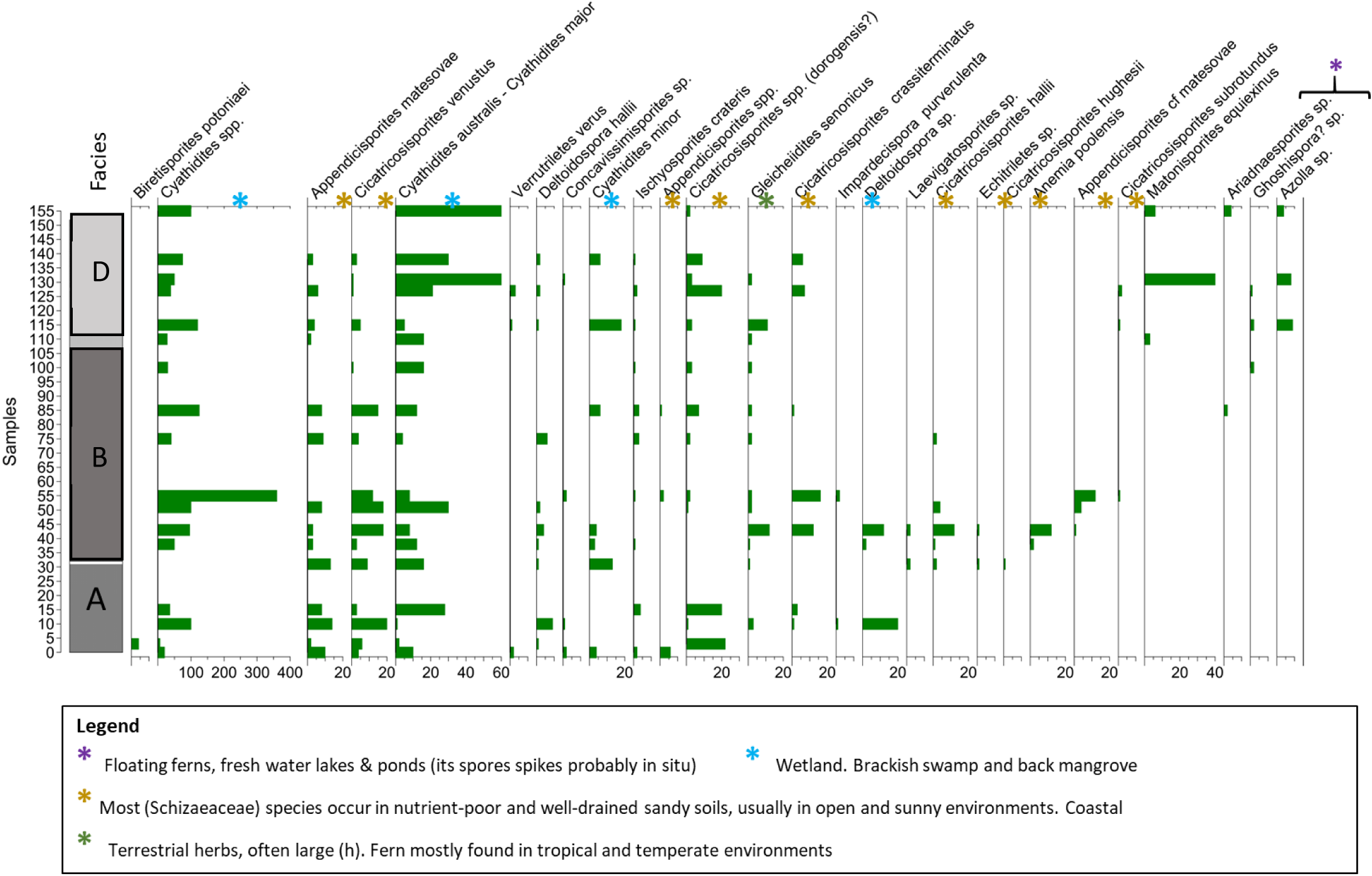
Ferns’ (pteridophytes’) distribution and abundance in AASP 3–4. Most of these spores are considered to have been transported to the section. However, some of the lowland brackish assemblage represented by the genera *Cyatheacidites* and *Deltoidospora* may be *in situ* or very close to their living environment. Those species are the most abundant components of the sporomorph assemblage.

*Anemia poolensis*, *Appendicisporites* spp., *Appendicisporites matesovae*, *Appendicisporites* cf*. matesovae*, *Ariadnaesporites* sp., *Biretisporites potoniaei*, *Camerozonosporites* sp., *Camarozonosporites rudis*, *Cicatricosisporites* sp. (*C. dorogensis*?), *Cicatricosisporites venustus*, *Cicatricosisporites crassiterminatus*, *Cicatricosisporites hallii*, *Cicatricosisporites hughesii*, *Cicatricosisporites subrotundus*, *Concavisporites rugulatus–Gleicheiidites senonicus*, *Cyathidites* spp., *Concavissimisporites* sp., *Cyathidites australis–Cyathidites major*, *Cyathidites minor*, *Deltoidospora* sp., *Deltoidospora hallii*, *Distaltriangulisporites* sp., *Echitriletes* sp., *Ghoshispora?* sp., *Impardecispora purverulenta*, *Ischyosporites crateris*, *Laevigatosporites* sp., *Matonisporiles excavates*, *Matonisporites equiexinus*, *Pilosisporites ericius*, *Verrutriletes verus*.

Freshwater floating ferns: *Ariadnaesporites* sp. (Salviniaceae, floating fern), *Azolla* sp. (Azollaceae, floating fern), *Glomospira* sp. (Salviniaceae, floating fern).

As Table 3 summarizes, ferns are cosmopolitan and can be found in many environments, from the uplands to the coastal plains. These plants prefer warm environments and need humid conditions (Abbink et al., 2004), so they are often found in swampy and even brackish water environments. They are most abundant in tropical and temperate climates. Some floating ferns may be abundant in lakes and ponds; abundance spikes indicate they are probably *in situ*.

**Table 3:**
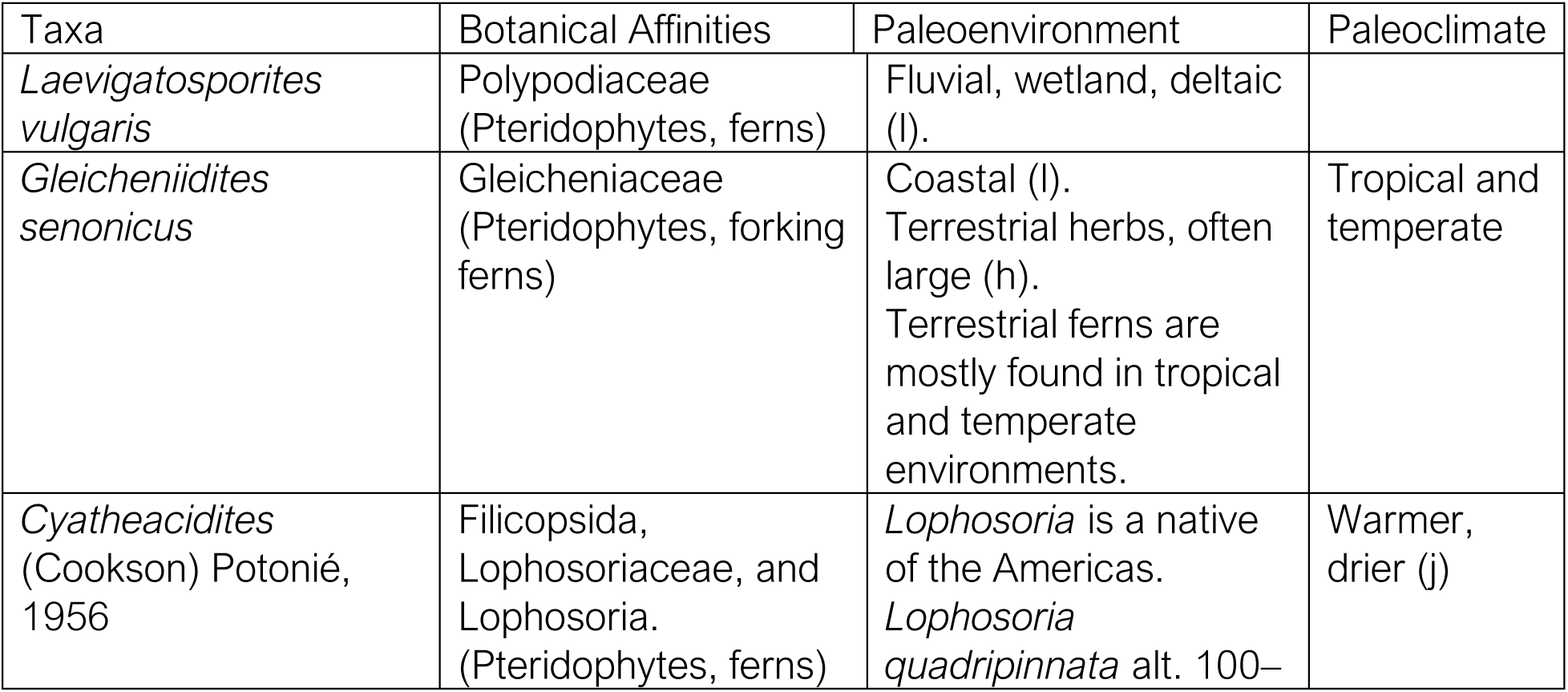
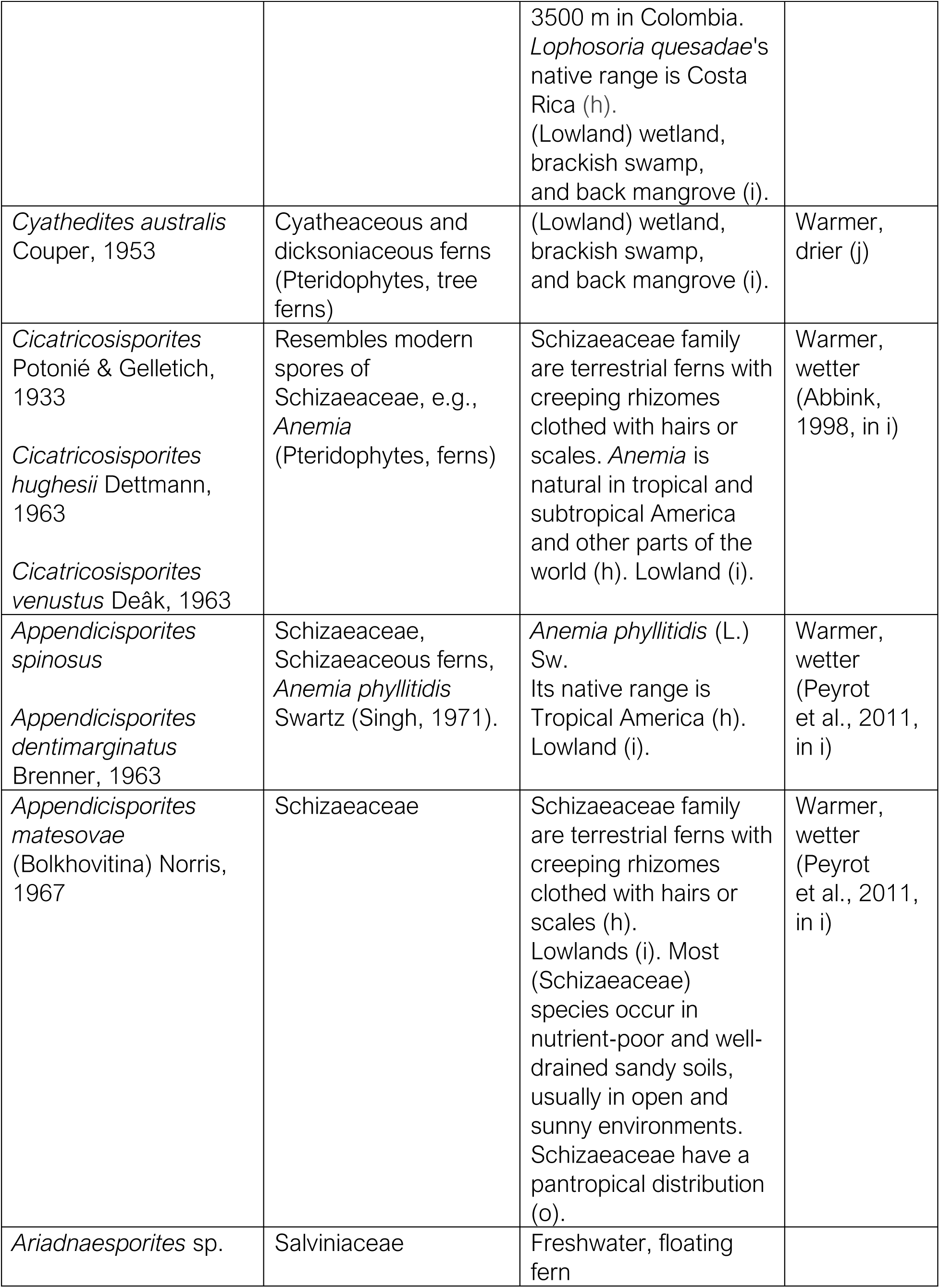
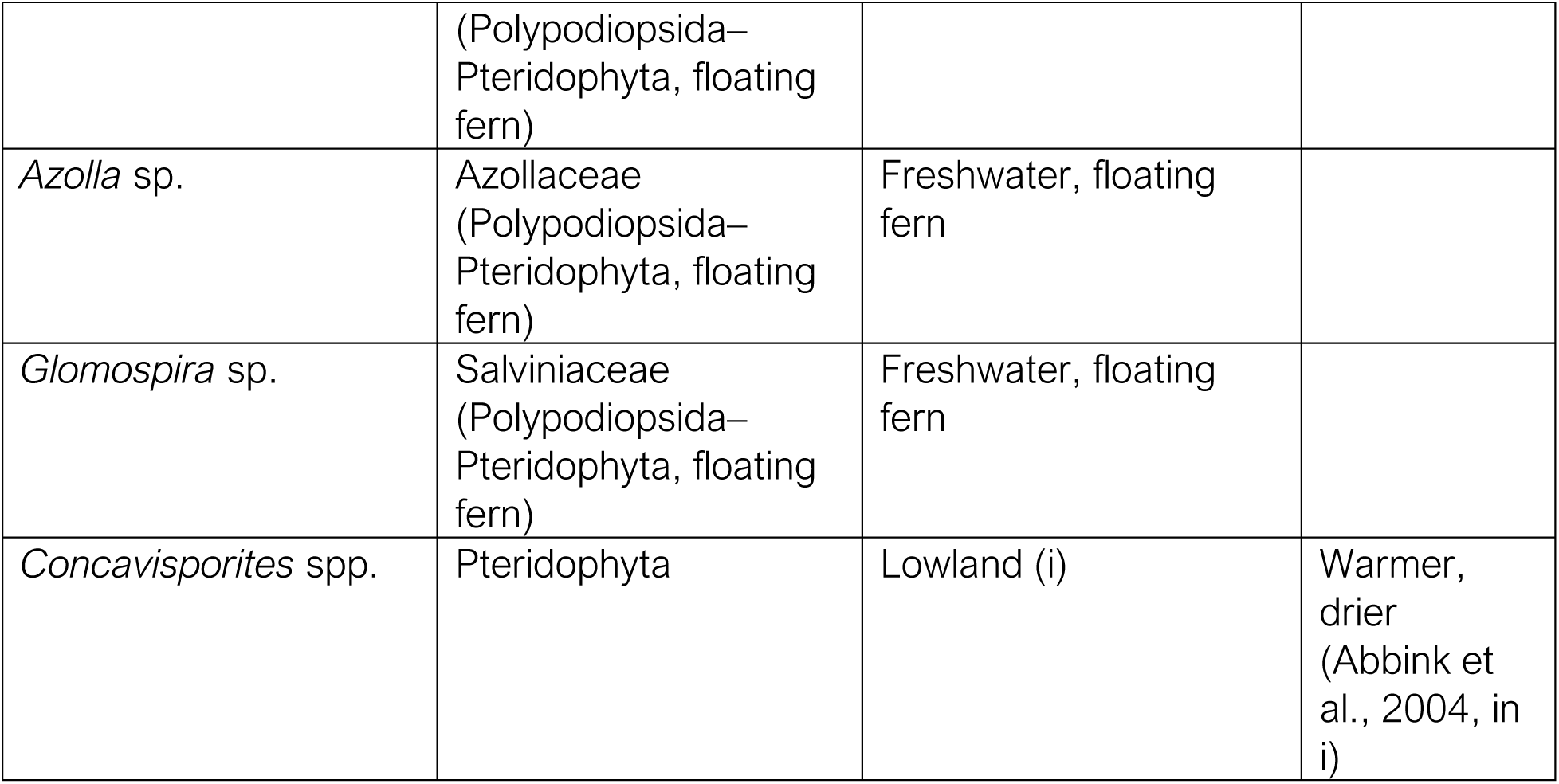
Terrestrial palynomorphs from pteridophytes found at the AAS, their botanical affinity, and their paleoecological habitat. Based on (a) Alvin (1982); (b) Armstrong (n.d.); (c) the Gymnosperm Database (n.d.); (d) Pittermann et al. (2012); (e) Basic Biology (2015); (f) Akyuz et al. (2016); (g) Boufford and Sytsma (2017); (h) Kew Royal Botanical Gardens (n.d.); (i) Rustad (2013); (j) Gary et al. (2009); (k) Tschudy et al. (1984); (l) Barron (2015); (o) Labiak and Karol (2017).

| Taxa | Botanical Affinities | Paleoenvironment | Paleoclimate |
| --- | --- | --- | --- |
| *Laevigatosporites vulgaris* | Polypodiaceae (Pteridophytes, ferns) | Fluvial, wetland, deltaic (l). |  |
| *Gleicheniidites senonicus* | Gleicheniaceae (Pteridophytes, forking ferns) | Coastal (l).<br>Terrestrial herbs, often large (h).<br>Terrestrial ferns are mostly found in tropical and temperate environments. | Tropical and temperate |
| *Cyatheacidites* (Cookson) Potonié, 1956 | Filicopsida, Lophosoriaceae, and Lophosoria. (Pteridophytes, ferns) | *Lophosoria* is a native of the Americas.<br>*Lophosoria quadripinnata* alt. 100– | Warmer, drier (j) |
|  |  | 3500 m in Colombia. *Lophosoria quesadae*'s native range is Costa Rica (h). (Lowland) wetland, brackish swamp, and back mangrove (i). |  |
| *Cyathedites australis*<br>Couper, 1953 | Cyatheaceous and dicksoniaceae ferns (Pteridophytes, tree ferns) | (Lowland) wetland, brackish swamp, and back mangrove (i). | Warmer, drier (j) |
| *Cicatricosisporites*<br>Potonié & Gelletich, 1933<br>*Cicatricosisporites hughesii* Dettmann, 1963<br>*Cicatricosisporites venustus* Deák, 1963 | Resembles modern spores of Schizaeaceae, e.g., *Anemia* (Pteridophytes, ferns) | Schizaeaceae family are terrestrial ferns with creeping rhizomes clothed with hairs or scales. *Anemia* is natural in tropical and subtropical America and other parts of the world (h). Lowland (i). | Warmer, wetter (Abbink, 1998, in i) |
| *Appendicisporites spinosus*<br>*Appendicisporites dentimarginatus*<br>Brenner, 1963 | Schizaeaceae, Schizaeaceous ferns, *Anemia phyllitidis* Swartz (Singh, 1971). | *Anemia phyllitidis* (L.) Sw.<br>Its native range is Tropical America (h). Lowland (i). | Warmer, wetter (Peyrot et al., 2011, in i) |
| *Appendicisporites matesovae*<br>(Bolkhovitina) Norris, 1967 | Schizaeaceae | Schizaeaceae family are terrestrial ferns with creeping rhizomes clothed with hairs or scales (h). Lowlands (i). Most (Schizaeaceae) species occur in nutrient-poor and well-drained sandy soils, usually in open and sunny environments. Schizaeaceae have a pantropical distribution (o). | Warmer, wetter (Peyrot et al., 2011, in i) |
| *Ariadnaesporites* sp. | Salviniaceae | Freshwater, floating fern |  |

Table 3: Terrestrial palynomorphs from pteridophytes found at the AAS, their botanical affinity, and their paleoecological habitat.
|  |  |  |  |
| --- | --- | --- | --- |
|  | (Polypodiopsida–<br>Pteridophyta, floating<br>fern) |  |  |
| *Azolla* sp. | Azollaceae<br>(Polypodiopsida–<br>Pteridophyta, floating<br>fern) | Freshwater, floating<br>fern |  |
| *Glomospira* sp. | Salviniaceae<br>(Polypodiopsida–<br>Pteridophyta, floating<br>fern) | Freshwater, floating<br>fern |  |
| *Concavisporites* spp. | Pteridophyta | Lowland (i) | Warmer,<br>drier<br>(Abbink et<br>al., 2004, in<br>i) |

### GYMNOSPERMS

Gymnosperms are plants that produce seeds and pollen but do not have flowers. They include conifers, cycads, ginkgos, and gnetophytes. Conifers belonging to Taxodiacea and Cupressacea are part of the same natural group based on molecular studies (The Gymnosperm Database, n.d.).

The following pollen species belonging to gymnosperms are present in the AAS sections:

Conifers und.: *Abietineaepollenites* sp., *Abietineaepollenites microreticulatus*, *Callialasporites* sp., *Callialasporites dampieri*, *Callialasporites segmentatus*, *Inaperturopollenites* spp., *Podocarpidites* sp., *Podocarpidites biformis*, *Pristinuspollenites microsaccus*, *Rugubivesiculites rugosus*, *Rugubivesiculites* sp., indet. bisaccate pollen.

During the Jurassic/Cretaceous era, a warm climate without significant seasonal variations was associated with a high frequency of *Araucariacites* and *Callialasporites*; they frequently grew near the shore, as they could withstand the influence of salt wind, according to Abbink et al. (2004), Reyre (1980), and Mohr (1989). Although we have not identified *Araucariacites* in the AAS sections, *Callialasporites* are sporadically present in Noto’s Facies B.

Pinophyta (extinct Cheirolepidiaceae): *Classopollis* sp., *Classopollis torosa*, *Classopollis classoides.* According to Abbink et al. (2004), the Cheirolepidiaceae were drought-resistant shrubs and trees, with some taxa showing xeromorphic and thermophilic characteristics. These plants might have inhabited sandy bars, coastal islands, and even mangroves (Abbink et al., 2004).

Taxodiaceae: *Taxodiaceaepollenites* sp. Some extant members of the Taxodiaceae prefer humid environments Abbink et al. (2004). The swamp cypress family belongs to this group.

Cycadophytes: *Cycadopites* sp., *Sabalpollenites scabrus*/*Eucommiidites* spp. According to Abbink et al. (2004), today’s Cycadales are typically found in tropical regions and have adapted to survive droughts. In general, these plants grew in lowland vegetation.

Gnetophytes (Gnetales)*: Ephedripites sp*.

A summary of favorable paleoenvironmental and paleoclimate conditions for some gymnosperm species present in the AAS sections is in Table 4.

**Table 4.**
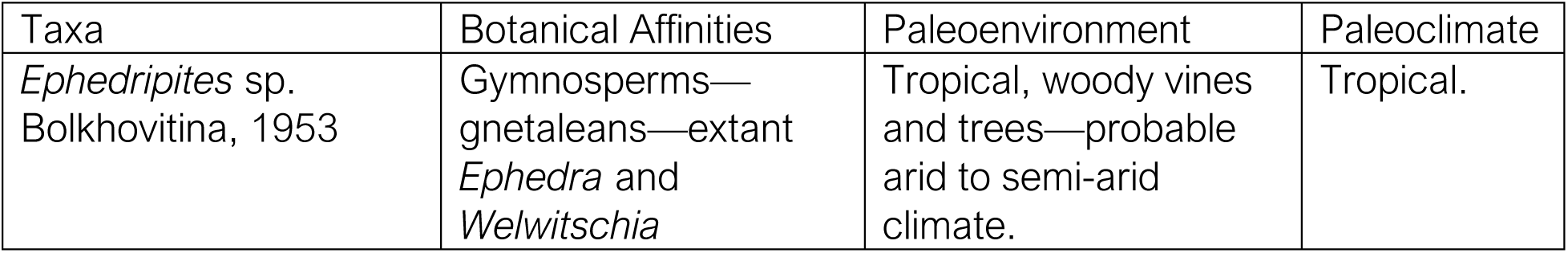
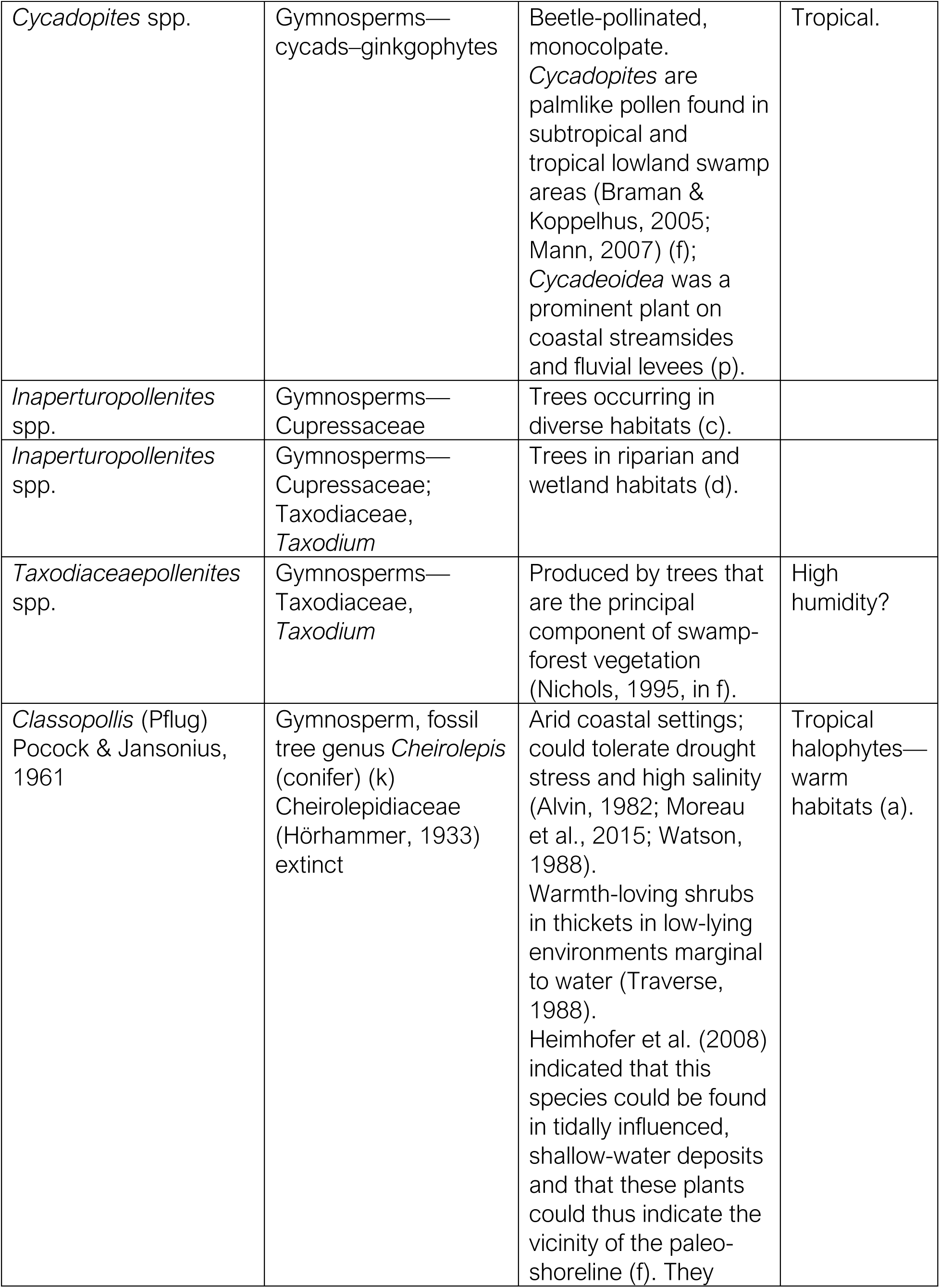
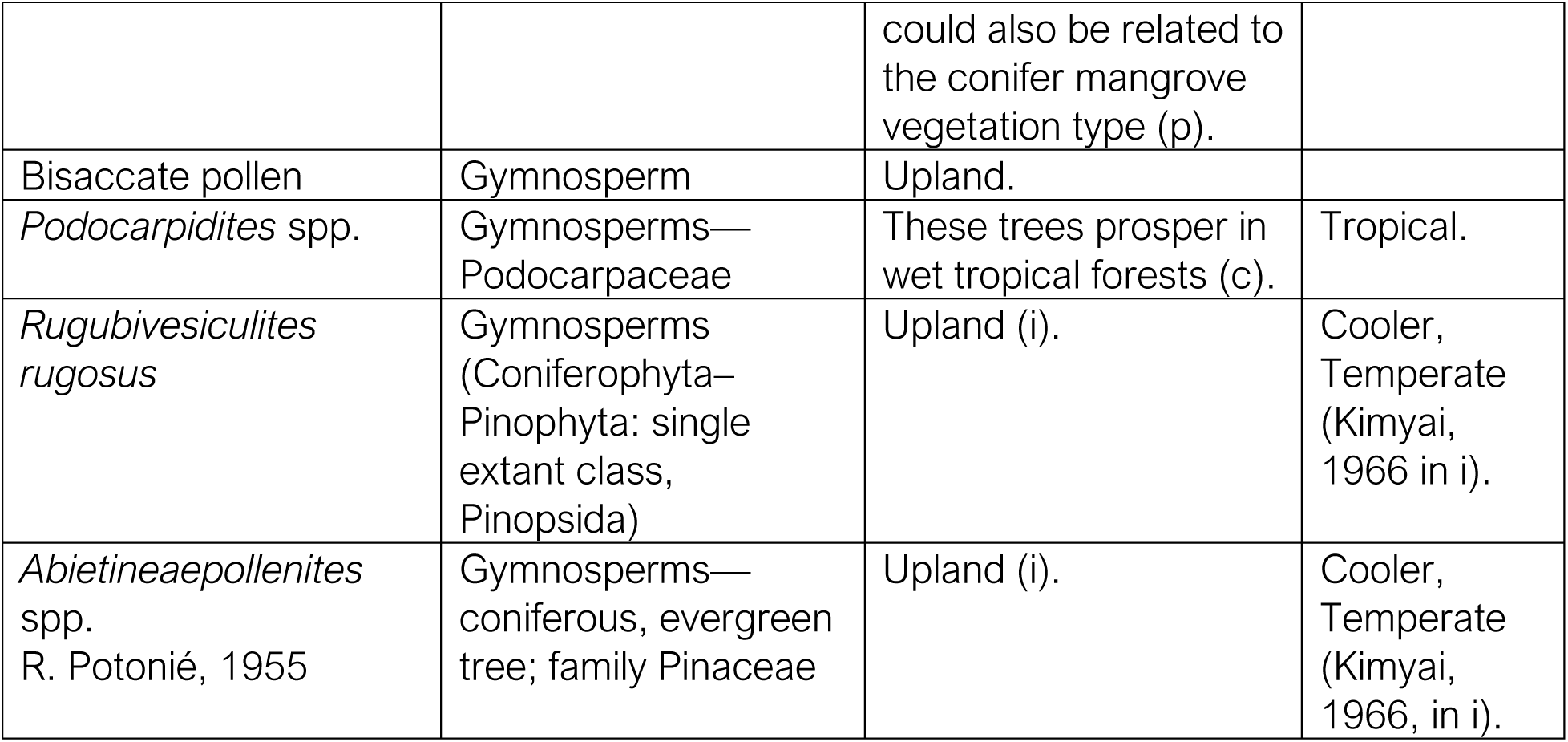
Terrestrial palynomorphs from gymnosperms found at the AAS, their botanical affinity, and their probable paleoecological habitat. Based on (a) Alvin (1982); (b) Armstrong (n.d.); (c) the Gymnosperm Database (n.d.); (d) Pittermann et al. (2012); (e) Basic Biology (2015); (f) Akyuz et al. (2016); (g) Boufford and Sytsma (2017); (h) Kew Royal Botanical Gardens (n.d.); (i) Rustad (2013); (j) Gary et al. (2009); (k) Tschudy et al. (1984); (l) Barron (2015); (p) Retallack and Dilcher (1981).

| Taxa | Botanical Affinities | Paleoenvironment | Paleoclimate |
| --- | --- | --- | --- |
| *Ephedripites* sp.<br>Bolkhovitina, 1953 | Gymnosperms—<br>gnetales—extant<br>*Ephedra* and<br>*Welwitschia* | Tropical, woody vines<br>and trees—probable<br>arid to semi-arid<br>climate. | Tropical. |
| *Cycadopites* spp. | Gymnosperms—<br>cycads—ginkgophytes | Beetle-pollinated,<br>monocolpate.<br>*Cycadopites* are<br>palmlike pollen found in<br>subtropical and<br>tropical lowland swamp<br>areas (Braman &<br>Koppelhus, 2005;<br>Mann, 2007) (f);<br>*Cycadeoidea* was a<br>prominent plant on<br>coastal streambanks<br>and fluvial levees (p). | Tropical. |
| *Inaperturopollenites*<br>spp. | Gymnosperms—<br>Cupressaceae | Trees occurring in<br>diverse habitats (c). |  |
| *Inaperturopollenites*<br>spp. | Gymnosperms—<br>Cupressaceae;<br>Taxodiaceae,<br>*Taxodium* | Trees in riparian and<br>wetland habitats (d). |  |
| *Taxodiaceapollenites*<br>spp. | Gymnosperms—<br>Taxodiaceae,<br>*Taxodium* | Produced by trees that<br>are the principal<br>component of swamp-forest vegetation<br>(Nichols, 1995, in f). | High<br>humidity? |
| *Classopollis* (Pflug)<br>Pocock & Jansonius,<br>1961 | Gymnosperm, fossil<br>tree genus *Cheirolepis*<br>(conifer) (k)<br>Cheirolepidiaceae<br>(Hörhammer, 1933)<br>extinct | Arid coastal settings;<br>could tolerate drought<br>stress and high salinity<br>(Alvin, 1982; Moreau<br>et al., 2015; Watson,<br>1988).<br>Warmth-loving shrubs<br>in thickets in low-lying<br>environments marginal<br>to water (Traverse,<br>1988).<br>Heimhofer et al. (2008)<br>indicated that this<br>species could be found<br>in tidally influenced,<br>shallow-water deposits<br>and that these plants<br>could thus indicate the<br>vicinity of the paleo-shoreline (f). They | Tropical<br>halophytes—<br>warm<br>habitats (a). |

Table 4. Terrestrial palynomorphs from gymnosperms found at the AAS, their botanical affinity, and their probable paleoecological habitat.
|  |  |  |  |
| --- | --- | --- | --- |
|  |  | could also be related to the conifer mangrove vegetation type (p). |  |
| Bisaccate pollen | Gymnosperm | Upland. |  |
| *Podocarpidites* spp. | Gymnosperms—Podocarpaceae | These trees prosper in wet tropical forests (c). | Tropical. |
| *Rugubivesiculites rugosus* | Gymnosperms (Coniferophyta–Pinophyta: single extant class, Pinopsida) | Upland (i). | Cooler, Temperate (Kimyai, 1966 in i). |
| *Abietineaepollenites* spp.<br>R. Potonié, 1955 | Gymnosperms—coniferous, evergreen tree; family Pinaceae | Upland (i). | Cooler, Temperate (Kimyai, 1966, in i). |

Based on the known thriving conditions of the species that produce some of the identified gymnosperm pollen species, we could deduct the provenance signal from the upper parts of the drainage basin. In the AASP sections, Conifers are generally the predominant group of gymnosperms, and although we consider many of them transported, their presence may give clues about the conditions that prevailed upstream of the final place of sedimentation. However, there are some exceptions, like the case of those Gymnosperms growing in lowlands under swampy brackish conditions, where pollen may have had a higher probability of being trapped and preserved in situ. (Figure 16).

**Figure 16.**
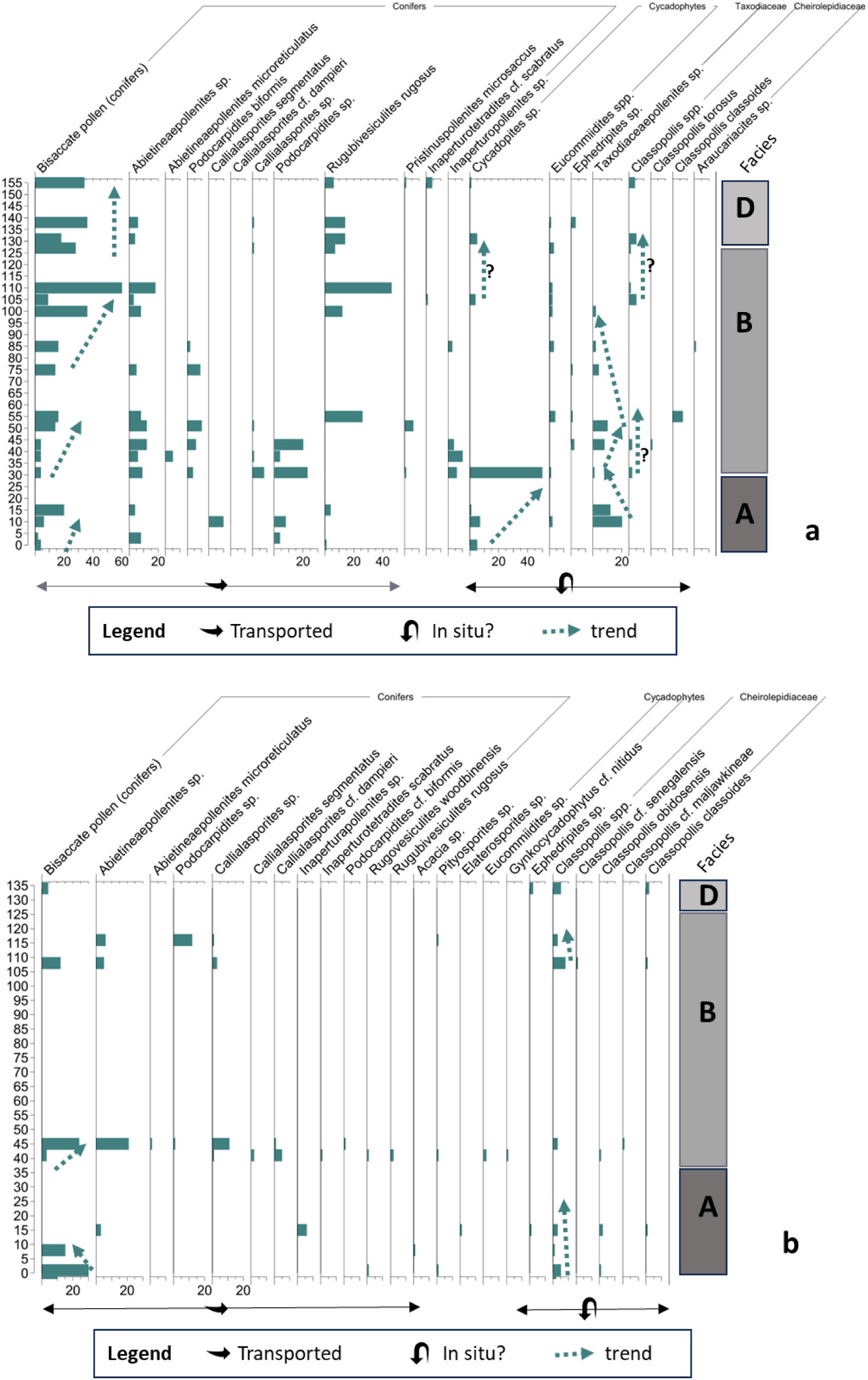
Gymnosperms histogram for AASP 3–4 (a) and AASP 1–2 (b). The conifers are dominant in the assemblages throughout the sections. There are some differences, e.g., in AASP 3–4, Taxodiaceae has an increase within Noto’s Facies A, probably indicating brackish swampy conditions, which is followed by a strong increase of Cycadophytes, which seems to be replacing the Taxodiaceae in the habitat for a relatively short time. After that, in Facies B, Taxodiaceae regains its habitat but disappears towards the upper part of the facies. Cheirolepidiaceae become continuously present throughout most of Noto’s Facies B and D, a probable indication of the climate becoming relatively dryer. Taxodiaceae are absent in AASP 1–2, but Cheirolepidiaceae seem present throughout the entire section.

### ANGIOSPERMS

These are flowering plants that produce seeds and pollen. This is the most diverse group of land plants, and it evolved during the Late Cretaceous. Angiosperms have two classes: Liliopsida (monocotyledons or monocots) and Magnoliopsida (dicotyledons or dicots), based on the seed’s number of cotyledons. These two classes of angiosperms produce different types of pollen. Monocots have pollen with a single furrow or pore, while dicots produce pollen with three or more furrows and/or pores. The following pollen species belonging to angiosperms are present in the AAS sections:

Monocotyledons: *Monosulcites* sp*., Monosulcites inspissiatus*, *Foveomonocolpites* sp., *Liliacidites* sp., *Perfomonocolpites* sp., *Proxapertites* spp., *Dichastopollenites dunveganensis*, *Liliacidites* cf. *inaequalis*, *Stellatopollis* sp., *Stellatopollis largissimus*, *Tucanopollis* sp., *lnaperturotetradites* cf. *scabratus* (Figure 17).

**Figure 17.**
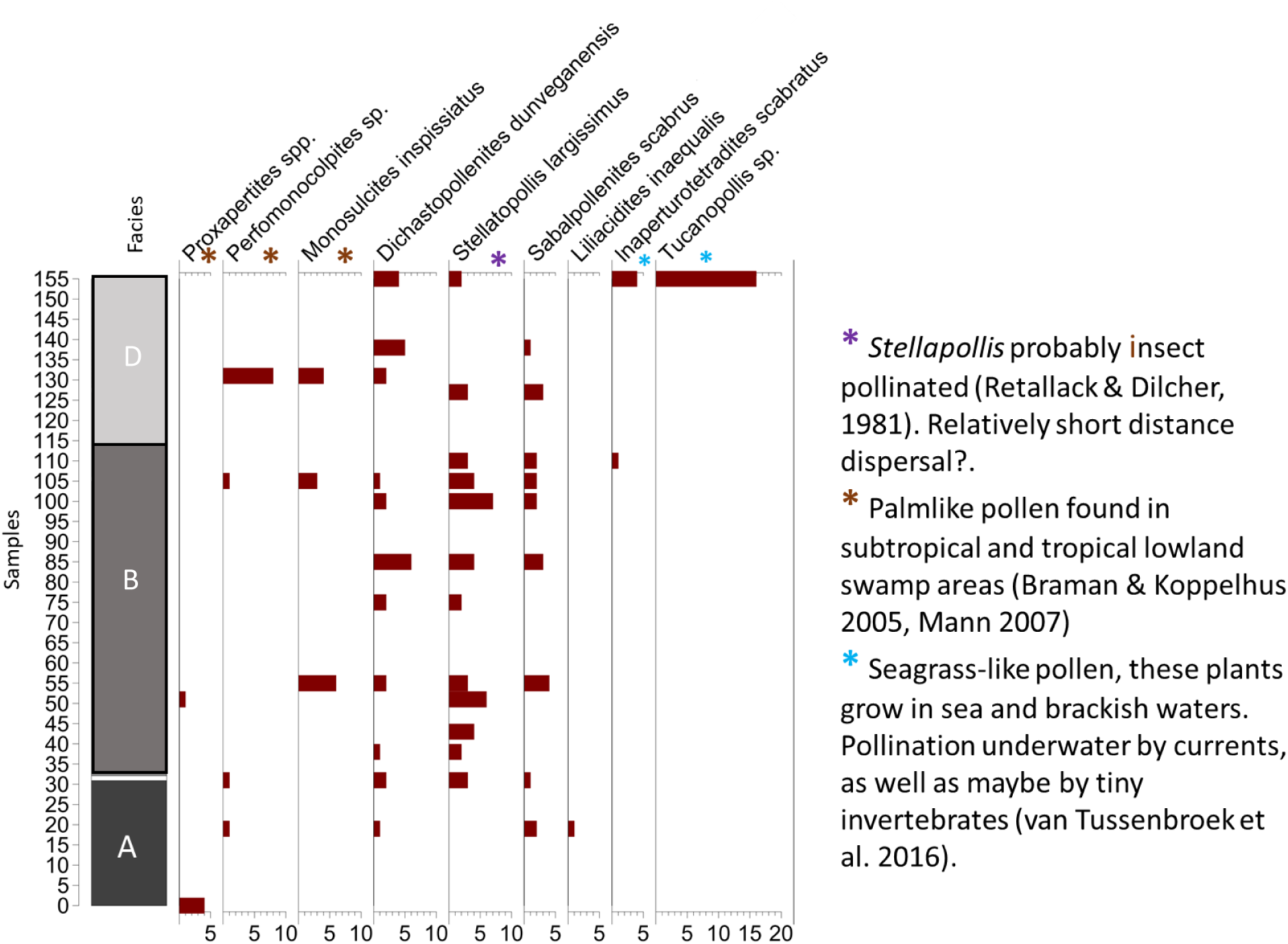
The monocot angiosperms’ distribution chart points to the presence of pollen coming from at least two different basin areas. The first is the swampy lowland areas, represented by the monosulcate and proxapertites pollen, probably from a type of palm forest; the second is inaperturate, thin-walled pollen and tetrads from marine seagrasses, representing the in situ part of some assemblages.

Dicotyledons: *Artiopollis indivisus*, *Aesculiidites dubius*, *Cupuliferoipollenites* sp. *Cupuliferoidaepollenites* cf*. microscabratus*, *Psilatricolpites* sp., *Retimonocolpites* sp., *Rousea* cf. *georgensis*, *Retitricolpites* sp., *Retitricolpites* cf. *geranioides*, *Tricolpites* spp., *Tricolpites hians*, *Retitricolpites maximus* (Figure 18).

**Figure 18.**
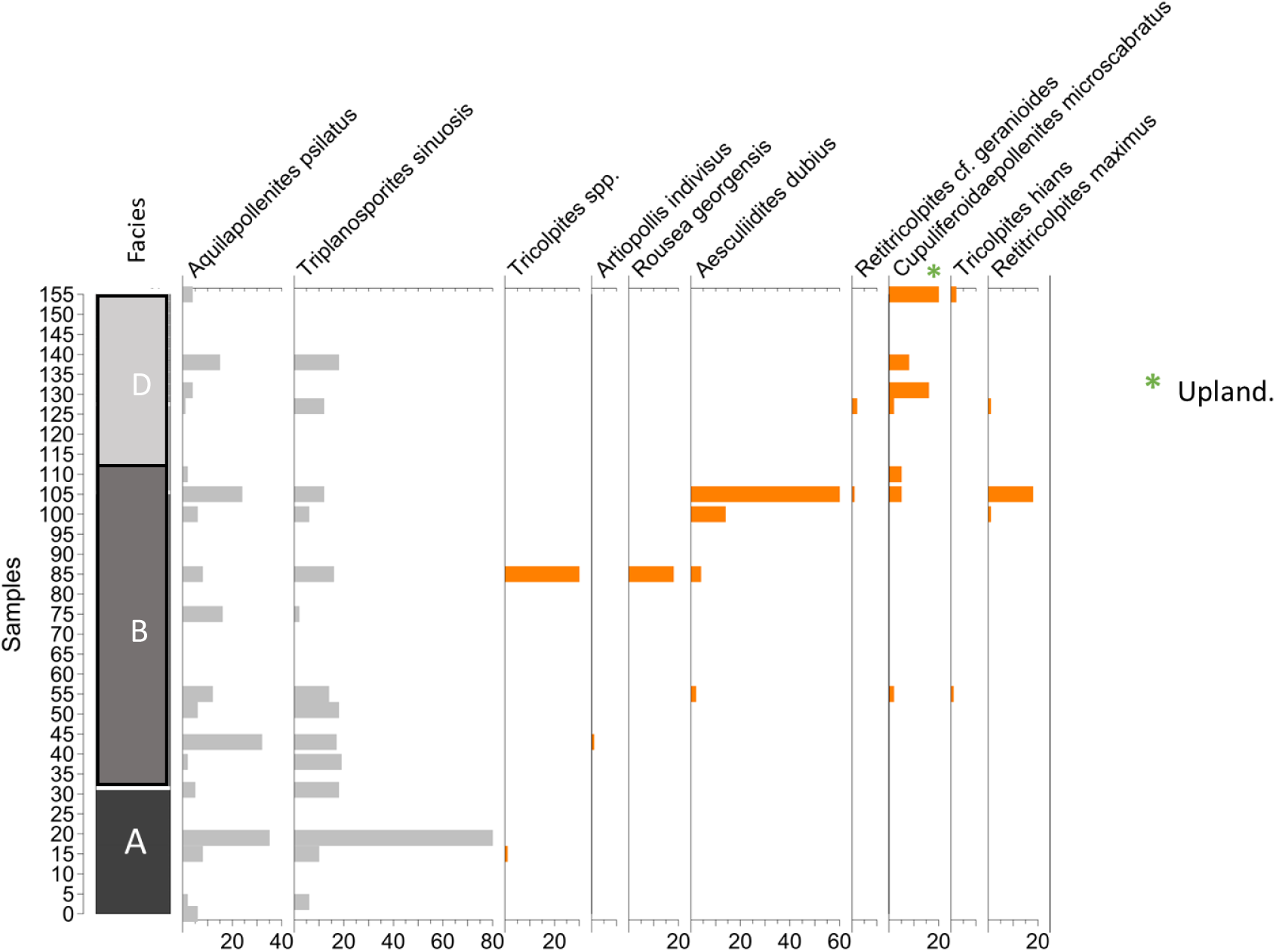
The dicot angiosperms’ distribution chart shows a very different occurrence pattern between *Aquilapollenites psilatus* and *Triplanosporites sinuosis* (in grey) and the other dicot angiosperms (in orange). While the former two occur abundantly throughout the entire section, more similar to the pteridophytes’ distribution, the second group starts appearing in the section with rare occurrences from the upper part of Noto’s Facies A to the lower part of Noto’s Facies B, but in the upper part of Noto’s Facies B and throughout Facies D, they “bloom.” There are two different considerations to make: First, this graph shows that *Aquilapollenites psilatus* and *Triplanosporites sinuosis* do not have the same paleoecological requirements; second, the other dicot angiosperms show an apparent base occurrence here, which could be either due to migration into the basin or a true record of the first occurrence of dicot angiosperms in this area of the CWIS geological record.

If angiosperms, then probably dicotyledons (?): *Triplanosporites sinuosis*, *Aquilapollenites psilatus*.

Table 5 summarizes the botanical affinities and paleoenvironment conditions for some angiosperm markers.

**Table 5:** Some terrestrial palynomorphs from angiosperms, their possible botanical affinities, and their paleoenvironmental conditions. Based on (a) Alvin (1982); (b) Armstrong (n.d.); (c) the Gymnosperm Database (n.d.); (d) Pittermann et al. (2012); (e) Basic Biology (2015); (f) Akyuz et al. (2016); (g) Boufford and Sytsma (2017); (h) Kew Royal Botanical Gardens (n.d.); (i) Rustad (2013); (j) Gary et al. (2009); (k) Tschudy et al. (1984); (l) Barron (2015); (p) Retallack and Dilcher (1981).

| Taxa | Botanical Affinities | Paleoenvironment |
| --- | --- | --- |
| *Monocolpites* spp./<br>*Monosulcites* spp. | Angiosperms—<br>monocotyledons. | Palmlike pollen is found<br>in subtropical and<br>tropical lowland swamp<br>areas (Braman &<br>Koppelhus, 2005;<br>Mann, 2007, in f). |
| *Cupuliferoidaepollenites*<br>sp. | Angiosperm (probably<br>Fagaceae). | Upland. |
| *Cupuliferoidaepollenites*<br>*microscabratus* | Angiosperm (probably<br>Fagaceae). | Upland. |

Table 5: Some terrestrial palynomorphs from angiosperms, their possible botanical affinities, and their paleoenvironmental conditions.
|  |  |  |
| --- | --- | --- |
| *Stellapollis* | *Stellapollis barghoornii* | It is comparable to the pollen of modern insect-pollinated angiosperms. |
| *Tricolpites* spp | Angiosperm—<br>Platanaceae (?). | They have little ecologic value because they could represent various angiosperms with various ecologic preferences (Nichols, 1995, in f). |
| Tricolporate & triporate pollen | Angiosperms—<br>eudicotyledons (?). | “Abundant in the majority of land-based ecosystems in tropical and temperate environments” (e). |

Most monocots are primarily tropical. Many are grasses, sedges, and palms and can be found in many different environments, e.g., along with and in streams and ponds; in coastal marine environments, there are monocots in deserts, and the largest floating and submerged aquatic plants are monocots, e.g., water hyacinths and “seagrasses” (UCMP, n.d., “Monocots”). Seagrasses grow in sea and brackish waters, mainly on gently sloping, protected coastlines, commonly found in shallow depths of 3 to 9 feet (Reynolds, 2018).

These plants’ pollen is usually spherical to ellipsoidal inaperturate grains, single or in tetrads with a thin exine (Ackerman, 2006; Tanaka et al., 2006). Based on the above-mentioned pollen characteristics, some species in the section may belong to seagrasses, e.g., *Inaperturotetradites scabratus* and *Tucanipollis* sp.

### FUNGAL AND ALGAL REMAINS

Fungal remains are part of the terrestrial realm, where they live in plants, even on the leaves’ surfaces, where they can be harmful to the plant, but they also live in soils on decaying plant tissues, degrading the organic matter. Fungi can also thrive in aquatic freshwater habitats. Freshwater habitats that include river and stream banks, ponds, and lakes, as well as forest tree canopies, all form excellent substrates for the aquatic and aero-aquatic fungi that are often common in palynological slides (Nuñez Otaño et al., 2021). A catalog of freshwater fungi can be found at https://freshwaterfungi.org/.

The following taxa are present in the AAS sections:

Fungal remains: linear septate fungal hypha, fungal spores, and fungal fruit bodies (Figure 19).

**Figure 19.**
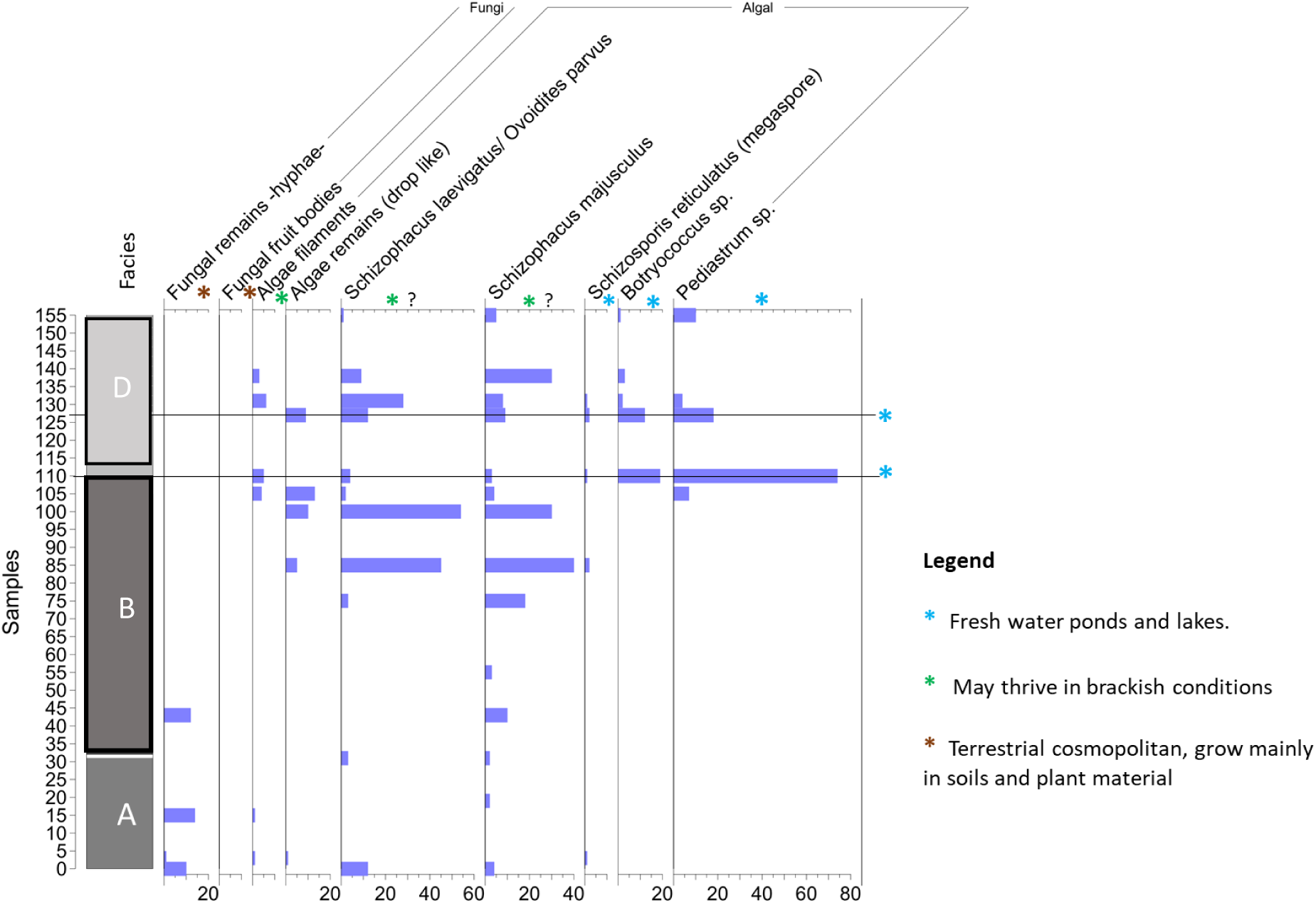
Distribution chart of the fungal remains and algal spores and colonies. The graphs show an increase in *Pediastrum* and *Botryococcus* towards the upper part of Noto’s Facies B, and in the lowermost Noto’s Facies D, they are assumed to be *in situ*.

Algae are a varied group of plant-like organisms. They can be single-celled, to colonial, or massive multicellular forms, all of them have chlorophyll, and provide the energetic foundation of the food chain. They are essential to aquatic ecosystems, but changes in the environment, both chemical and physical, have a significant effect on the abundance levels and community structure of algae.

The following types of algae are present in the AAS sections:

Algal: filaments, other algae remains (drop-shaped), *Schizophacus laevigatus*/*Ovoidites parvus* (zygospores or aplanospores of the modem zygnemataceous chlorophyte algae *Spirogyra*, according to van Geel, 1976), *Schizophacus majusculus* (algal cyst), *Schizosporis reticulatus* (Chlorophyta according to Dettman, 1986), *Pediastrum* sp. (colonial algae), and *Botryococcus* sp. (planktic colonial algae, a good indicator of the changes in the trophic state in lakes, usually an indicator of shallow and quiet water bodies, and freshwater influence). Associations of colonial algae with other green algae are often indicators of lowland eutrophic lakes (Shumilovskikh et al., 2021; Figure 19).

A summary of the botanical affinities and paleoenvironment conditions for some fungi and algal species present at the AAS is in Table 6.

**Table 6.** Fungal and algal palynomorphs found in the AAS sections, their botanical affinities, and paleoecological habitat. Based on (a) Alvin (1982); (b) Armstrong (n.d.); (c) the Gymnosperm Database (n.d.); (d) Pittermann et al. (2012); (e) Basic Biology (2015); (f) Akyuz et al. (2016); (g) Boufford and Sytsma (2017); (h) Kew Royal Botanical Gardens (n.d.); (i) Rustad (2013); (j) Gary et al. (2009); (k) Tschudy et al. (1984); (l) Barron (2015); (q) Guiry and Guiry (2023); (r) John and Tsarenko (2002); (s) Microbiological Society.

| Taxa | Botanical Affinities | Paleoenvironment |
| --- | --- | --- |
| Fungi |  | Fungi can be found in any habitat, but most live on land, mainly in soil or on plant material (s). |
| *Ovoidites* (Potonié) Thompson & Pflug, 1953 emend. Krutzsch, 1959 | Superdivision Chlorophyta Pascher, 1914<br>Division Chlorophyceae Kutzing, 1834<br>Subdivision Zygnematales Borge & Pascher, 1931. | *Spirogyra* is common in shallow ponds, ditches, and vegetation at large lakes' edges, generally growing free-floating. It can also grow in short-lived ponds during wet weather.<br>*Zygonium* is amphibious or |

Table 6. Fungal and algal palynomorphs found in the AAS sections, their botanical affinities, and paleoecological habitat.
|  |  |  |
| --- | --- | --- |
|  | Zygospores of many species of *Spirogyra* and *Zygogonium* (Zippi, 1998). | terrestrial, often on acid substrates and in moderately warm thermal springs (q). |
| *Schizosporis reticulatus* Cookson & Dettmann, 1959 emend. Pierce, 1976 | Superdivision Chlorophyta Pascher, 1914<br>Division Chlorophyceae Kutzing, 1834<br>Subdivision Zygnematales Borge & Pascher, 1931. | Zygospores of the Zygnemataceae (?). Zygnemataceae are freshwater or subaerial cosmopolitan, living in different habitats. |
| Zygnemataceae spores | Superdivision Chlorophyta Pascher, 1914<br>Division Chlorophyceae Kutzing, 1834. | An excellent indicator of a freshwater realm, floodplain ponds, or lake-fill deposits (f). |
| *Botryococcus* sp. | *Botryococcus braunii*. Superdivision Chlorophyta Pascher, 1914. | Freshwater species are widely distributed in many habitats, e.g., ditches, bog pools, ponds, and lakes (r in q). |
| *Pediastrum* spp. | *Pediastrum duplex* | Freshwater species (q). |

### PALEOECOLOGY OF MARINE PALYNOMORPHS

At the AAS, marine palynomorphs are present; if *in situ*, they are a good indication of marine conditions. Nonetheless, due to the observed assemblage’s characteristics, the conditions seem to be shallow marine, mainly inner platform near the coast.

### DINOFLAGELLATE CYSTS

Dinoflagellate cyst diversity is related to the amount of environmental stress. Environmental stress decreases, and the richness of the dinoflagellate cyst assemblages increases from near-shore to distal-shore (e.g., Dodsworth, 2016; Olde et al., 2015b).

On the other hand, a specimen’s abundance may indicate a bloom of individual species that thrive in specific conditions. Figures 20 and 21 show the characteristics of the dinoflagellate assemblages in the section AASP 3–4 in relation to the total abundance of Peridiniaceae vs. Gonylacaceae, as well as the diversity and richness of the assemblages.

**Figure 20.**
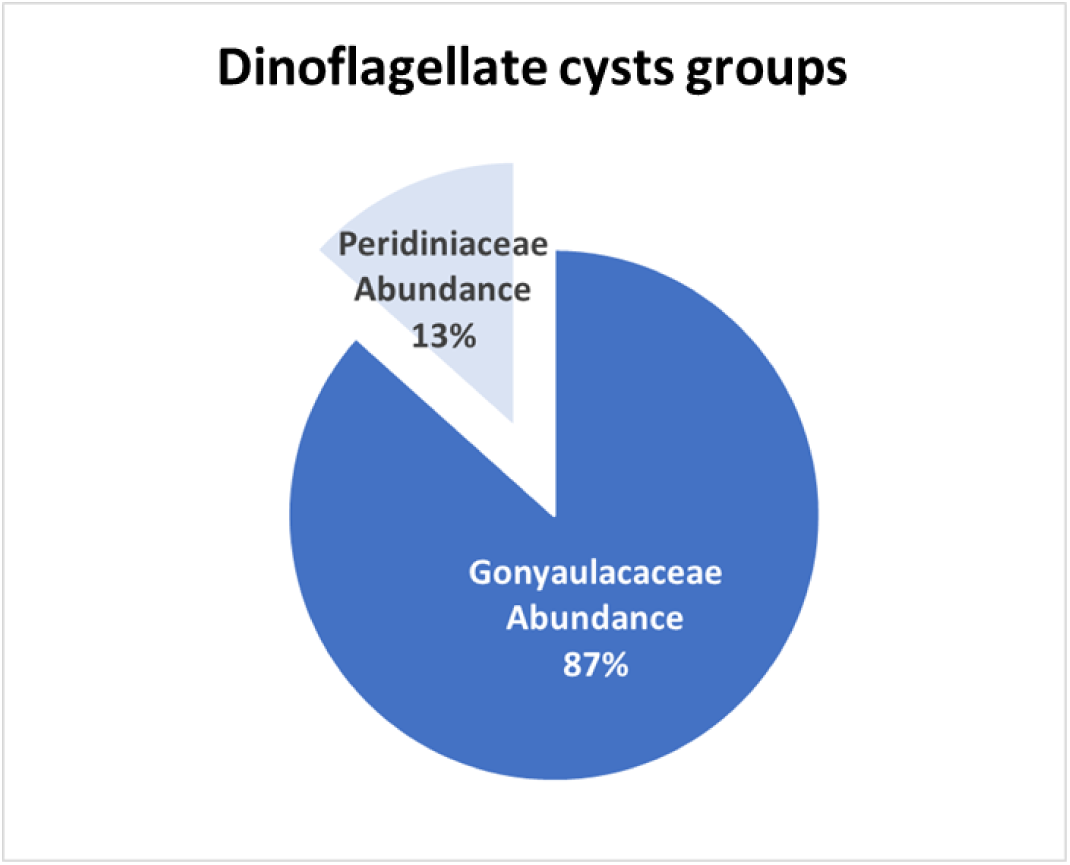
Relative abundance of the two major groups of dinoflagellate cysts present in AASP 3–4.

**Figure 21.**
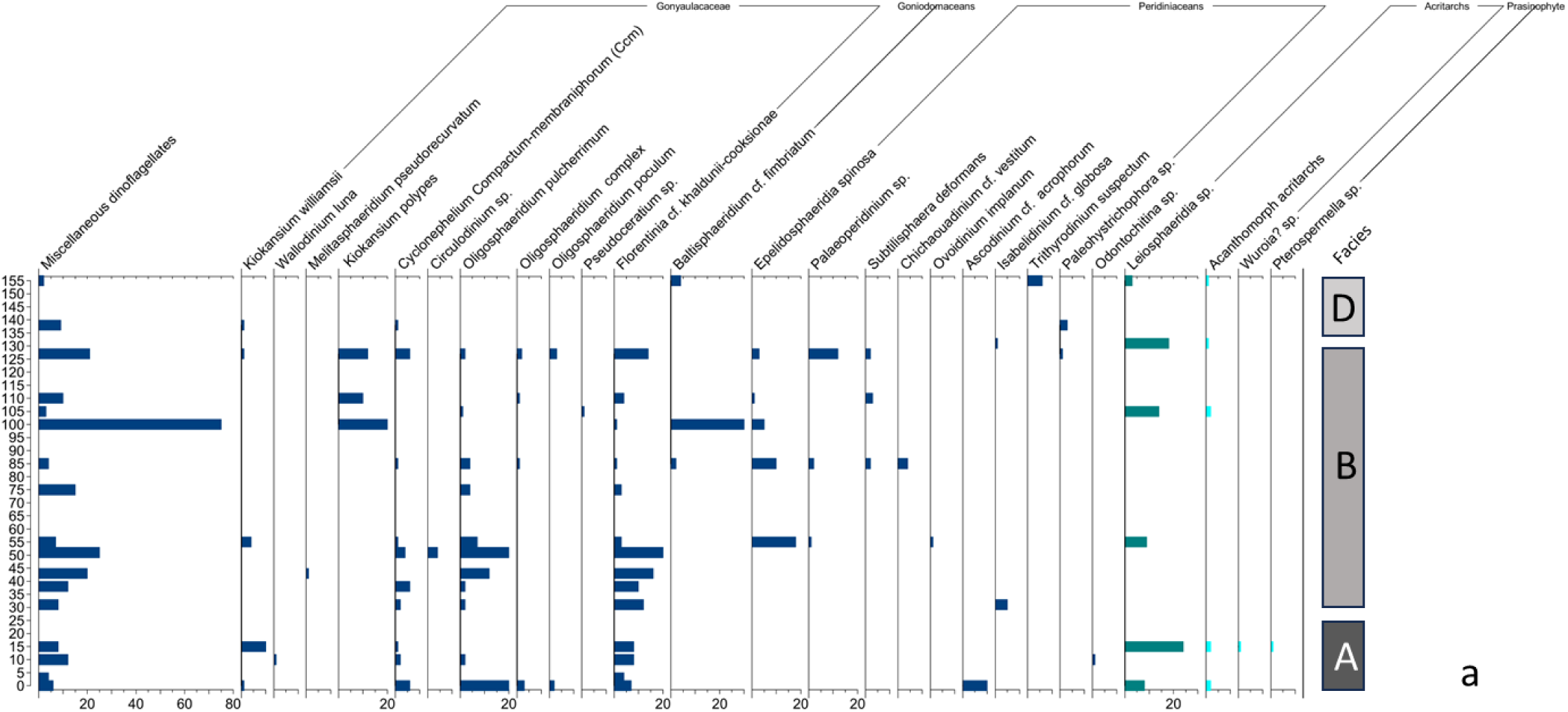
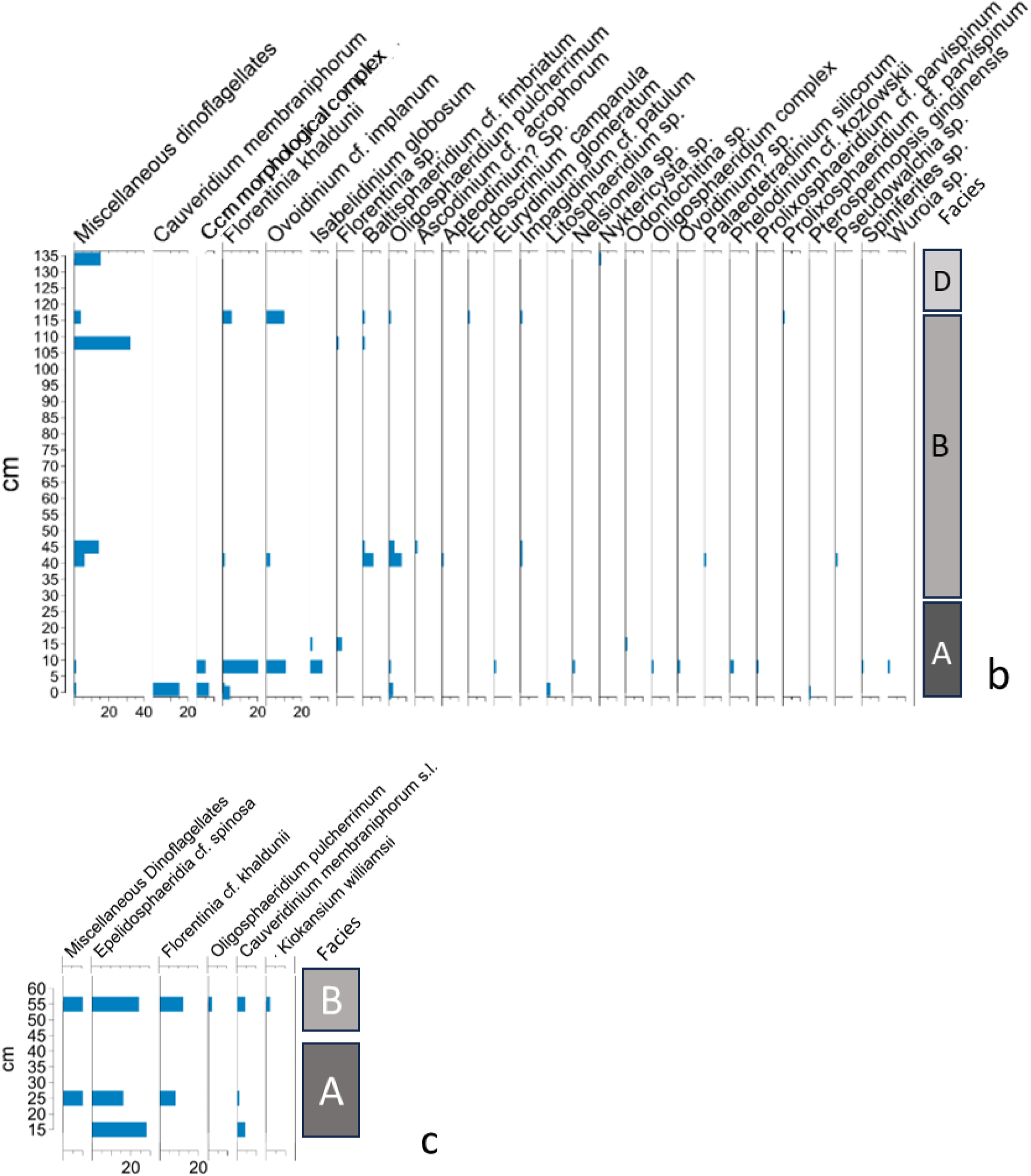
(a) Dinoflagellate cysts, Acritarch, and Prasinophycean distribution chart, showing richness and specimen abundance in AAS 3–4. (b) Dinoflagellate cysts distribution chart section AASP 1-2; (c) Dinoflagellate cysts distribution chart section AASP 6-7.

Dinoflagellate cysts are present throughout the section in most samples. However, each sample’s absolute number of individual cysts is always much less than the total number of sporomorphs. Twenty-two species were identified, and the most numerous group belongs to the gonyaulacaceans (Figures 20 and 21), which include the species *Kiokansium williamsii* (Gonyaulacaceae—uncertain), *Wallodinium luna* (Gonyaulacaceae—uncertain), *Melitasphaeridium pseudorecurvatum* (Gonyaulacaceae—uncertain), *Bacchidinium polypes* now *Kiokansium polypes* (Gonyaulacaceae—uncertain), *Cauveridinium membraniphorum* s.l. (Gonyaulacaceae–Areoligeraceae), *Circulodinium* sp. (Gonyaulacaceae–Areoligeraceae), *Cyclonephelium* sp. (Gonyaulacaceae– Areoligeraceae), *Cauveridinium* sp. (Gonyaulacaceae–Areoligeraceae), *Oligosphaeridium pulcherrimum* (Gonyaulacaceae—Leptodinioideae), *Oligosphaeridium complex* (Gonyaulacaceae—Leptodinioideae), *Oligosphaeridium poculum* (Gonyaulacaceae— Leptodinioideae), *Pseudoceratium* sp. (Gonyaulacaceae—Ceratiaceae), *Odontochitina* sp. (Gonyaulacaceae—Ceratiaceae), *Florentinia? torulosa* (Gonyaulacaceae— Cribroperidinioideae), *Florentinia khaldunii* (Gonyaulacaceae—Cribroperidinioideae).

Also present are *Baltisphaeridium* cf. *fimbriatum* (Goniodomaceae—Pyrodinioideae) and *Baltisphaeridium* cf. *fimbriatum* (Goniodomaceae—Pyrodinioideae, now in the *Filisphaeridium* genus). The paleoecological conditions for some of these dinoflagellate cysts are summarized in Table 7.

**Table 7.** Paleoecological conditions required by some species of dinoflagellate cysts found at the Arlington Archosaur Site, following Harris and Tocher’s (2003).

|  | Euryhaline | Stenohaline | Offshore |
| --- | --- | --- | --- |
| Restricted | *Kiokasium williamsii*<br>(Gonyaulacaceae). |  |  |

|  |  |  |  |
| --- | --- | --- | --- |
| Abundant, but not restricted | *Florentinia* spp.<br>(Gonyaulacaceae—Cribroperidinioideae). |  | *Cyclonephelium membraniphorum*<br>(Gonyaulacaceae—Areoligeraceae). |
|  | *Cyclonephelium vannophorum*<br>(Gonyaulacaceae—Areoligeraceae). |  |  |
|  | *Oligosphaeridium pulcherrimum*<br>(Gonyaulacaceae—Leptodinioideae). |  |  |
|  | *Oligosphaeridium* spp.<br>(Gonyaulacaceae—Leptodinioideae). |  |  |

According to Harris and Tocher (2003), in the assemblages from the Western Interior Sea, it is possible, using cluster analysis, to differentiate three paleoecological groups of dinoflagellate cysts based on their tolerance of salinity variations:

- Euryhaline, tolerating or perhaps preferring lowered salinities. Some species in this group are, among others, *Cyclonephelium brevispinatum*, *Cyclonephelium vannophorum*, and *Oligosphaeridium pulcherrimum*.
- Stenohaline, preferring normal marine salinities. Species that cluster together per Harris and Tocher (2003) are, among others, *Oligosphaeridium totum* and *Spiniferites ramosus reticulatus*.
- Offshore, only tolerant of stenohaline conditions. Species in these clusters include

*Cyclonephelium membraniphorum,* among others.

Most of the species found in the AAS support euryhaline conditions, with tolerance to lowered salinity conditions (Table 7).

The other major group of dinoflagellate cysts present is the Peridiniaceans, which in this section are represented by *Epelidosphaeridia spinosa* (Peridiniaceans—Ovoidinioideae), *Palaeoperidinium* sp. (Peridiniaceans—Palaeoperidinioideae), *Subtilisphaera deformans* (Peridiniacean—Palaeoperidinioideae), *Chichaouadinium* cf. *vestitum* (Peridiniaceans— Palaeoperidinioideae), *Ovoidinium implanum* (Peridiniaceans—Ovoidinioideae), *Ascodinium* cf. *acrophorum* (Peridiniaceans—Ovoidinioideae), *Isabelidinium* cf. *globosum* (Peridiniaceans—Deflandreoideae), *Trithyrodinium suspectum*–*Fromea amphora* (Peridiniaceans—Deflandreoideae), and *Paleohystrichophora* sp. (Peridiniaceae).

The succession of the associations of dinoflagellate cysts in the AASP 3-4 section in Figure 22 shows that the major abundance peaks are related to gonyaulaceans. Those peaks are often closely associated with smaller peaks of peridiniaceans occurring immediately before or after the main increases in gonyaulaceans.

**Figure 22.**
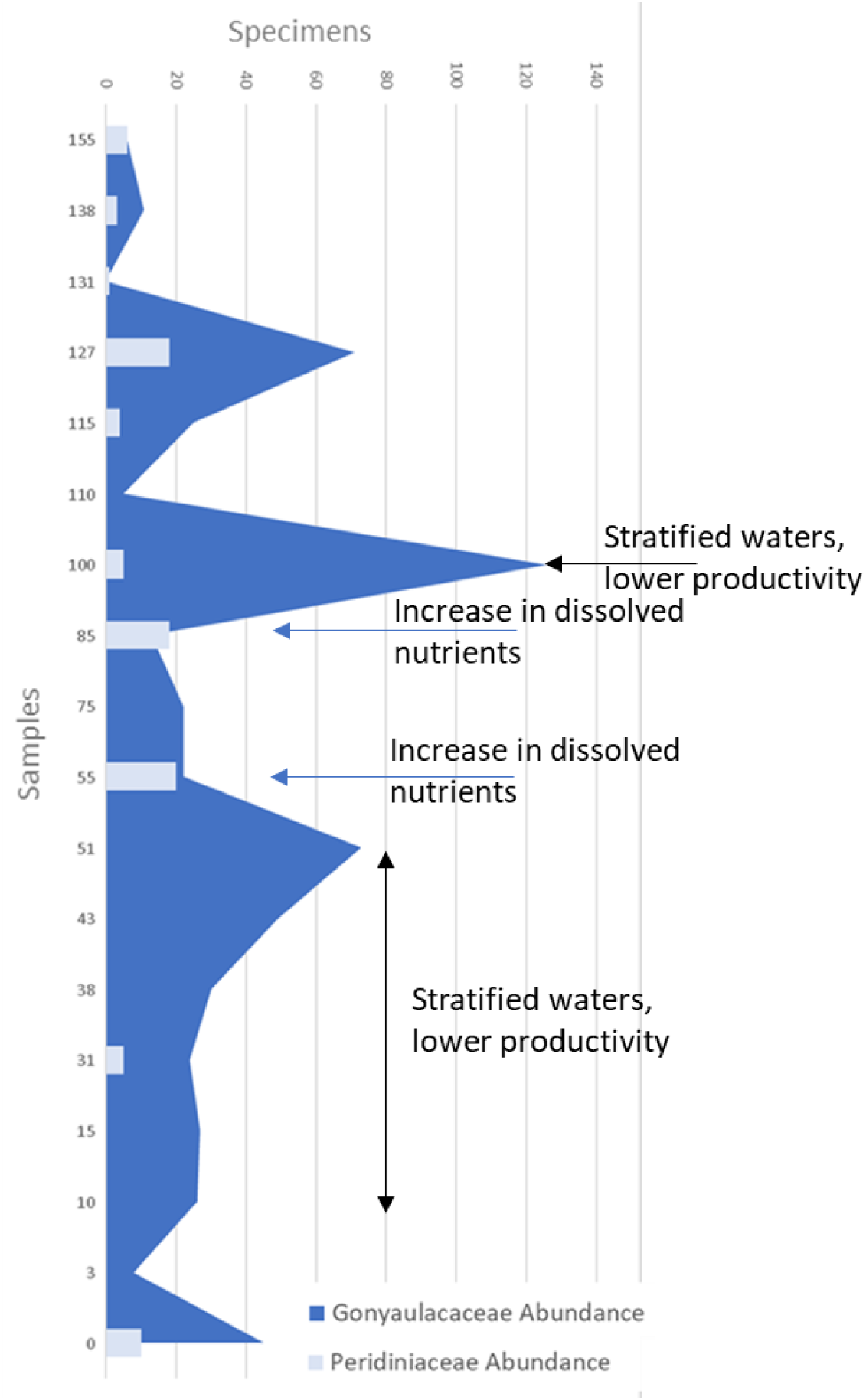
Changes in the abundance of specimens in the two main dinoflagellate cysts groups throughout AASP 3–4, and possible changes in relation with the amount of nutrients dissolved in the water and the productivity.

For the rest of this discussion, we want to highlight that we follow the proposal by Marshall and Batten (1988) and van Helmond et al. (2016) to use the Ccm morphological plexus for the species *Cauveridinium membraniphorum* and *Cyclonephelium compactum* and all the transitional forms, rather than separating them, due to the almost impossible task of dealing with the intricate variation of forms among them. Therefore, this paper oftentimes refer to *Cyclonephelium compactum–membraniphorum* as the Ccm morphological plexus.

### ACRITARCHS

According to Harris and Tocher (2003), many species of acritarchs in the assemblages from the Western Interior Sea were largely restricted to shallow water with reduced salinity. In the section AASP 3–4, acanthomorph acritarchs are present, but low in abundance and diversity, and only in a handful of samples (Figures 21a). Besides, the low abundance of specimens occurs mainly in intervals just after the shallow marine dinoflagellate cysts’ spikes and just before the freshwater colonial algal spikes. This coincides with the observations of Harris and Tocher (2003).

### PALYNOLOGICAL PALEOENVIRONMENTAL INTERPRETATION

We adopted a Source-to-Sink (S2S) analysis approach for the paleoenvironmental interpretation. The model used (Figure 23) shows the complex dynamic of palynomorph transport and redeposition at the AAS.

**Figure 23.**
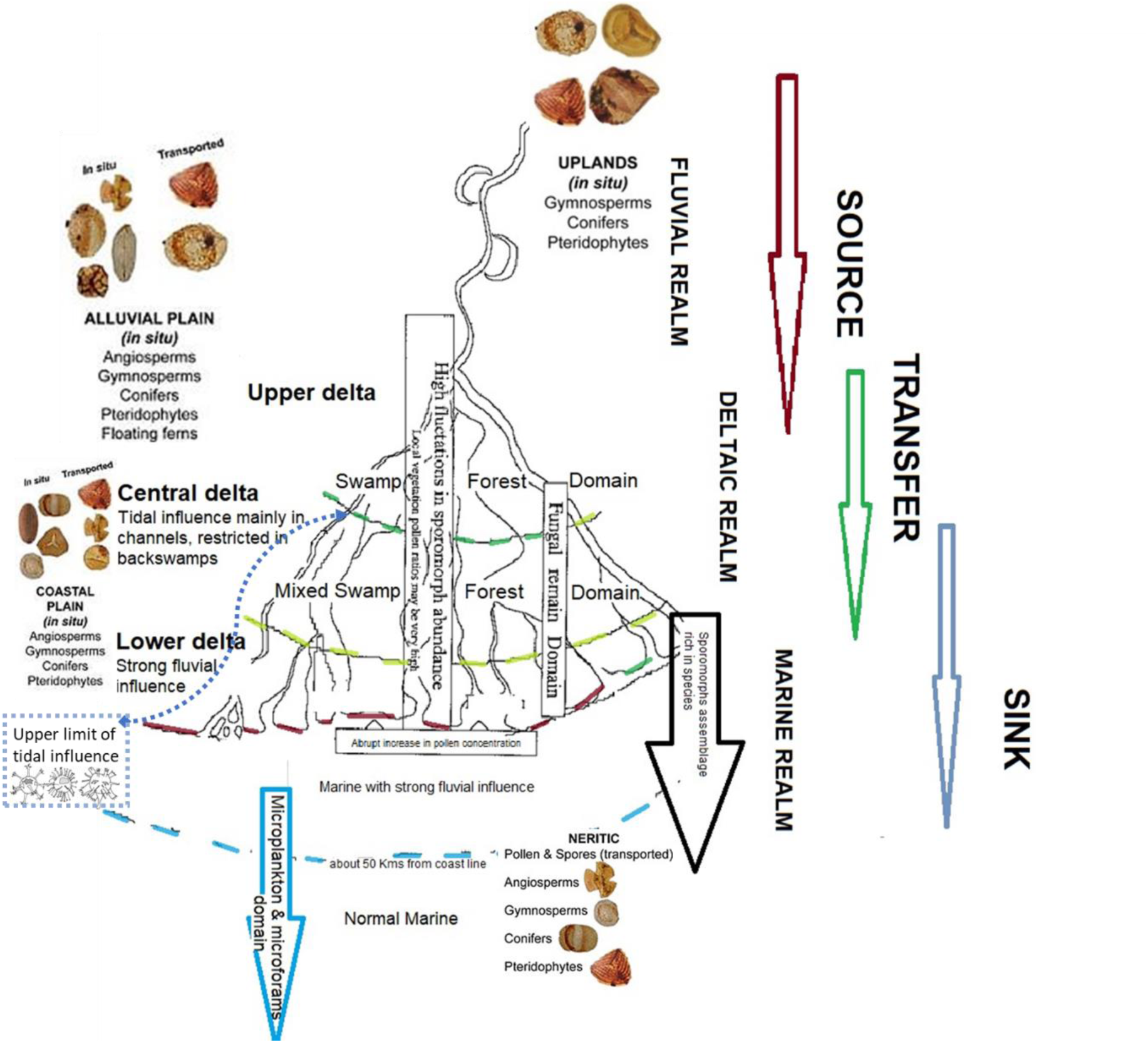
Schematic representation of a Source-to-Sink (S2S) system, in an idealized model representing the interpretation of paleoenvironments at the AAS and the complex dynamic of palynomorphs transport and redeposition. The sources for this model are the author’s own experience and different studies (e.g., Barron, 2015; Brinkhouse, 1994; Harris & Tocher, 2003; Heimhofer et al., 2018; Lorente, 1986; Muller, 1959; Poumot, 1986; Sluijse et al., 2005; Tshudy, 1969).

The palynological elements produced upstream were probably transported by running water and wind currents into the sink area (Figure 23). In purely terrestrial environments favorable for sedimentation, like lakes and ponds, the assemblage would contain in situ sporomorphs mixed with freshwater palynomorphs, but no evidence exists of these exclusively terrestrial environments in the AAS assemblages.

The abundance of megaspores is an indicator of proximity to active terrestrial sources, being abundant in fluvial, marsh, lagoon, and proximal marine facies, with abundance decreasing with the distance to the parent plant (e.g., Speelman & Hills, 1980; Winslow, 1962). The AAS sections show the presence of megaspores of *Goshisphira* sp. from fresh water in assemblages otherwise dominated by other pteridophytes and conifers from upland areas, together with angiosperms, as well as other palynomorphs from fresh water, all mixed up with a dinoflagellate cysts assemblage moderately rich in species, signaling tidal and shallow marine influence, in what otherwise would be terrestrial type associations.

In tidal flats and other coastal tidal-influenced environment, due to the tidal currents bidirectional transport effect, dinoflagellate cysts are oftentimes mixed in the assemblages, although terrestrial palynomorphs dominate. In the nearshore, terrestrial sporomorphs are abundant and mixed with some dinoflagellate cysts species that support salinity variations, including reduced salinities. In the inner neritic, dinoflagellate cysts, saccate pollen, and small sporomorphs will tend to increase in diversity and abundance. Dinoflagellate cyst diversity and abundance increases from near-shore to the distal-shelf, and specimen’s abundance picks may indicate a bloom of individual species that thrive in specific conditions, e.g., red tides.

In the AASP sections, a few rich dinoflagellate cyst assemblages are mixed up with bisaccate spores and smaller angiosperm grains, pointing out the presence of inner neritic shallow marine conditions in some parts of the section.

Towards open marine and oceanic conditions, the palynological assemblages would mainly have dinoflagellate cysts, saccate pollen, and very small sporomorphs carried through turbidite currents and reworked by marine currents to the deep ocean floors. In the AAS, this type of assemblage has not been identified.

An example of the complexity of signals along the AASP 3-4 section is illustrated in Figures 24 and 25.

**Figure 24.**
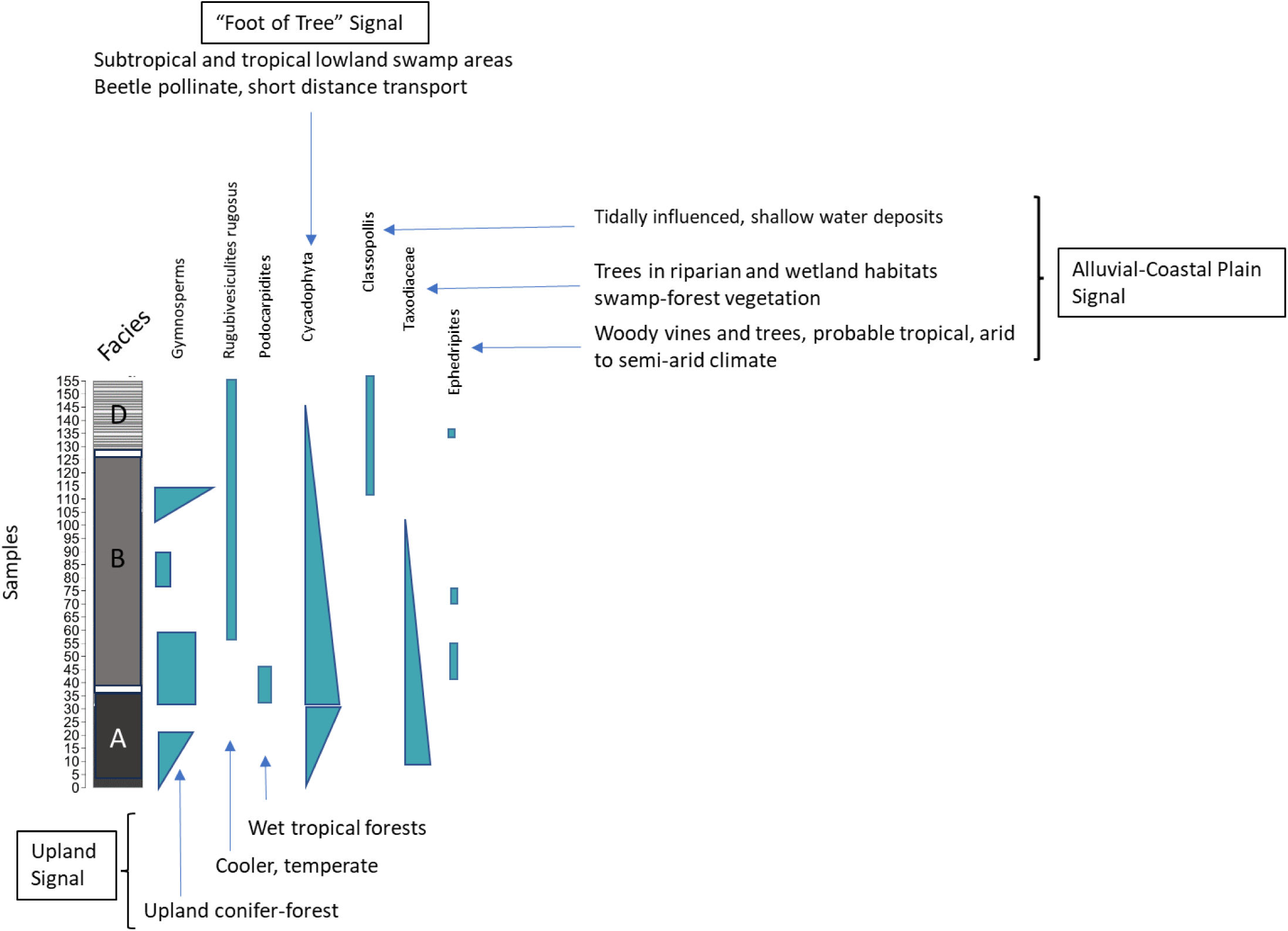
Source and transfer zones’ signals in the AAS assemblage along the AASP 3-4 section. It is possible to observe that upstream of the sink area, the vegetation indicates the presence of tropical to subtropical swamp areas, wetlands, and swamp forests from alluvial coastal plains, while in the sink area, tidally influenced areas are present. Signals from at least three drainage basin areas are in the gymnosperms’ assemblage. The upland signal indicates some conifer-forest cover during the sedimentation of the lower part of Noto’s Facies B, with tropical temperatures and high humidity, but with conditions changing from the middle part of Noto’s Facies B to cooler temperatures. The alluvial and coastal plain signal points to wetlands and swamp forest vegetation, especially during the sedimentation of Noto’s Facies A and B. At Noto’s Facies D, tidally influenced shallow water areas were present. The presence of Cycadophyta, plants associated with beetle pollination with limited pollen dispersion, probably represents “foot of tree” conditions, pointing to sedimentation in or very near widespread tropical lowland swamp areas.

**Figure 25.**
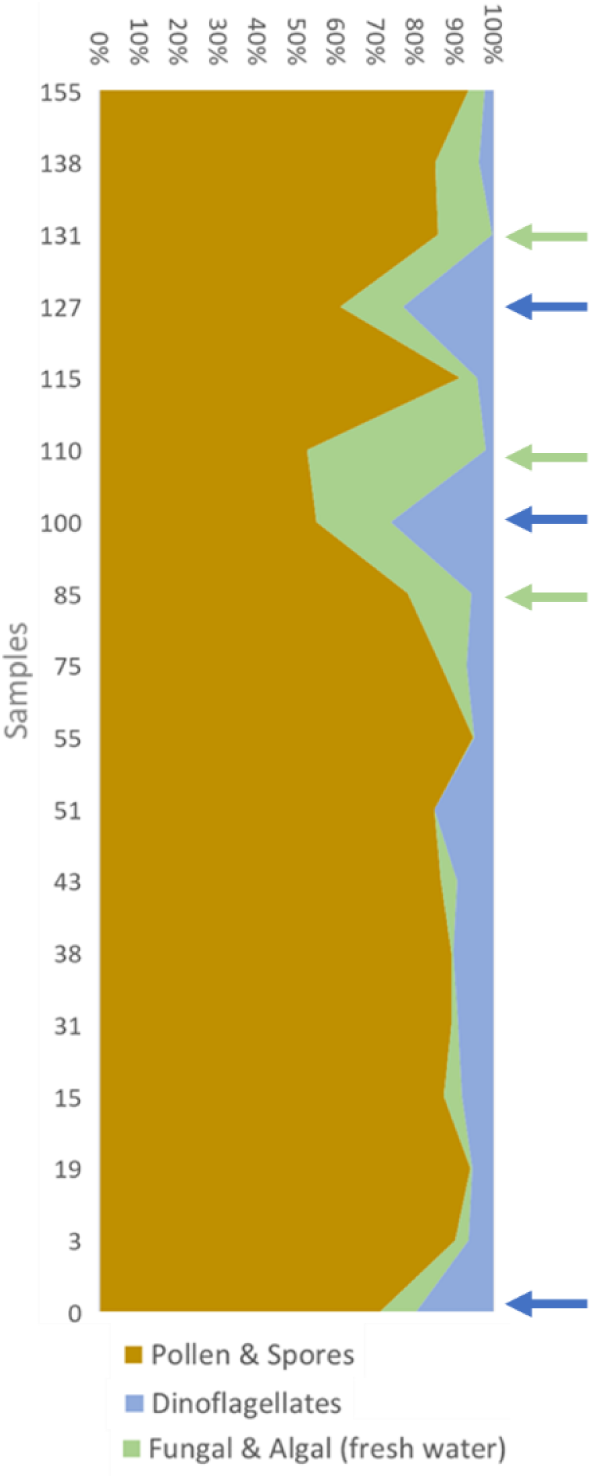
Percentage composition of the palynomorph assemblage. Pollen and spores from terrestrial plants are dominant throughout the section. Still, dinoflagellate cysts are present throughout the section, showing at least three population increases, indicated by blue arrows, probably representing the maximum marine water influence. Those peaks are often closely associated with smaller peaks of fresh water algae and fungal remains occurring immediately before or after the main increases in dinoflagellate cysts, indicated by green arrows.

The major paleoenvironmental groups, organized from terrestrial and freshwater to brackish and marine waters, are displayed in Figure 26. There, it is possible to see the constant and abundant presence of terrestrial palynomorphs and marine palynomorphs throughout the AASP 3-4 section. The marine and brackish palynomorphs represent the signal from the sink, while the terrestrial and fresh water palynomorphs represent the signal from the source and transfer zones of the S2S system.

**Figure 26.**
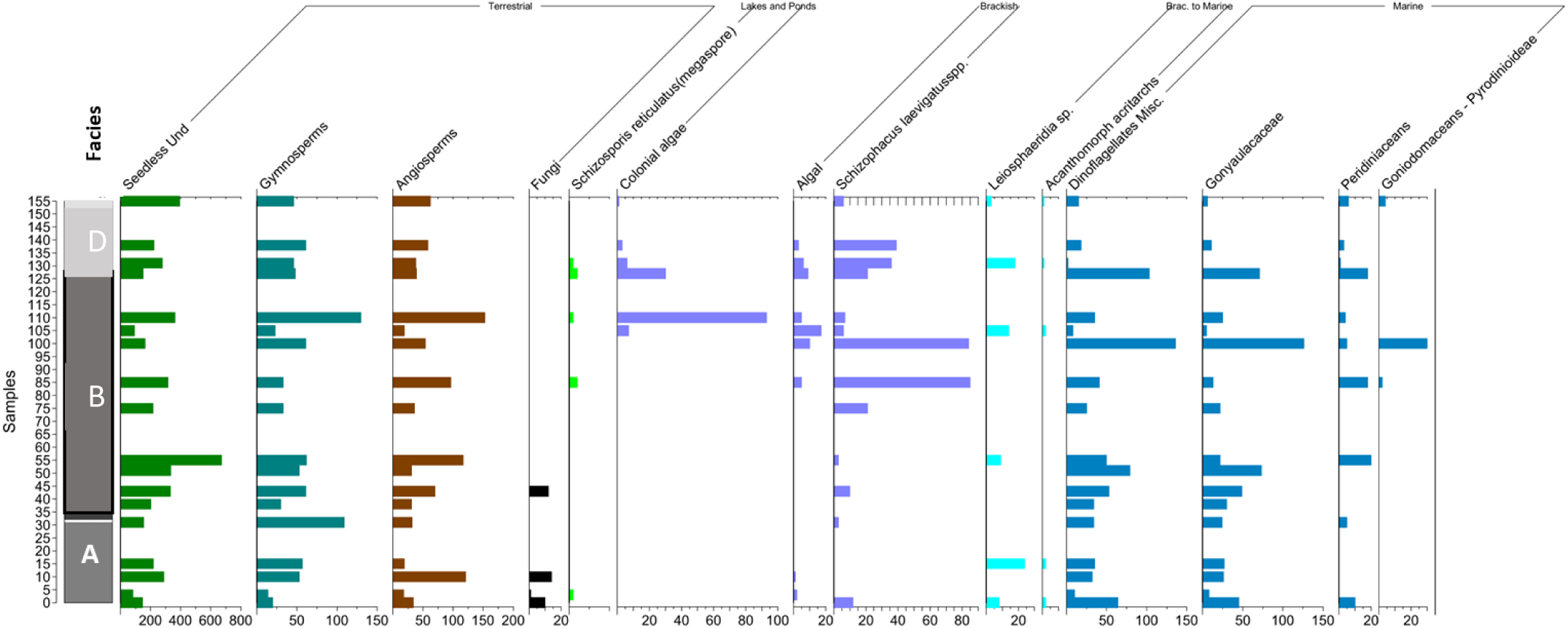
AASP 3–4. Distribution histogram of the major paleoenvironmental groups, organized from terrestrial and freshwater to brackish and marine waters. This graph highlights the multisource characteristic of the palynological assemblages. More upstream elements, representing the source and transfer of the S2S system, with a higher probability of downstream transport, are to the left, while elements typical of the basinal (sink) area are plotted to the right.

The diversity and abundance of dinoflagellate cysts in the assemblages show several levels in which the increase of marine influence is conspicuous. The general characteristics of the assemblage tend to indicate marine but with lowered salinity conditions, as Harris and Tocher (2003) described based on their cluster analysis of samples from elsewhere in the Western Interior Sea. The levels with the aforementioned increased diversity and abundance of the dinoflagellate cyst assemblages are proposed here as possible flooding surfaces (Figure 27).

**Figure 27.**
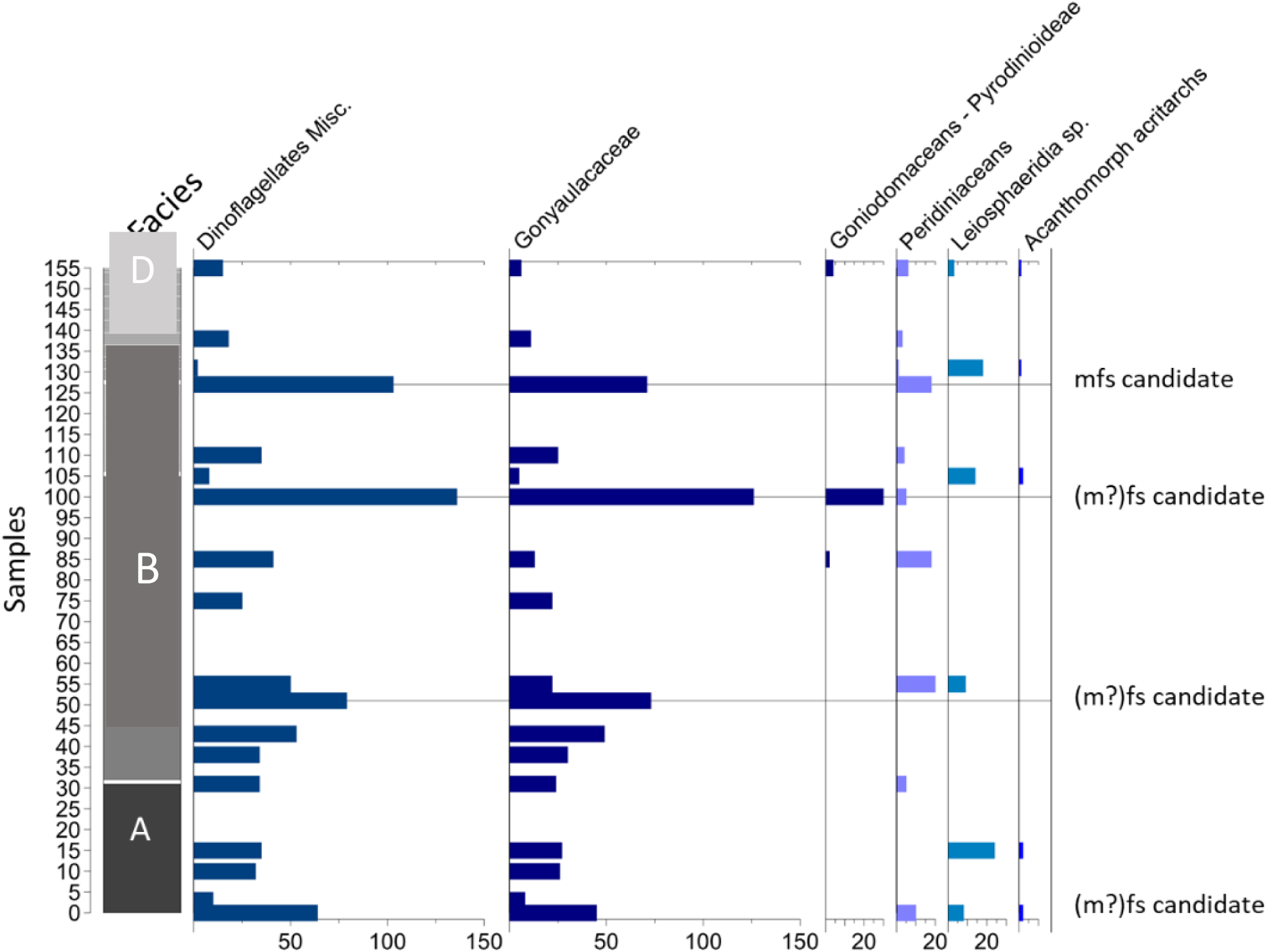
Histogram showing the variations in abundance of the dinoflagellate cysts, acritarchs, and Leiosphaeridia assemblages throughout the section. The levels with the highest abundance of dinoflagellate cysts are indicated in the graphs as (maximum?) flooding surface candidates.

In the AAS assemblages, the presence of Ccm morphological complex may signal the boreal dinocyst migration into the AAS area. The migration of this taxon is mentioned in the literature (van Helmond et al., 2016) as one of the dinoflagellate cysts taxa that migrated south during the Plenus Cold Event (PCE; Eldrett et al., 2017). According to Eldrett et al. (2017), “a major re-organization [of the currents’ flow occurred] during the latest Cenomanian–Turonian [95–94 Ma] as a full connection with a northerly boreal water mass was established during peak transgression.”

Figure 28 shows the occurrence of this taxon along the combined AASP1-2-3-4 combined section. It is also present in sections AASP 5 and AASP 6–7.

**Figure 28.**
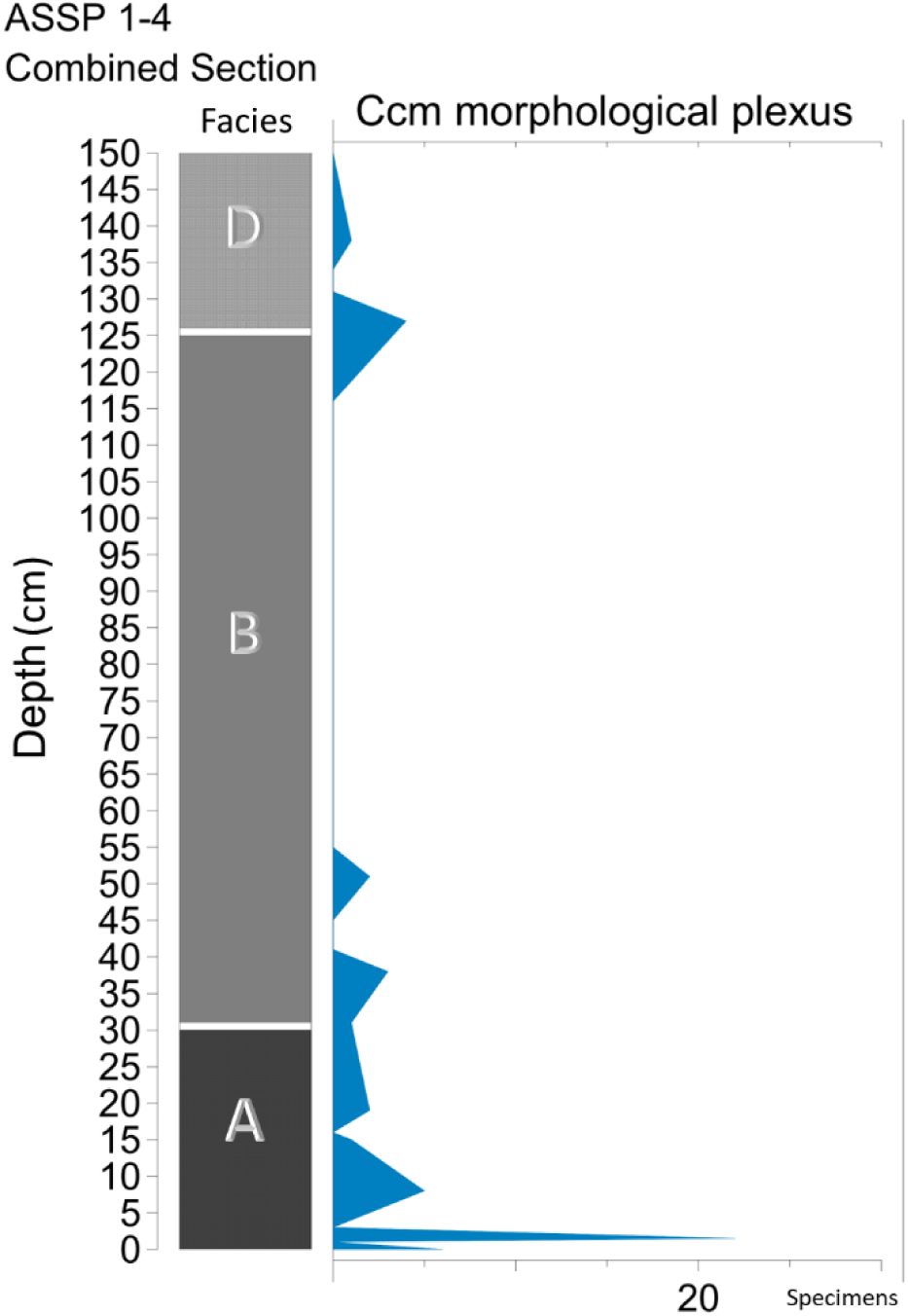
The Ccm morphological plexus is present in almost every sample in the studied AAS sections. This figure shows the abundance of the type in the combined section AASP 1–2-3-4.

Paleoenvironmental-palaeoecological studies are frequently associated with palynostratigraphic studies, but to make this discussion feasible, we will focus on a few of the works that may have relevance to the paleoenvironmental–paleoecological analysis done here.

One of the most important papers for the novelty of the applied “ecogroup” concept is by Abbink et al. (2004), who recognized in the Jurassic of the North Sea several sporomorph ecogroups (SEG), including upland, lowland, river, pioneer, coastal, and tidally influenced SEGs. The SEG communities with modifications were successfully applied in terrestrial coal-bearing series across Hungary’s Triassic/Jurassic boundary (Ruckwied et al., 2008). To apply Abbink’s SEG communities, Ruckwied et al. (2008) modified and redefined the terrestrial SEG communities for various reasons, i.e., the different climatic conditions in their studied section, the unknown relationship of many Mesozoic sporomorph morphospecies with the parent plants, and the unknown tolerance to stress and disturbance of many of the types used, a factor that was not included in Abbink et al.’s (2004) grouping and analysis. Ruckwied et al. (2008) further subdivided and rearranged Abbibk’s lowland SEG communities, including three additional ones: “warmer lowland,” “wetter lowland,” and “dryer lowland.”

Table 8 presents a compilation of the SEG ecogroups’ composition in terms of palynomorphs, as defined by Abbink et al. (2004), and our observations in the AAS assemblages. We finally decided not to applied the SEG groups concept in the sense of Abbink et al. (2001, 2004) because besides the differences in age, paleolatitude, paleoclimatology, it is specially the paleoenvironmental differences between our S2S model and the SEG communities, due to the more or less continue marginal marine signal present through out the AAS sections, but absent in the ecogroups model.

**Table 8.** Summary of Abbink’s SEG vs. AAS key paleoecological markers.

| SEG | Sporomorphs | Abbink et al. (2004) | AAS assemblages (Tables 1–5) |
| --- | --- | --- | --- |
| Upland | Bisaccate pollen grains | No climate inference was given. | Upland<br>*Podocarpidites* sp.: Trees prosper in wet tropical forests.<br>*Rugubivesiculites rugosus*: Cooler, temperate.<br>*Abietineaepollenites* spp.: Cooler, temperate. |
| Lowland | *Cicatricosisporites* spp.,<br>*Concavisporites* spp.,<br>*Cycadopites* spp.<br>*Deltoidospora-Dictyophyllidites*<br>*Cyathidites* spp.<br>*Eucommiidites trøedsonii*<br>*Gleicheniidites* spp.<br>*Ischyosporites* spp.<br>*Monosulcites* spp.<br>*Plicatella* spp. | marsh: “wetter”;<br>“warmer”<br>“drier”; “warmer”<br>“drier”; “warmer”<br>“drier”; “warmer”<br>“drier”; “warmer”<br>“drier”; “warmer”<br>“wetter”; “warmer”<br>“drier”; “warmer”<br>“drier”; “warmer” | *Cicatricosisporites* spp.: Lowland, tropical and subtropical forest. Warmer, wetter.<br>*Concavisporites* spp.: Lowland, warmer, drier.<br>*Cycadopites* spp.: Subtropical and tropical lowland. Short dispersal.<br>*Cyatheadites australis*: Lowland, wetland; brackish swamp, and back mangrove. Warmer, drier.<br>*Gleicheniidites senonicus*: Coastal. Tropical and temperate.<br>*Monocolpites* spp./<br>*Monosulcites* spp.: Subtropical and tropical lowland swamp areas. |
| River | *Lycopodiacidites* spp. | No climate inference was given. | *Lycopodiumsporites* spp.: Terrestrial cosmopolitan fern, warmer (?). |
| Pioneer | None present | No climate inference was given. |  |
| Coastal | *Araucariacites* spp.<br>*Callialasporites* spp.<br>*Corollina* spp. (*Classopollis* spp.) | “cooler”<br>“cooler”<br>“warmer” | *Classopollis* spp.: Warm habitats (a), arid coastal settings. Could tolerate drought stress and high salinity. It can be found in tidally influenced, shallow-water deposits, and these plants could thus indicate the vicinity of the paleo-shoreline.<br>*Taxodiaceaeapollenites* spp.: trees that are the principal component of swamp-forest vegetation |
| Tidally-influenced | None present | No climate inference was given. | Mainly euryhaline dinoflagellate cyst assemblages. |

### PALEOCLIMATE EVIDENCE

Paleoclimatic evidence is primarily registered in the regional pollen and spore signal from the transported portion of the assemblage. Regional changes in the plant community due to climate changes (temperature and humidity) are reflected in the composition (diversity and richness) of the sporomorph assemblages.

Recently, much attention has been given to the climate as a driver of significant changes occurring during the Cretaceous. Among the latest studies, Heimhofer et al. (2018) discussed the Ocean Anoxic Event 2 (OAE2) in France. According to Heimhofer et al. (2018), in the Casis region (France), the change from the exceptionally warm conditions that prevailed during the Cretaceous thermal maximum to the cooler PCE was characterized by a change in the structural composition of the plant communities. This change is seen as a proliferation of open, savanna-type vegetation rich in angiosperms with a decrease in conifer forests. Those authors proposed two different scenarios for the vegetation during the OAE2, a so-called “mode a” with an exceptionally warm and humid climate and conifer-dominated forests with subordinated angiosperms, and another scenario called “mode b” with a cooler and less humid climate, with the development of open savannah vegetation and increased abundance of angiosperms.

Other authors, e.g., Lowery et al. (2018), proposed that at the onset of the OAE2, boreal waters reached paleolatitudes of 35°N in the western CWIS. Van Helmond et al. (2014) reported a *Cyclonephelium compactum–membraniphorum* bloom at Bass River (New Jersey, slightly north of 30°N, western Atlantic). That boreal dinoflagellate cyst was also recognized in well and outcrop samples from Texas (Shell Iona-1 Core: Eldrett et al., 2014; Lozier Canyon: Dodsworth, 2016), suggesting that the influx of boreal surface waters reached the CWIS southern-central region (paleolatitudes of 30–35°N), in agreement with Eldrett et al.’s (2017, Fig. 9B) reconstruction of the water mass circulation. Falzoni and Petrizzo (2022) proposed other alternatives for the persistent occurrence of *Cyclonephelium compactum–membraniphorum* in Europe and the CWIS. According to those authors, it would mean a persisting supply of boreal surface waters after the PCE or that the boreal dinoflagellate cysts “became seasonal and adapted to lower latitude environments.”

We observed variations in the behavior of these vegetation groups throughout the Facies in AAS. In Facies A, there is a change from an angiosperm-dominated to a conifer-dominated assemblage, signaling a regional climate change affecting the upland drainage area, but also, the dinoflagellate cysts Ccm morphological plexus’s presence allows us to propose that cooler surface waters reached as south as the AAS most probably during the PCE of the OAE2.

Following the vegetation trends proposed by Heimhofer et al. (2018) and the evidence from the Ccm morphological plexus (Figure 28) in the AAS, we suggest that the climate was cooler and probably less humid at the start of Facies A’s and middle part of Facies B sedimentation (Figure 29), but the conditions warmed up towards the top of Facies A, and the lower part of Facies B, allowing the development of conifer-forest communities elsewhere in the drainage basin. In Facies B, the trend seems to reverse to cooler and probably dryer conditions towards the middle part, favoring the growth of open savannah vegetation. No clear dominance of any groups exists at the upper part of Facies B and D. Still, the income of a more diversified angiosperm population that included monocots and dicots may be pointing out the maintenance of somewhat cooler (Figures 18 and 19) conditions, but not as cold as seen at the lowermost part of Facies A and middle part of Facies B.

**Figure 29.**
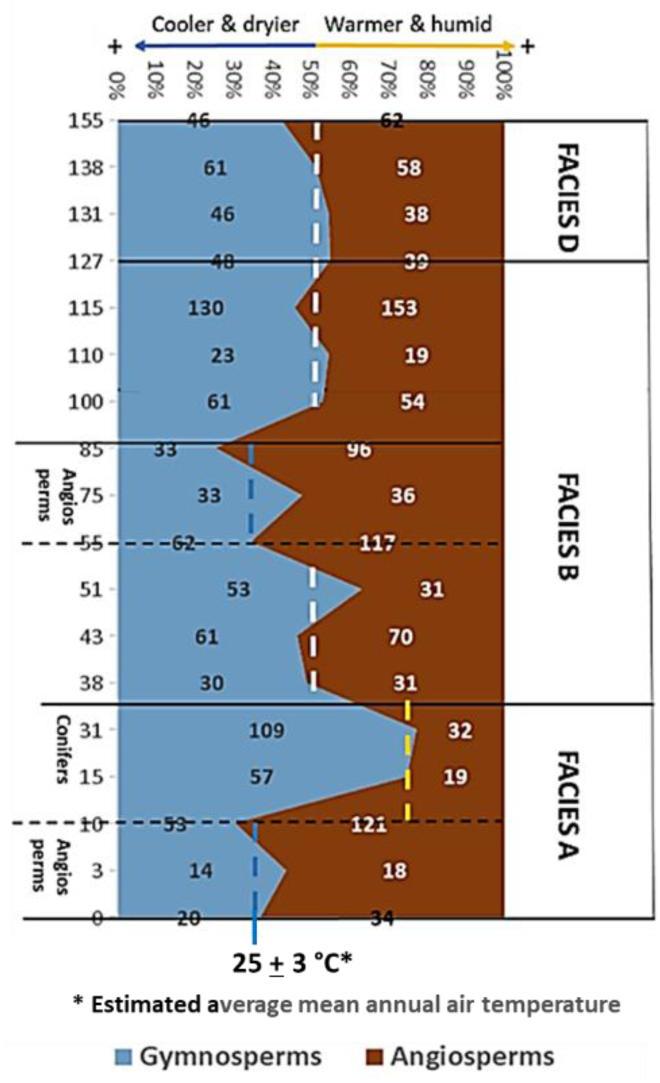
The 100% stacked area graph shows the variation along AASP 3–4 of relative percentage abundance between conifers and angiosperms. Please note that the graph excludes pteridophytes and other seedless plants that otherwise account for more than 50% of the pollen and spore population. The variation in the percentages of conifers vs. angiosperms within the lithological facies follows some trends described in the text. Vertical dotted lines highlight each interval’s climate trend, e.g., the blue dotted line shows a cooler- dryer trend (about 25 ± 3 °C average air temperature), the yellow dotted line shows a warm and more humid trend (29 ±3 °C average air temperature?), while the white dotted line (50%) represents cooler temperatures than the yellow line trend.

Based on pedogenic phyllosilicate analyses from a sample taken in the middle part of Noto’s Facies B (Andrzejewski and Tabor, 2020), the average mean annual soil temperature for the sample’s clay formation was 27 ± 3 °C, which would indicate a “cooler” average mean annual air temperature of about 25 ± 3 °C. That would also be the temperature during the sedimentation of the lower part of Facies A (Figure 29). Although the calculated temperature is warmer than nowadays since the last 100 years historical average mean annual air temperature in North Texas is 19 °C (https://www.weather.gov/fwd/dmotemp), a 25 ± 3 °C temperature would be aligned with what is known about the temperatures during the Aptian-Cenomanian in other parts of north-central Texas and southern Oklahoma that reached up to 29 ±3 °C (Andrzejewski and Tabor, 2020).

### PALYNOSTRATIGRAPHY

Since its discovery, the age and stratigraphic position of the Arlington Archosaur Site outcrop was traditionally assigned to the early Middle Cenomanian based on its lithological characteristics similar to the Lewisville Formation of the Woodbine Group. Nevertheless, several recent revisions of the stratigraphy of Texas’ Late Cretaceous proposed changes to the traditional lithostratigraphic subdivision of the Woodbine and the Eagle Ford units (e.g., Denne et al., 2016), while conflicting stratigraphic evidence for the Lewisville Formation and its equivalents suggests some parts may be as young as the late Cenomanian/early Turonian (Ambrose et al., 2009; Christopher, 1982; Gifford, 2021; Jacobs et al., 2005; Kennedy & Cobban, 1990). Besides, Cloos (2018), in her thesis, suggested an upper Cenomanian– lower Turonian age for the upper HW section 360 due to the presence of triporate pollen of Normapolles affinity (*Pseudoplicapollis* sp.?).

#### Pollen and Spores’ Palynostratigraphy

The sporomorph assemblage at the Arlington Archosaur Site is diverse in species and relatively rich in specimens. At least a dozen pollen and spore taxa identified are typical of the Cenomanian associations of the Western United States and the CWIS.

Although most species have a relatively long range in the Northern Hemisphere, their range is more restricted in the Western Interior Sea (Figure 30). An essential aspect of the AAS section is the angiosperm pollen presence, which agrees with the Cenomanian angiosperm boom registered worldwide.

**Figure 30.**
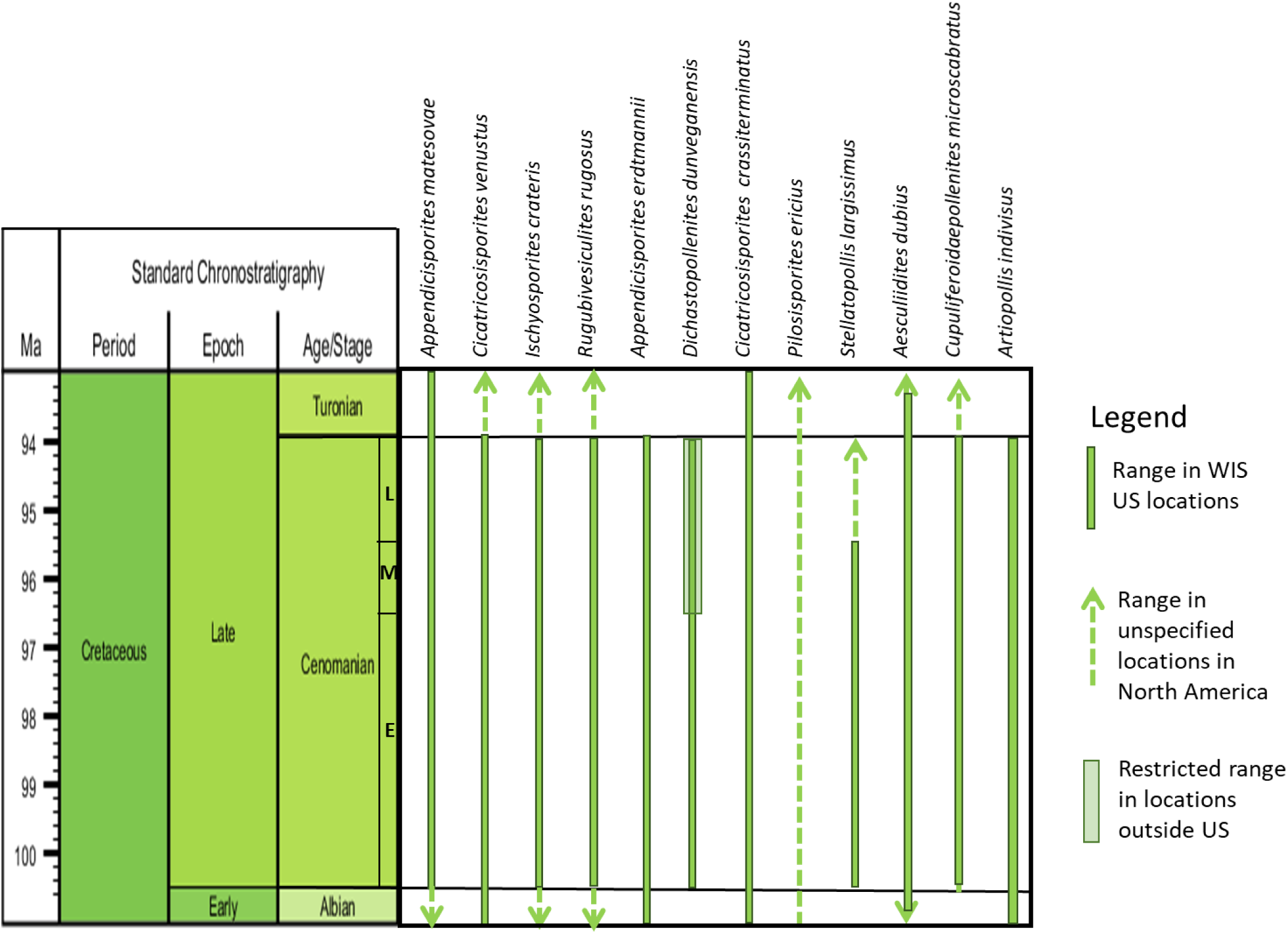
A selection of AAS section key sporomorph species, showing their general stratigraphic range, as is known from locations in the Western United States and the Western Interior Sea, based on Palynodata Inc. & White, J. M. (2008).

The key species identified in the AAS are *Appendicisporites matesovae*, *Cicatricosisporites venustus*, *Ischyosporites crateris*, *Rugubivesiculites rugosus*, *Appendicisporites erdtmannii* (here including forms from very similar morphospecies, e.g*., Appendicisporites auratus* and *Plicatella fucosa*), *Dichastopollenites dunveganensis*, *Cicatricosisporites crassiterminatus*, *Pilosisporites ericius*, *Stellatopollis largissimus*, *Aesculiidites dubius*, *Cupuliferoidaepollenites microscabratus*, and *Artiopollis indivisus.* The abundance of those key species is displayed in Figure 31.

**Figure 31.**
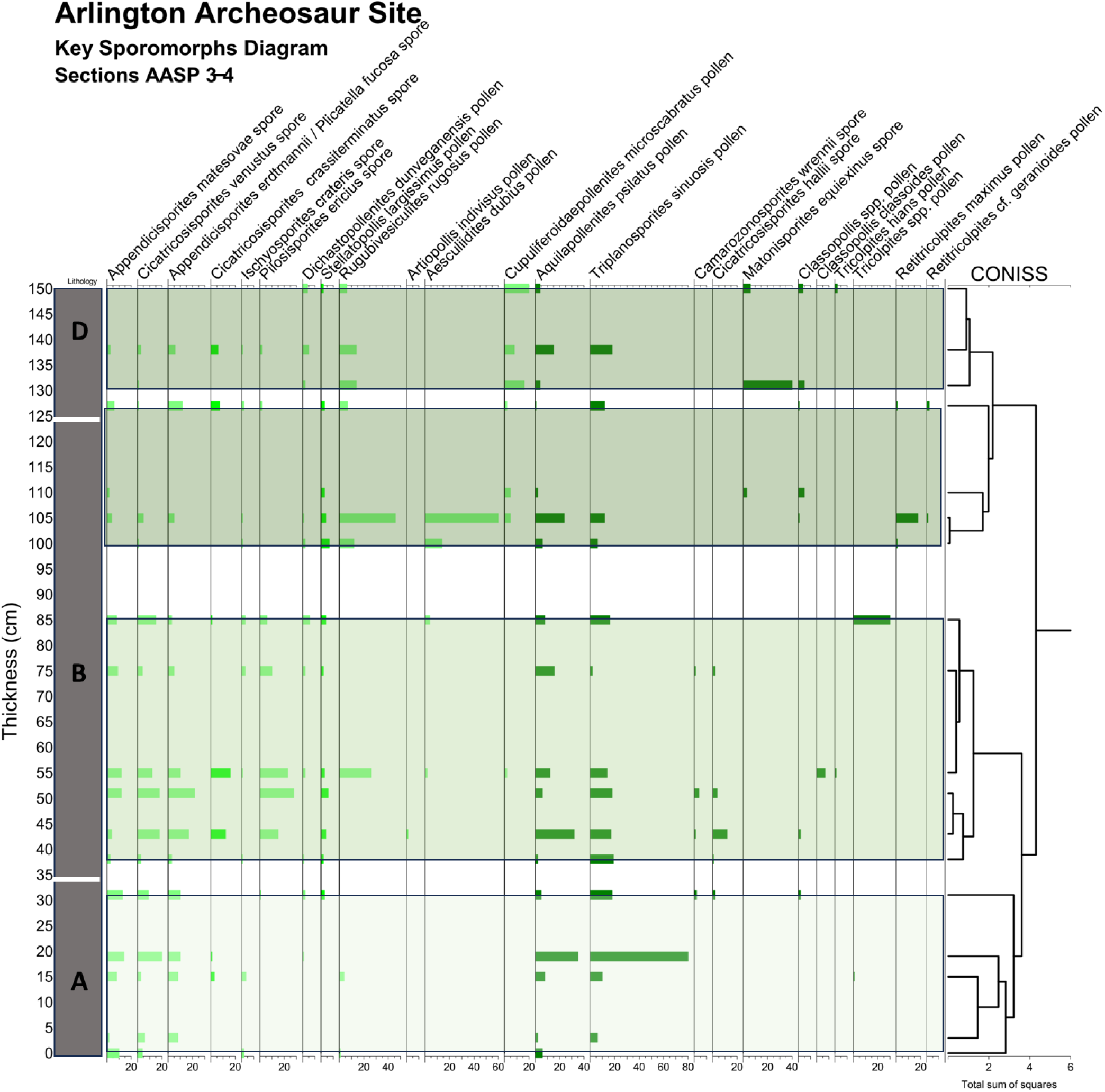
Key sporomorphs diagram. In light green, species with regionally defined tops or bases. In dark green, species with apparent tops or bases in the AASP 3-4 section. The CONISS cluster is displayed to the right of the diagram, highlighted in different shadows of green are the four clusters based on the stratigraphically constrained cluster analysis. In shadow patterns, the main lithological facies, from base to top: Noto’s Facies A, B, C, D. No samples are available from Noto’s Facies C.

Several apparent first and last occurrences are registered in the AAS section, e.g., *Rugubivesiculites rugosus*, an extinct conifer mainly restricted to the Late Albian– Cenomanian of the CWIS, *Dichastopollenites dunveganensis* (Mid-Cenomanian or younger), *Cicatricosisporites crassiterminatus* (Mid-Cenomanian or younger, Dakota Formation), and *Pilosisporites ericius*, all of which have their first apparent occurrence close to the base of Noto’s Facies B. Also, close to the top of Noto’s Facies B, *Artiopollis indivisus* has its apparent base, *A. dubious* has an acme, and immediately afterward, it has its apparent last occurrence. At the same time, *A. indivisus* continues through Noto’s Facies D. *Cupuliferoidaepollenites microscabratus* (Cenomanian, Minnesota) and *Stellatopollis largissimus* (Cenomanian) have their first apparent occurrence (FAO) within Noto’s Facies A and continue throughout the section.

Some species have their last apparent occurrence (LAO) at or close to the base of Noto’s Facies D, e.g., *Cicatricosisporites venustus*, *Ischyosporites crateris*, *Appendicisporites erdtmannii* inc. *Plicatella fucosa*, *Cicatricosisporites crassiterminatus*, *Pilosisporites ericius*, and *Aesculiidites dubius*. Those species’ sudden disappearance may indicate a regional paleoenvironmental change or a discontinuity within Noto’s Facies D.

*Cicatricosisporites crassiterminatus* is one relevant species present throughout the AAS section. Previous authors had reported it from the Woodbine Formation in Oklahoma (Hedlund, 1966), the “Dakota Sandstone” in Arizona, Utah, Iowa, and Nebraska (Agasie, 1969; May & Traverse, 1973; Ravn & Witzke, 1994; Romans, 1972, 1975), and the Lower Tuscaloosa Group in Louisiana and Mississippi (Ravn & Witzke, 1994). When Ravn and Witzke (1994) discussed this species’ occurrence in their samples from Iowa and Nebraska, they said that it “appears to be a reliable indicator of strata no older than Cenomanian, and the available evidence suggests that most occurrences are from middle Cenomanian and younger strata,” leaving open the possibility that the upper part of their Dakota Formation section could be Late Cenomanian. However, they said that they had found no proof of it.

This apparent last occurrence datum of *Cicatricosisporites crassiterminatus* is relevant for the Cenomanian of the CWIS, although recently reported from the Turonian of Utah (Akyuz et al., 2016). This finding reinforces the idea of a slightly younger age, as Ravn and Witzke (1994) suspected for the upper levels of their Iowa and Nebraska sections, hence Woodbine’s correlative Dakota Formation. Other species discussed by Ravn and Witzke (1994) that are also relevant in our section are *Plicatella fucosa* (i.e., *Appendicisporites erdtmannii*) and *Stellatopollis largissimus*.

The main apparent bioevents based on these FAO and LAO are shown in Figure 32.

**Figure 32.**
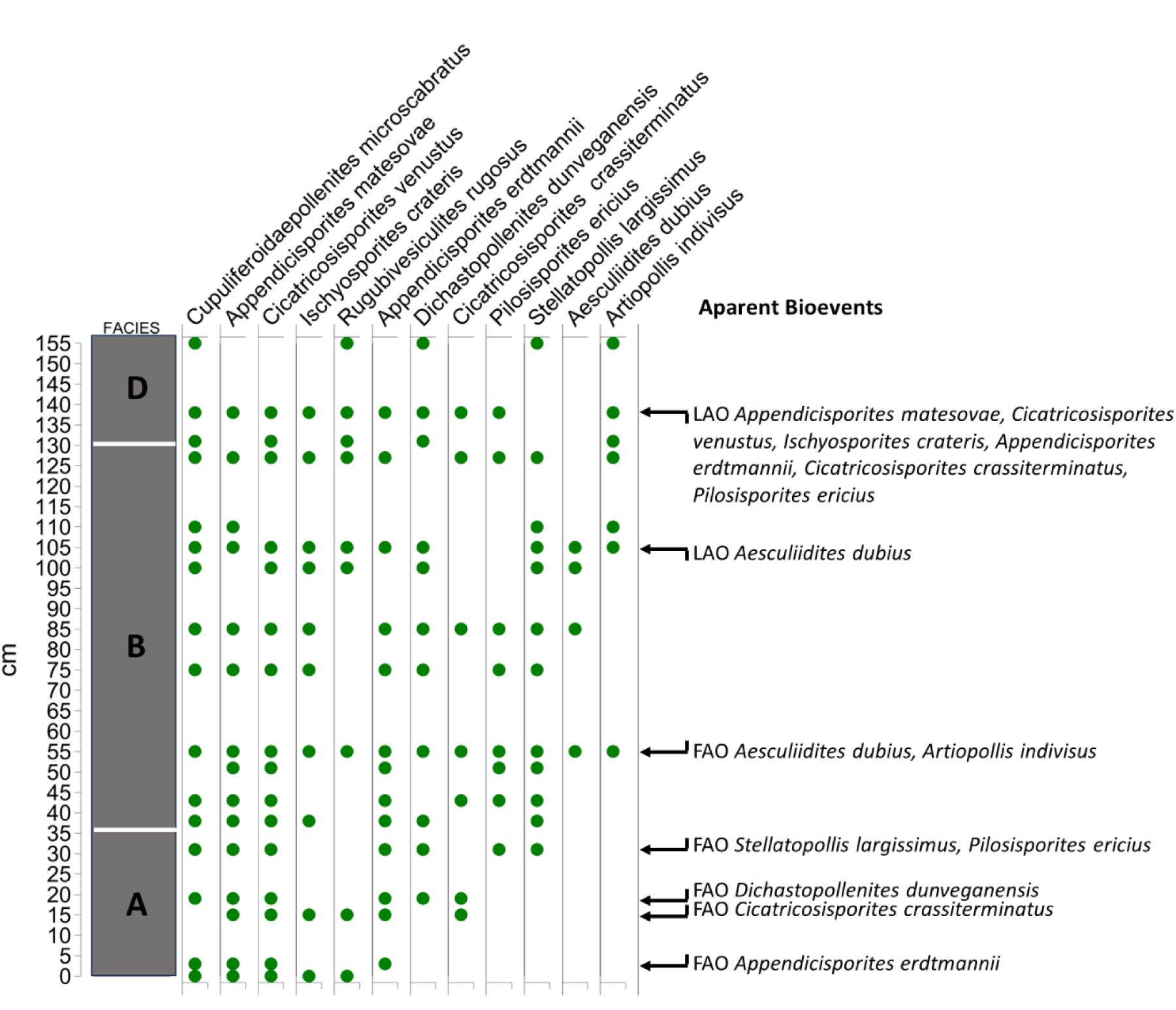
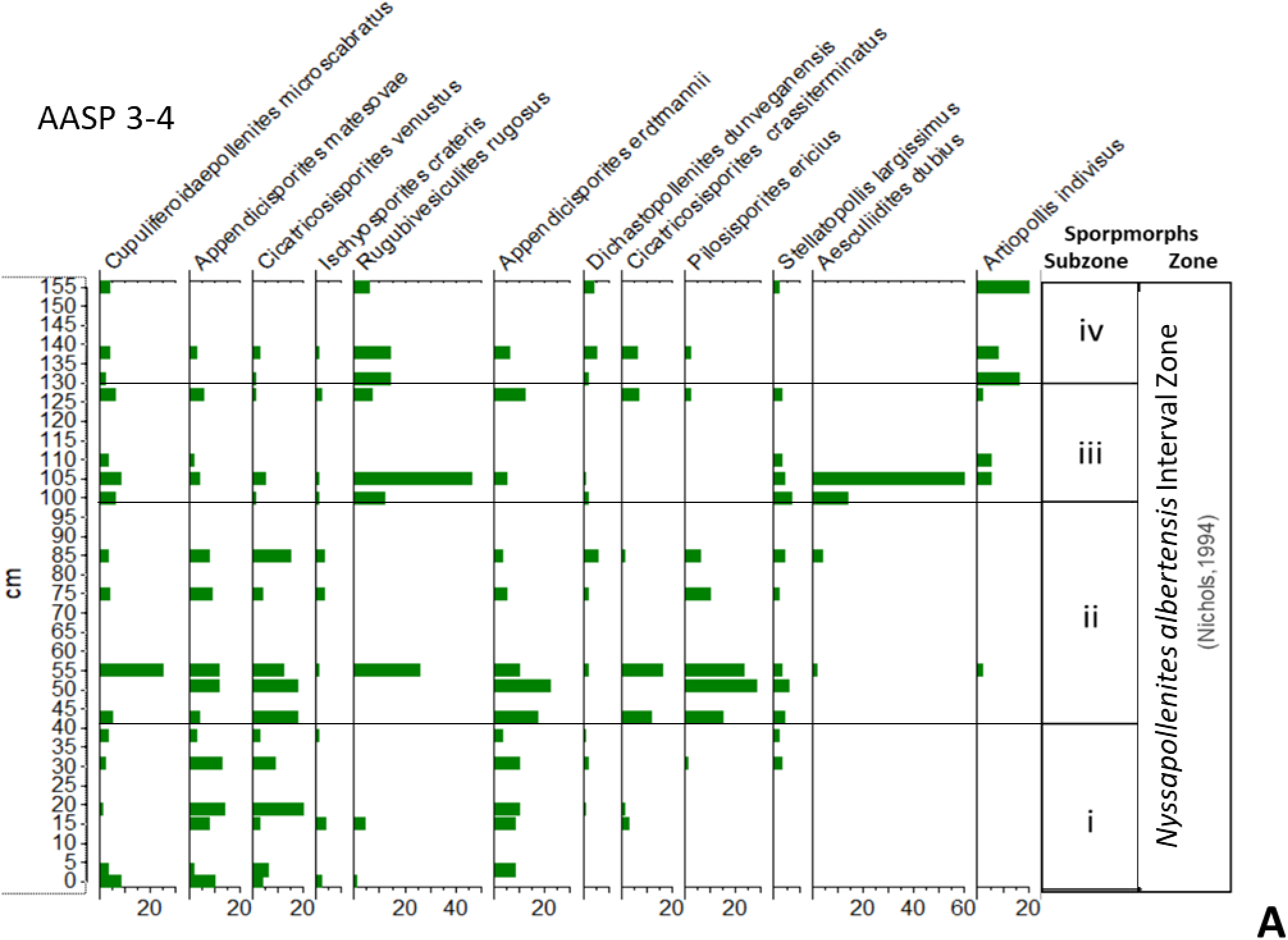
Presence–absence diagram showing the main bioevents observed that are the apparent first and last occurrences of key sporomorphs in AASP 3–4.

Other stratigraphically relevant species present in the Arlington Archosaur section and seldomly reported from other CWIS sites are:

- *Aquilapollenites psilatus* is a relatively common species in the Arlington Archosaur Site but not frequently reported in other CWIS sites. It was previously reported by Akyuz et al. (2016) from the Turonian Ferron Sandstone Member in Utah. However, there is some controversy about this morphospecies’ phylogeny. After careful consideration, we followed the long-established perspective that this is pollen with a triprojectate morphology, similar to *Phrygilanthus acutifolius* with possible affinity to extant Loranthaceae (Jarzen, 1977).

Worldwide, this species has been reported from the undifferentiated Late Cretaceous. Still, in the CWIS, it seems to occur only in sections from the Turonian or younger, which is quite relevant to our study from an age point of view.

- *Triplanosporites sinuosus*, reported from the Cenomanian–Coniancian (Alabama) but also from younger sections from the Campanian to the Maastrichtian (Montana) or even the Late Cretaceous–Paleocene (Montana). It is frequent and often abundant in samples from the Arlington Archosaur Site.

- *Plicatella fucosa*, reported from Iowa, the Dakotas, Nebraska, and Wyoming, Late Albian to Cenomanian. It is often abundant in samples from the Arlington Archosaur Site, but we did not separate this morphotype from *A. erdmanii*.

- Apparent first regular occurrence and acme of angiosperms (dicotyledonous) from the middle part of Facies B and upwards.

Based on the Arlington Archosaur Site samples’ association, we identified the *Nyssapollenites albertensis* Interval Zone (Nichols, 1994) of the Late Cenomanian to mid- Coniacian age.

Although the zonal marker *Nyssapollenites albertensis\** has not been seen in the AAS outcrop, we have a small tricolporate species associated with the zone, *Aesculiidites dubius*. There are also other species known to be associated with that zone throughout the section, including various *Cicatricosisporites* spp., *Classopollis* spp., *Cupuliferoidaepollenites microscabratus*, *Deltoidospora minor*, *Gleicheniidites senonicus*, *Laevigatosporites* sp. Also relevant is the absence of *Proteacidites*, a triporate angiosperm pollen. The section has the continuous presence of triprojectate pollen, e.g., *Aquilapollenites psilatus* and *Triplanosporitis sinuosus*, resembling some morphotypes from the *Aquilapollenites* palynofloral province. There is also occasionally present *Neoraistrickia* cf. *robusta* in the assemblage, a species with a known last apparent occurrence in the Cenomanian of the United States.

We propose that this zone is probably equivalent to Unit 4 described from the Dakota Formation (Rawn & Witzke, 1995, in Ludvigson et al., 2010), in which *Cicatricosisporites crassiterminatus* has its first appearance and angiosperms show a marked increase in diversity.

We also recognized four informal subzones within the *Nyssapollenites albertensis* Interval Zone in the AAS outcrop, which roughly coincide with the four stratigraphically constrain cluster analysis of the AASP 3-4 section. The subzone’s definitions are the following:

##### Subzone i

(Figure 33)

**Figure 33.**
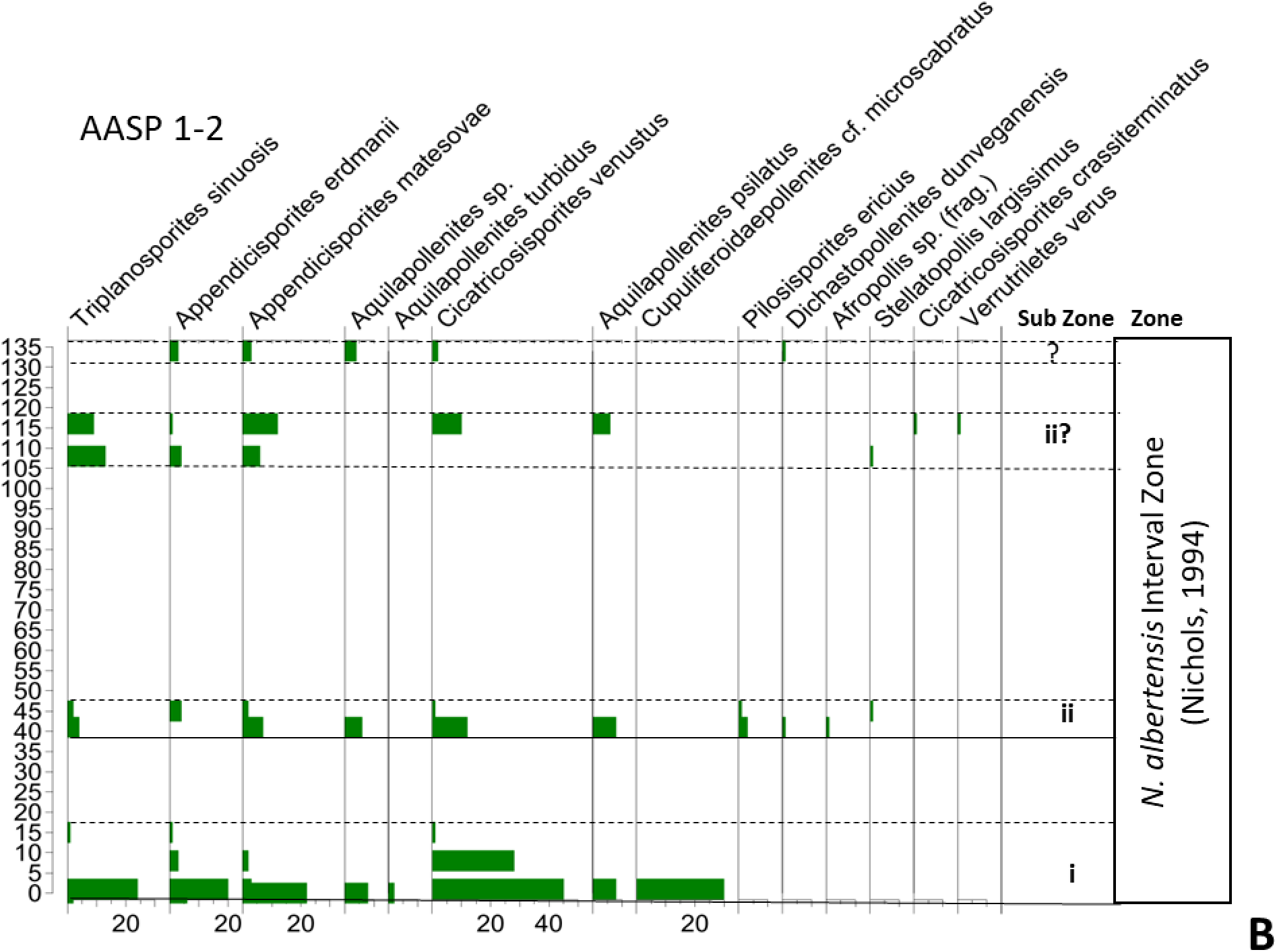
Histograms showing the abundance of the key spores and pollen species associated with the informal zonation proposed here for the AAS. The histograms are for sections (A) AASP 3– 4 and (B) AASP 1–2.

Characterized by the continuous presence of *A. matesoave*, *C. venustus*, and *A. erdtmanii*. Species absent from this subzone are *A. dubius* and *A. indivisus*. *P. ericius* is present but very rare in this subzone.

##### Subzone ii

(Figure 33)

The first apparent occurrence of *P. ericius* and *C. crassiterminatus* marks the base of the subzone. There is a significant change in the abundance of the assemblage towards the upper part of the subzone.

##### Subzone iii

(Figure 33)

The base of the subzone is characterized by the first apparent occurrence of *A. dubius*, and close to the base, *A. indivisus* also has its first regular occurrence. An acme of *R. rugosus* occurs close to the base of the subzone.

##### Subzone iv

(Figure 33)

It is characterized by the presence of *A. indivisus* and the absence of *A. dubius*. Other species absent or present with a single occurrence are *S. lagissimus* and *I. crateris*.

Figure 33 shows the informal sub-zonal subdivision defined for sections AASP 3–4, AASP 1–2, and the formal zonation (Nichols, 1994).

#### Dinoflagellates’ Palynostratigraphy

In a retrospective of the dinoflagellate cyst zonation evolution relevant to this paper, Williams (1977) did the first attempt to synthesize the previously existing dinoflagellate zonations; this effort continued in Williams and Bujak’s (1985) paper that summarized the ranges of hundreds of dinoflagellate cysts worldwide. Jarvis et al. (1988) published a detailed calibration for the Cenomanian–Turonian OAE2 using macro- and microfossils, including foraminifera and ostracods, calcareous nanoplankton, and dinoflagellate cysts, with δ^13^C and δ^18^O isotopes chemostratigraphy for sections in Dover, England. Williams et al. (2004) compared the Southern Ocean and the updated global dinoflagellate cyst events from the Late Cretaceous onward. They presented absolute ages for the first and last occurrences of some key species used in this paper.

Nøhr-Hansen (2012) presented a detailed identification of Cenomanian–Turonian dinoflagellate cysts bioevents in Greenland. This work is relevant because it identifies the FO of cold boreal waters’ Ccm morphological plexus, a key dinoflagellate cyst in the AAS section. We will further discuss its implications in the Age part of this paper.

Many dinoflagellate cyst taxa in the AAS section have stratigraphical ranges broader than Cenomanian and Turonian in the Northern Hemisphere, but some taxa seem more restricted in the CWIS. Although, in general, most Northern Hemisphere dinoflagellate zonation can be applied to the CWIS, in this work, we follow the CWIS dinoflagellate zonation proposed by Dodsworth and Eldrett (2019), as it is the last published for the CWIS with an updated age calibration. According to those authors, the ages assigned to their palynological zonation are based on an astronomical (obliquity) age model applied to the bioevents in Portland-1 (Eldrett et al., 2017) and Iona-1 (Eldrett et al., 2015a; Figure 7).

For the Arlington Archosaur Site, we identified the *Cauveridinium membraniphorum* Zone (Dodsworth & Eldrett, 2019) based on the presence of the Ccm morphological plexus throughout the sections and the absence of *Litosphaeridium siphoniphorum*.

Another relevant marker for this section is *Kiokansium polypes.* Its top (last) occurrence is restricted to the Late Cenomanian/Early Turonian in Utah, New Mexico, Colorado, and Arizona (Li, 1996).

According to Dodsworth and Eldrett (2019), the *Cauveridinium membraniphorum* Interval Zone is defined as follows: “From the first sample stratigraphically above the LO of common *Litosphaeridium siphoniphorum* (base of the zone) to the higher of either the LO of *Cauveridinium membraniphorum* or the LO of *Isabelidinium magnum* (top of the subzone)”.

Dodsworth and Eldrett (2019) proposed the *C. membraniphorum* Zone subdivision in the Western Interior Sea in four subzones: *Adnatosphaeridium tutulosum*, *Isabelidinium magnum*, *Senoniasphaera turonica*, and *Senoniasphaera rotundata*.

We propose that the AAS section probably belongs to the subzone *Isabelidinium magnum*, but we do not have the marker species *Adnatosphaeridium tutulosum* that characterizes the basal subzone of the *Cauveridinium membraniphorum* Interval Zone. Dodsworth and Eldrett (2019) defined the subzone of *Isabelidinium magnum* just above the *Adnatosphaeridium tutulosum* subzone as follows: “From the first sample stratigraphically above the LO of consistent *Adnatosphaeridium tutulosum* (base of the subzone) to the first sample stratigraphically below the FO of common *Senoniasphaera turonica* (top of the subzone).”

Following that definition of the subzone, we are in the interval in which *Adnatosphaeridium tutulosum* and *Senoniasphaera turonica* are absent. Still, other markers of the *Cauveridinium membraniphorum* Interval Zone are present.

Furthermore, based on the stratigraphically constrained cluster analysis, we subdivided the AAS section into three informal dinoflagellate cysts subzones:

##### Subzone I

(Figure 34)

**Figure 34.**
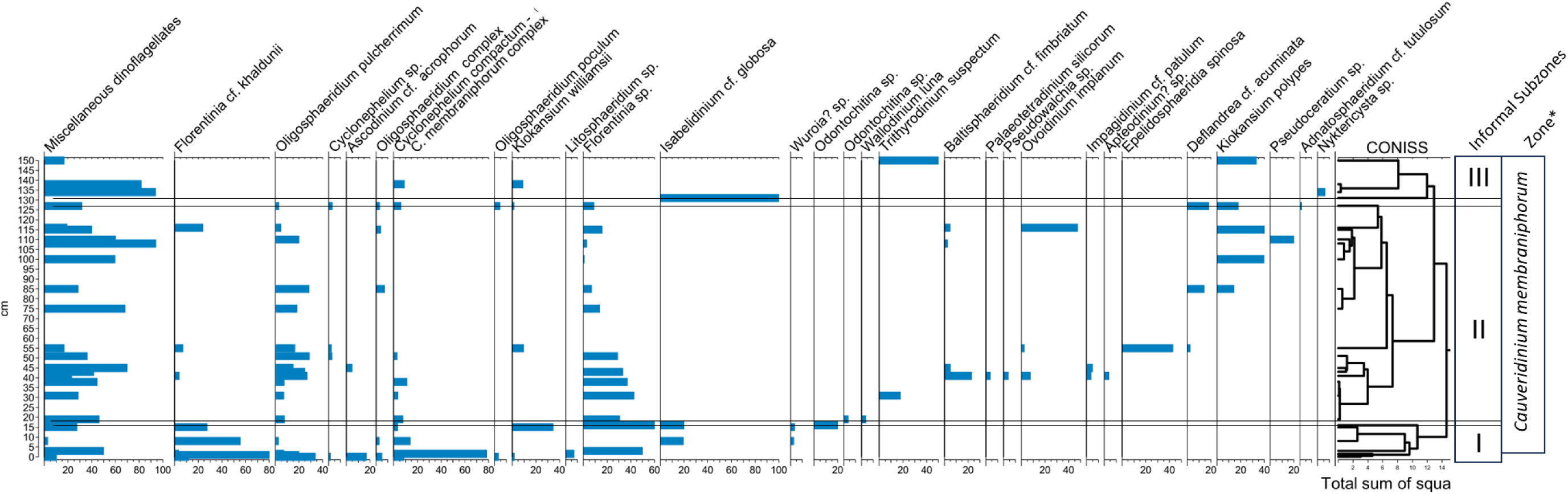
Dinoflagellate cysts distribution chart with abundances in absolute counts, combined Section 1–4, showing the constrained cluster analysis, total sum of squares (CONISS, Tilia). Based on the result of the cluster, three informal dinoflagellate cysts subzones were defined. Formal palynological zonation is based on Dodsworth and Eldrett’s (2019) zonal scheme.

This subzone roughly coincides with Facies A and is characterized by a lower diversity of the assemblages, with variations in the dominant species. *O. pulcherrimum* is dominant in the lower part of the subzone, followed by an increase in *Kiokansium williamsii.* The Ccm morphological plexus is present through this subzone. The diversity and abundance of dinoflagellate cysts in these assemblages decrease toward the top of the interval.

##### Subzone II

(Figure 34)

It is characterized by the presence of *Deflandrea* cf*. acuminata*. At the base are isolated occurrences of *Ovoidinium implanun* and *Epelidosphaeridia* cf. *spinosa*. The FAO and acme of *Kiokansium polypes* occur toward the middle part of the subzone (Figure 34).

##### Subzone III

(Figure 34)

This subzone is characterized by very poor dinoflagellate cysts assemblages, with the occasional occurrence of *Kiokansium polypes. K. williamsi* and the Ccm morphological plexus. Most species present in Subzones I and II are absent from this interval.

### CHRONOSTRATIGRAPHY AND AGE

According to Denne et al. (2016), the Woodbine Group’s age control is ammonites and inoceramids in marine interbedded shales and in the Pepper Shale found by earlier authors. Denne et al. (2016) also found foraminifera and calcareous nannofossils in the Pepper Shale. But at the Arlington Archosaur Site, the local paleoenvironment is unfavorable for the presence of the ammonites generally used for dating the Cenomanian. Since not a single specimen of those markers has been found at the site, its stratigraphic position is complicated to elucidate based only on lithology, because during the late Middle and Late Cenomanian, at least two delta complexes developed nearby, the Harris Delta and the Templeton Delta (Denne et al., 2016). So, lithological similitude is not a strong argument to support the age determination of the AAS section.

For that reason, it is palynology and the microvertebrate and vertebrate fossil faunas that may help to age-date the site and solve its stratigraphic position within the regional chronostratigraphic framework. A summary of the known biochrons in the CWIS of the key sporomorph species found in the AAS is in Figure 30, and the key dinoflagellate cysts in Figure 35.

**Figure 35.**
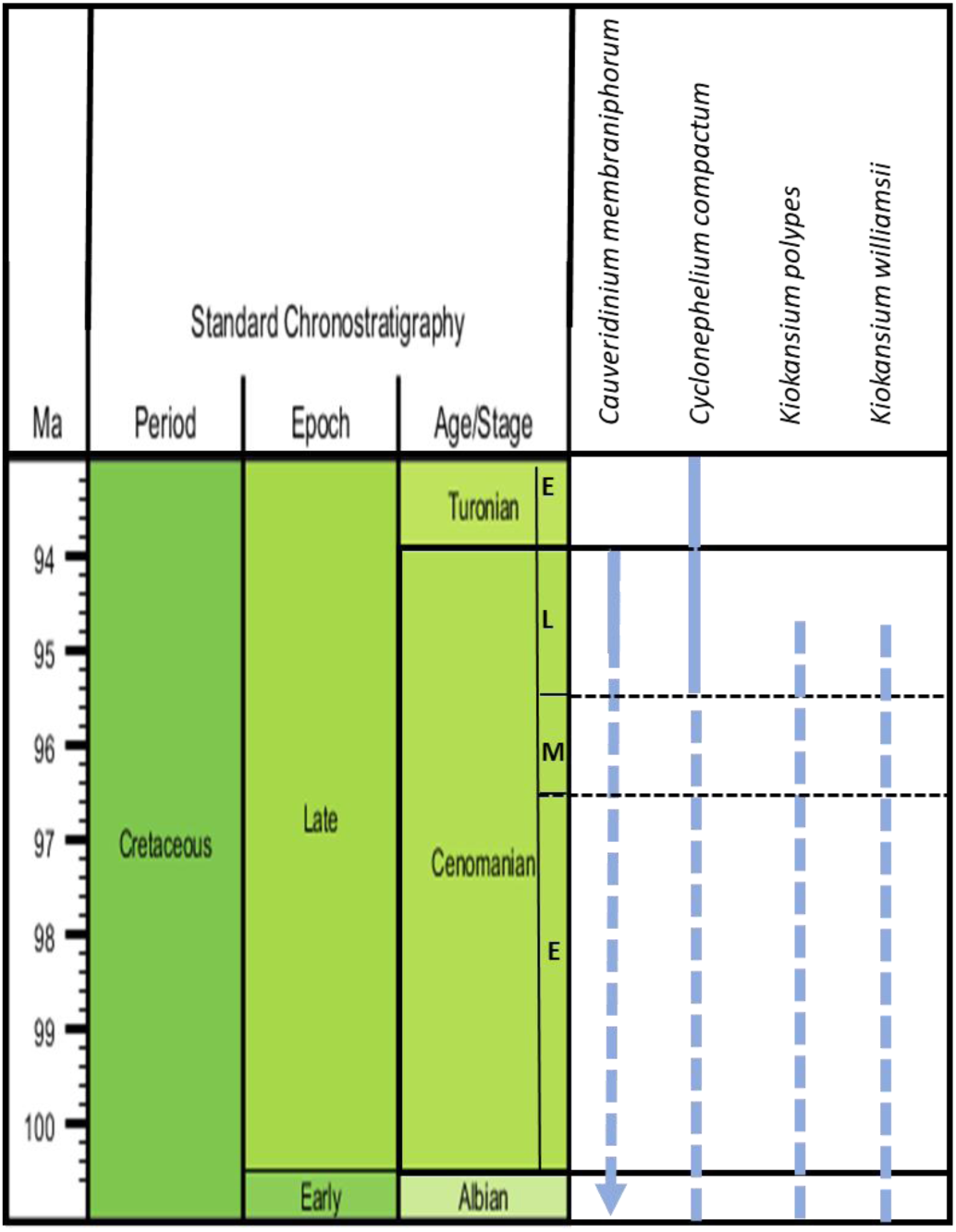
Known ranges for key dinoflagellate cysts as they have been reported from the CWIS to date (Dodsworth & Eldredt, 2019; Eldrett et al., 2014, 2015a, 2017; Harris & Tocher, 2003; PALYNODATA, 2006, updated to 2020 by author). General ranges taken from Dinoflaj3 (http://dinoflaj.smu.ca/dinoflaj3/index.php/Main_Page).

According to Nøhr-Hansen (2012), the FO of *Cauveridinium membraniphorum* indicates a Late Cenomanian age. It is also known that *Cauveridinium membraniphorum* has its first common occurrence in the upper Cenomanian onshore in the United Kingdom (Dodsworth, 2000; Pearce et al., 2009).

Dodswoth and Eldrett’s (2019) findings in other CWIS areas that are relevant to the AAS section are the consistent FO of *C. membraniphorum* at higher stratigraphic levels (i.e., intra-Upper Cenomanian) in mid-latitude sites, such as Pueblo, CO. Also, other locations within the central CWIS (Harris & Tocher, 2003) and Texas (Eldrett et al., 2014, 2015a, 2017) may have evidence of a southerly incursion of a boreal water mass during the Plenus Cold Event (PCE).

The last occurrence of the dinoflagellate species *K. polypes* and *K. williamsi* are restricted to the earliest part of the Late Cenomanian, according to Dodsworth and Eldredt (2019), when referring to the FO of *Alterbdinium* “*daveyi*” (i.e., the comparable form *Deflandrea* cf*. acuminata*) and the LO (last or top occurrence) of *Kiokansium williamsii*.

The LO of *Kiokansum williamsii* and *Deflandrea* sp. (*D*. cf. *acuminate*) was documented by Eldrett et al. (2017) in the Portland-1 well, where both bioevents occur just above the base of the Upper Cenomanian. Dodsworth and Eldrett (2019) suggested an age of ca. 95.6 Ma for those events. Figure 35 summarizes the known rages of several key dinoflagellate cyst species in the AAS section.

According to Ogg and Hinnov (2012, in Gradstein et al., 2012, p. 802), when discussing the position of the boundaries of the Cenomanian substages, specifically between the Middle and Upper Cenomanian, “the replacement of *Acanthoceras* ammonites by the *Calycoceras* genus is commonly used to mark the base of the Upper Cenomanian (Hancock, 1991).” However, the marker for the base of the Upper Cenomanian has not yet been selected. So, not even with ammonites or any other marine fossil group has the official boundary been defined. Those authors commented, “probably placement would be near the *Dunveganoceras pondi* Zone of the Western Interior, the base of which Cobban et al. (2006) used for their Upper/Middle substage boundary.”

From another point of view, the Cenomanian falls within Chron 34. It is important to remember that the entire Cenomanian has a duration of 6.6 Ma from 100.5 Ma to 93.9 Ma.

At the studied AAS, there is an apparent top occurrence of *K. williamsii* and the presence along the section of the Ccm morphological plexus and *K. polypes*. Since *Cyclonephelium membraniphorum* has its FO in the CWIS intra-Upper Cenomanian, then the AAS section should not be older than that age. Also, taking into consideration that the long-ranging *K. williamsii* and *K. polypus* have their LO during the early Late Cenomanian in the CWIS, as was discussed before, an (early) Late Cenomanian age for the AAS seems consistent with the observations about the age of other sections of the CWIS.

The integration of the palynological (pollen, spores, and dinoflagellate cysts) evidence in this study supports that the age of the AAS section is most probably (early) Late Cenomanian, in the sense of Gradstein et al. (2012).

### STRATIGRAPHIC RELATIONSHIPS

From a paleogeographic point of view, the AAS section would have been at the southernmost extent of the CWIS. During the late Middle and Late Cenomanian, several delta complexes developed in paleogeographical proximity to the AAS at different times: the Woodbine Delta, the Harris Delta, and the Templeton Delta (Hetz et al., 2014; Denne et al., 2016; Gifford, 2021).

We ruled out the possibility that the AAS section belongs to the Arlington Member because that lithostratigraphic unit has been described as “fine-grained sandstones with specimens of *Ostrea* and *Exogyra* (oysters). Powell (1968) reported the presence of the ammonite *Acanthoceras wintoni* (a junior synonym of *Conlinoceras tarrantense*),” according to Denne et al. (2016, p. 41). Due to the shallow marine to coastal paleoenvironments that characterized the AAS section, none of those fossils are present, but on the other hand, there are abundant palynomorphs, vertebrate remains, mollusks, many other types of macrofossils as well as some coalified plant material, and palynomorphs.

In spite of the East Texas Basin being explored for oil since the early 1900s and the existence of well logs, cores, and seismic data, besides the field geology work done, the relationships between the Eagle Ford and Woodbine groups is still unclear due to the complexity of the stratigraphic architecture of these units, recently illuminated by the seismic stratigraphic and log analysis work by Gifford (2021) that found the Harris Delta primarily deposited in unfilled accommodation space left by the prograding Woodbine Delta. But the correlation between the multiple outcrops and the subsurface is still unclear.

The consensus is that the Eagle Ford overlies the Woodbine strata unconformably, but according to Gifford (2021), “In sharp contrast within the southern ETB, Late Cenomanian sandstones as well as underlying Middle Cenomanian organic-rich mudstones, are typically included within the Woodbine Group (Hentz et al., 2014).” Hence, as previously discussed, complications and inconsistencies arise due to assumed equivalence between lithostratigraphic and chronostratigraphic units, especially in this case’s complex stratigraphic architecture setting.

A further complication is the similarity in the lithologies deposited at different ages within the Woodbine Delta and the Harris Delta. Although the sedimentation of the Harris Delta system has been recognized as a younger Late Cenomanian event, many authors took a lithostratigraphic approach that, at times, resulted in the inclusion of the Harris Delta Late Cenomanian sands and underlying Middle Cenomanian organic-rich rocks as part of the Woodbine Group; as a consequence, the Woodbine became partially equivalent to the Lower Eagle Ford Formation of South Texas (Gifford, 2021; Hentz et al., 2014). From a historical point of view, the “Eagle Ford Formation” was named the Tarrant, Britton, and Arcadia Park formations in the outcrop belt in the Dallas area. In more recent revisions of the stratigraphy of the Cenomanian–Turonian of Texas, several authors (e.g., Breyer et al., 2013; Denne & Breyer, 2016; Denne et al., 2016; Donovan et al., 2015, 2019; Gifford, 2021; Hentz et al., 2014), using sequence stratigraphic approaches, applied the Eagle Ford nomenclature of South Texas (Lower and Upper Eagle Ford Formations) to the subsurface and outcrops of the East Texas Basin.

Modern papers also use seismic terminology to key surfaces, like sequence boundaries, to separate the main units, e.g., K600sb, K630sb, and K720sb (Gifford, 2021). Also, Denne and Breyer (2016) and Fenne et al. (2016) named the regional Cenomanian–Turonian depositional episodes in the subsurface EGFD100, EGFD200, EGFD300, EGFD400, EGFD500, and EGFD600.

According to Gifford (2021), “The Early Cenomanian Freestone (Woodbine) Delta prograded into the basin, depositing thick successions of marginal-marine and fluvial sandstone, as well as offshore mudstone. Woodbine progradation ceased, and Middle Cenomanian organic-rich Eagle Ford mudstone was deposited over distal Woodbine mudstone, sequentially onlapping as accommodation space was filled. … Because of the facies similarities between the K600 (Freestone) and the K645 (Harris) Deltas, a regional perspective based on surface (sequence boundary) mapping is necessary to distinguish them from each other.” How these episodes can be recognized on outcrops is not straightforward and requires much biostratigraphic work.

From the above-explained complex lithostratigraphy, palynology studies can help to age- date and solve the outcrop belt’s chronostratigraphic puzzle. Based on the results of this study, we propose answers to the following stratigraphic questions:

To which paleodelta (Harris or Templeton) does the AAS belong? To neither of them—the AAS outcrop seems to be an (early) Late Cenomanian unnamed lateral equivalent of the lower Harris delta, whose deposition was confined to a more southeasterly position (Gifford, 2021, Figure 15) in the subsurface of the Leon, Madison, Brazos, Grimes, and Walker counties).

To which lithostratigraphic unit does the AAS belong: Woodbine, Lewisville, Britton, or the lowermost Eagle Ford? Depending on the lithostratigraphic scheme used, it would belong to the Woodbine Group (Hentz et al., 2014) or the Eagle Ford Group in Gifford (2021, Figure 10). If an age approach is taken, the AAS outcrop would be related to one of the sandier, less organic-rich intervals of the Eagle Ford stratigraphic “event.”

What is the age of the rocky outcropping at the AAS, Middle or early Late Cenomanian? It is (early) Late Cenomanian, at the onset of the PCE of the OAE2..

How does it correlate with other key sites in the CWIS? It correlates with part of the Harris delta recognized in the subsurface of Northeast Texas and with the sections that contain the PCE (OAE 2), which was identified in the Pueblo and other Cenomanian–Turonian sections along the east and west paleo-shores of the CWIS.

A tentative correlation between AAS and other outcropping sections studied with palynology by Cloos (2018) on the Outcrop Belt is shown in Figure 36. The correlation is based on two key palynological events: the Ccm morphological plexus, which could be present in the Acme Brick section where Cloos identify *Cyclonephelium* sp., and the arrival of the angiosperm *Normapolles* pollen types in the area, which are also recorded in the same section but are absent from the AAS outcrop.

**Figure 36.**
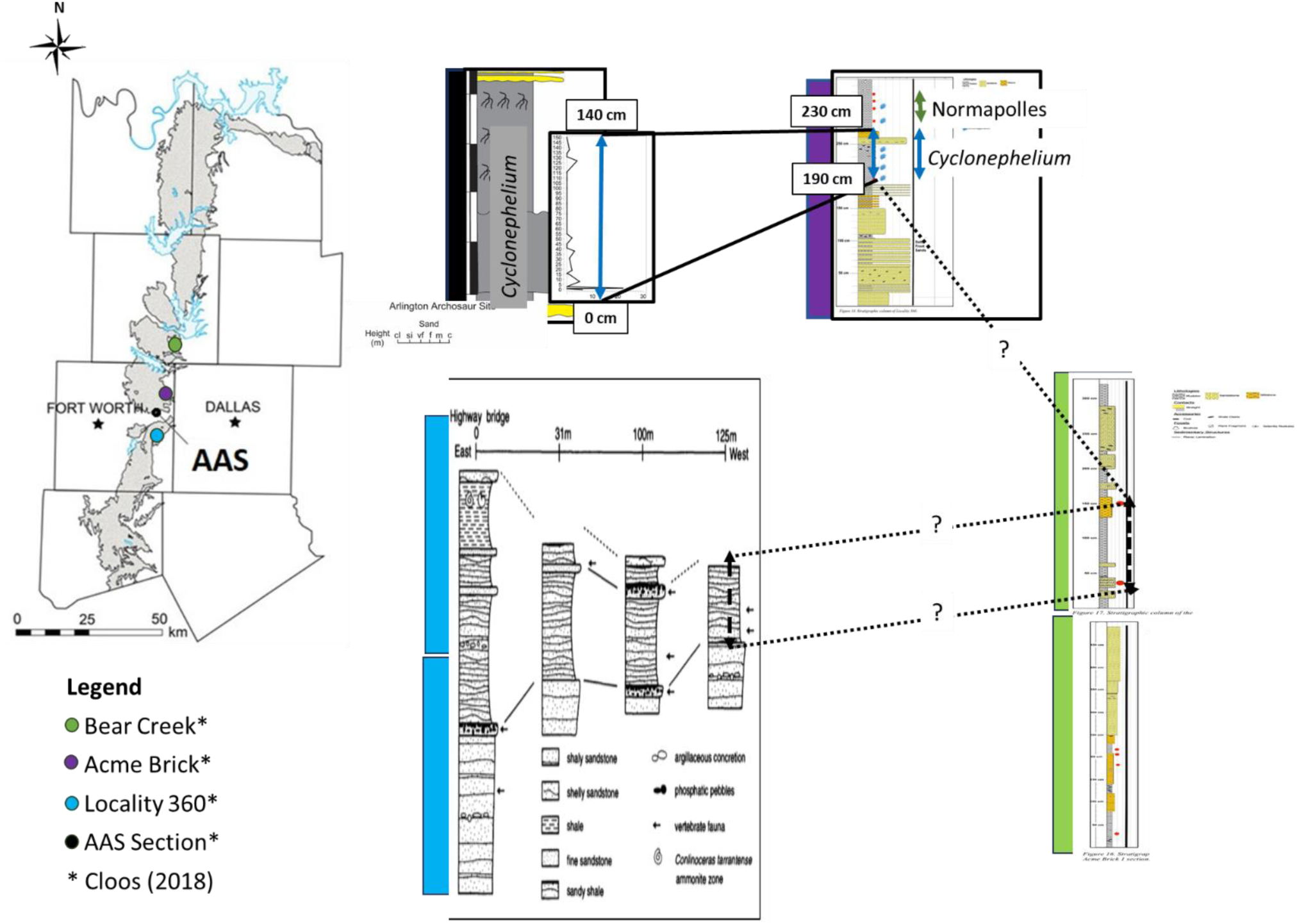
Proposed palynological correlation between the outcrop belt sections near Fort Worth, based on the occurrence of the *Cyclonephelium* group and the first appearance of *Normapolles* sp.

This tentative palynological correlation shows the complexity of the chronostratigraphic relationship between the North Texas (Dallas) outcrop belt sections. It shows that even over short distances, there are biostratigraphy detectable gaps related to sedimentation interruptions and lateral facies changes during the Middle and Late Cenomanian on the southernmost shore of the CWIS.

## CONCLUDING REMARKS

A complex stratigraphic architecture characterizes the Middle and Late Cenomanian of North Texas, as shown in the most recent sequence stratigraphic works discussed in this paper. The area of the AAS sedimentation was located in the interface between the coastal sedimentation and the marine settings that characterize other areas of Texas’s CWIS. From a lithostratigraphic point of view, the lithofacies changes due to the variation in the conditions of the coastal settings under the influence of high-order stratigraphic cycles impacted the sedimentation in these settings, creating clastic wedges that generally thinned laterally and basinward, depending on the changes of the paleo-coast over time. For those reasons, a sequence stratigraphic-chronostratigraphic approach based on robust biostratigraphic data is key to illuminating the resulting lithological puzzle.

This work does not aim to change the age of widely used lithostratigraphic units but to place the sedimentation of the rocks outcropping at the Arlington Archosaur Site in time and space based on new high-resolution palynological data. Based on our results, we propose that from a paleogeographic point of view, the AAS outcrop was part of the sedimentation of an unnamed easterly lateral equivalent of the lower Harris delta, which sedimentation took place during the onset of the PCE of the OAE2 in the (early) Late Cenomanian. The findings of the Ccm morphological plexus, supported by the other characteristics of the palynological assemblages in the AAS, show that this section probably represents the southernmost reach of the PCE (OAE 2) incursion of boreal waters into the CWIS, an event registered in other sections in the basin and elsewhere in the world.

## Plate I

1. Appendicisporites erdmanii Pocock, 1964/Plicatella fucosa (Vavrdova) Davies 1985. Section AASP 3-3
2. Stellatopollis largissimus Singh, 1983. Sample AASP 3-4
3. Cicatricosisporites venustus DeЎk, 1963. Sample AASP 4-1
4. Pilosisporites ericius Delcourt, 1955. Sample AASP 4-3
5. Ischyosporiles (Cicatricosisporites?) crateris Balme, 1963. Sample AASP 3-9
6. Dichastopollenites dunveganensis Singh, 1983. a) Sample AASP 3-5; b) Sample AAS 3-8
7. Cupuliferoidaepollenites microscabratus Jarzen and Dilcher, 2008. Sample AASP 3-10
8. Cluster of Aesculiidites dubius (Jardine and Magloire) Tschudy B. 1973. Sample AASP 3-9
9. Zlivisporis cenomanianus (Agasie) Braman, 2010. a) Sample AASP 4-3; b); Sample AASP 3-10

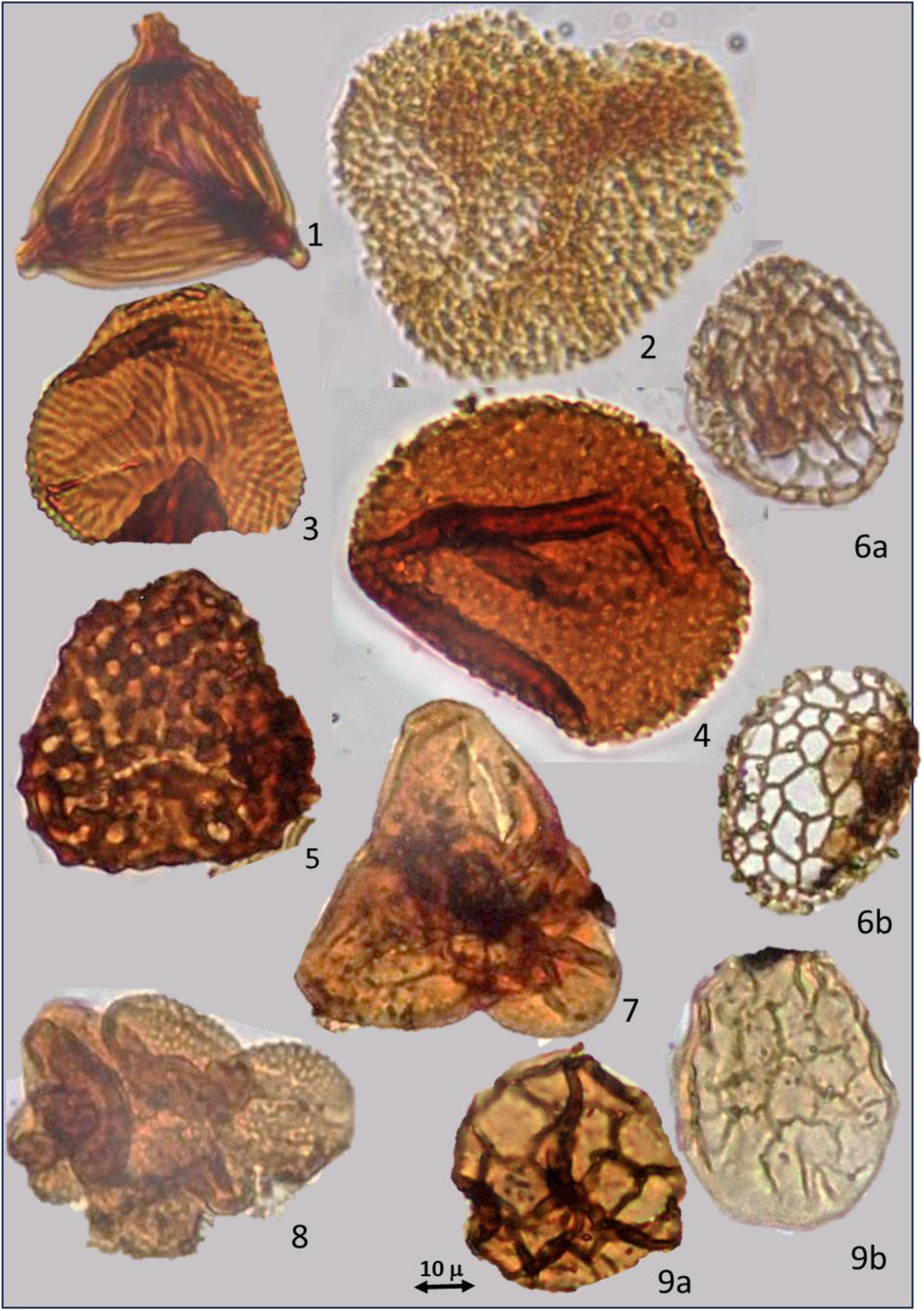

## Plate II

1. Kiokansium williamsii Singh 1983. Sample AAS 3-11.
2. Kiokansium polypes. Sample AAS 3-8.
3. Cauveridinium membraniphorum – Cyclonephelium compactum (Ccm morphological plexus). a) C. membraniphorum end-member, Sample AAS 2-1; b) C. compactum, Sample AAS 3-11.
4. Oligosphaeridium pulcherrimum. Sample AAS 3-11.
5. Micrhystridium sp. Sample AAS 3-11
6. Cyclonephelium vannophorum. Sample AAS3-2.

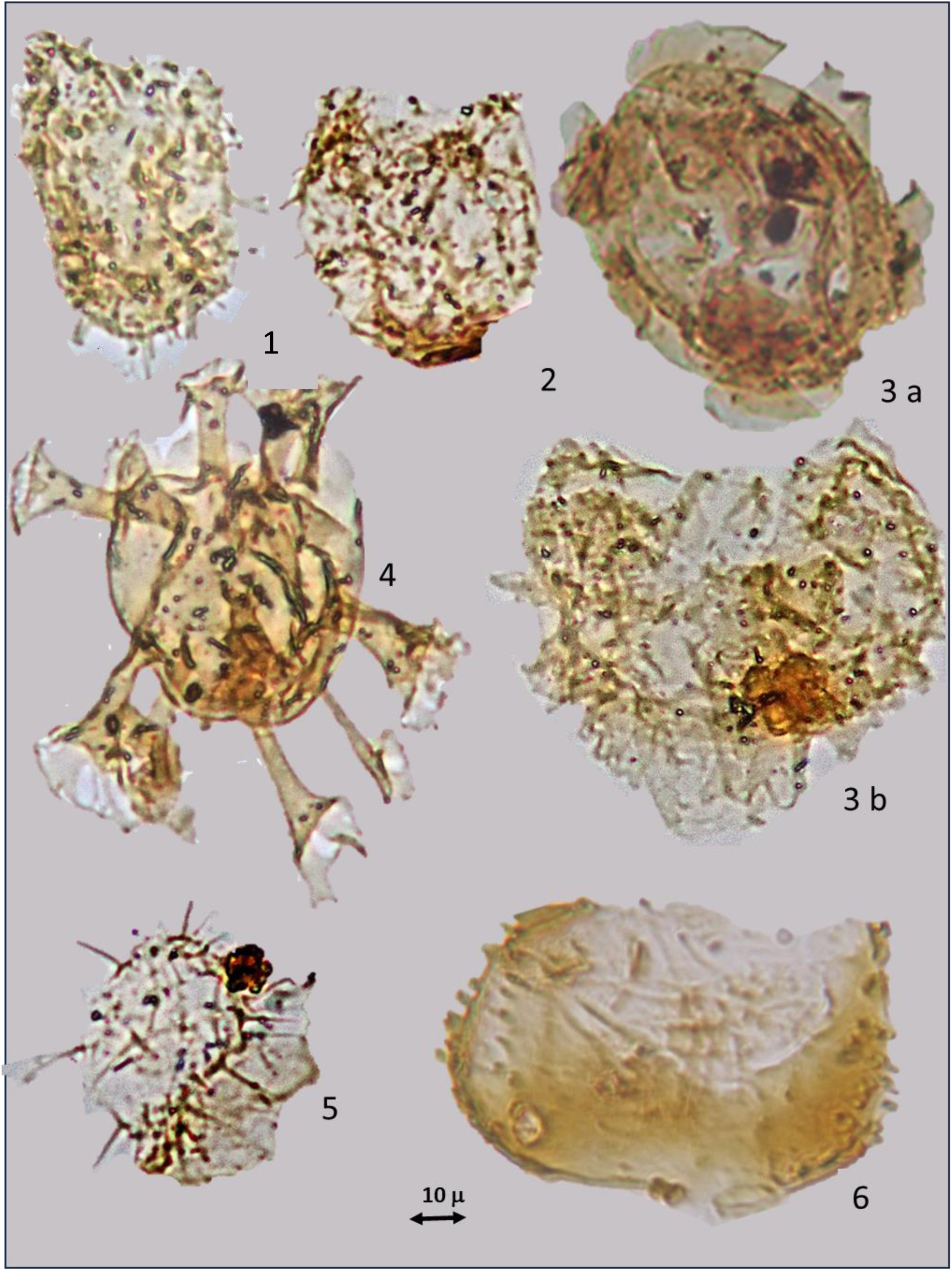

